# Octopamine receptors at a glance: from expression and anatomical maps to their role in development and behavior in the *Drosophila melanogaster* larva

**DOI:** 10.64898/2026.05.05.722892

**Authors:** Alexandra Großjohann, Vincent Richter, Franziska Reinhardt, Marvin Hah-mann, Ronja Badelt, Juliane Kinnigkeit, Jana Breitfeld, Peter Kovacs, Peter F. Stadler, Irene Coin, Andreas S. Thum

## Abstract

Octopamine is involved in a variety of different physiological and behavioral mechanisms in *Drosophila melanogaster*. Throughout the life cycle of the fruit fly, from the larva to the adult, octopaminergic neurons in both the central and the peripheral nervous system target a multitude of neurons and even non-neuronal tissues, making it challenging to analyze individual mechanisms of octopamine function. One approach to deconstructing this complex system is to examine the postsynaptic components of signal transmission. In *Drosophila*, octopamine interacts with six distinct G-protein-coupled receptors. For some of these receptors, expression maps and functional implications have been described. In contrast, other receptors have been neglected, partly due to the lack of suitable genetic tools. Here, for the first time, we compiled a complete set of mutant lines of all known octopamine receptors, all generated using the same genetic tool, the recently established Trojan Exon system. It integrates the Gal4/UAS binary expression strategy while simultaneously impairing receptor function. This enabled us to generate a comprehensive anatomical map of receptor expression in the larva and, at the same time, analyze the function of individual octopamine receptors during larval development, chemosensory perception and locomotion. All octopamine receptors (Oamb, Octα2R, Octβ1R, Octβ2R, Octβ3R, and Oct-TyrR) showed extensive signal in the central nervous system. The same was found for the peripheral nervous system, with the exception of Octβ2R, which showed pronounced expression in the somatic muscles. We also observed a previously undescribed role of Octβ1R, Octβ3R, and Oct-TyrR in larval hatching and in the survival of larvae and pupae. Molecular evaluation of the Trojan Exon octopamine lines supports our analysis. In addition, we combined the experimental results with gene expression data from the different development stages of *Drosophila melanogaster* and from different tissues and cell populations throughout the body. Overall, we compiled, analyzed and validated a complete set of octopamine lines which, together with gene expression analysis, provides a basis for further functional studies on the larval octopaminergic system.

## Introduction

Octopamine (OA) is a biogenic amine that acts as a neuromodulator, neurotransmitter, and neurohormone in invertebrates, and plays an important role in regulating a variety of behavioral and physiological processes. Extensive research has been conducted on its functions in a range of species, including *Drosophila melanogaster*, *Apis mellifera* and *Caenorhabditis elegans*, as well as on its structural analog norepinephrine in vertebrates (reviewed in David and Coulon, 1985; Roeder, 2005; Farooqui, 2012; Roeder, 2020; Mezheritskiy et al., 2024).

In *Drosophila*, octopaminergic cells, that produce OA via Tyramine β-hydroxylase (Tβh), are found not only in the central and peripheral nervous system of both adults (Monastirioti, 2003; Zhou et al., 2008; Busch et al., 2009; Busch and Tanimoto, 2010; Schneider et al., 2012; Pauls et al., 2018; Babski et al., 2024; Selcho, 2024) and larvae (Koon et al., 2011; Selcho et al., 2012), but also in the adult female reproductive sys-tem (Monastirioti et al., 1996; Monastirioti, 2003; Rezaval et al., 2014). OA is versatile in its function being involved in memory formation (Schwaerzel et al., 2003; Yarali and Gerber, 2010; Burke et al., 2012; Huetteroth et al., 2015; Berger et al., 2024), decision-making (Classen and Scholz, 2018), sleep (Crocker and Sehgal, 2008; Crocker et al., 2010; Erion et al., 2012; Li et al., 2016; Deng et al., 2019 but see also Reyes et al., 2026), arousal (Kula-Eversole et al., 2010), the circadian clock (Shang et al., 2011; Huang et al., 2013), gut-brain axis (Schretter et al., 2018), feeding (Li et al., 2016; Wang et al., 2016; Youn et al., 2018), starvation response (Yang et al., 2015; LeDue et al., 2016; Li et al., 2016; Yu et al., 2016; Damrau et al., 2018; Sayin et al., 2019; Pauls et al., 2021), heartbeat (Johnson et al., 1997), flight activity (Brembs et al., 2007; Suver et al., 2012; van Breugel et al., 2014; Manjila et al., 2019), climbing activity (Li et al., 2016; El-Kholy et al., 2022), movement endurance (Sujkowski et al., 2017; Sujkowski and Wessells, 2018), courtship behavior (Certel et al., 2007; Certel et al., 2010; Zhou et al., 2012; Andrews et al., 2014) and male aggression (Hoyer et al., 2008; Andrews et al., 2014). In larvae, OA plays a role in learning and memory (Selcho et al., 2014; Franke et al., 2026), sleep (Szuperak et al., 2018), chemotaxis and startle response (Ma et al., 2016), feeding (Branch et al., 2017), starvation response (Koon et al., 2011; Koon and Budnik, 2012; Zhang et al., 2013; Branch et al., 2017; Schützler et al., 2019), locomotion (Saraswati et al., 2004; Fox et al., 2006; Koon et al., 2011; Koon and Budnik, 2012; Selcho et al., 2012; Pauls et al., 2015) and nociceptive es-cape behavior (Li et al., 2023; Boivin et al., 2026). In contrast to behavior, the involvement of OA in early development ranging from the embryo to the pupa has only been scarcely investigated.

OA can bind to six distinct G-protein-coupled receptors (GPCRs) in *Drosophila*, which are classified into three groups: (1) α-adrenergic-like receptors, including Oamb and Octα2R; (2) β-adrenergic-like receptors, consisting of Octβ1R, Octβ2R, and Octβ3R; and (3) a tyramine (TA) type 1 receptor, known as Oct-TyrR (see Table 1). Each receptor class differs in its structure, with its classification derived from the vertebrate adrenergic receptors. Upon OA binding, different G-proteins interact with the receptors, inducing different downstream signals (see Table 1). Oamb activation leads to an increase in Ca^2+^ levels via the Gαq mediated phospholipase C pathway (Han et al., 1998; Balfanz et al., 2005). Interestingly, depending on the ligand concentration, the Ca^2+^ response can be a single event that slowly declines (high concentration) or can be in the form of oscillations (low concentration; Balfanz et al., 2005). A Ca^2+^/Calmod-ulin-dependent protein kinase II activity (CaMKII) in the oviduct epithelium has also been reported (Lee et al., 2009). The second α-adrenergic-like receptor, Octα2R, in-hibits the adenylyl cyclase pathway via Gαi/o, leading to a decrease in cyclic adenosine monophosphate (cAMP) level (Andrews et al., 2014). All three β-adrenergic-like receptors have been found to interact with the Gαs protein which leads to an activation of the adenylyl cyclase pathway and an increase of cAMP (Octβ1R: Maqueira et al., 2005; Balfanz et al., 2005; Octβ2R and Octβ3R: Maqueira et al., 2005). Yet for Octβ1R, conflicting evidence suggests that it may reduce cAMP levels by coupling to the inhibitory Gαi/o (Koon and Budnik, 2012). The Oct-Tyr receptor can be activated by two ligands, OA and TA, resulting in two possible downstream signaling cascades: either Gαq activity leads to a Ca^2+^ level increase (Robb et al., 1994) or Gαi/o leads to a cAMP level decrease (Saudou et al., 1990; Robb et al., 1994; Chatwin et al., 2003). OA is more potent to evoke the Gαq mediated downstream pathway, while TA is more potent to evoke the Gαi/o mediated downstream pathway. The broad diversity of OA receptors (OAR) and their different downstream signaling pathways highlight the intricate nature of OA’s involvement in regulating numerous physiological processes and behaviors. This complexity also complicates efforts to link individual receptor types to specific functions, making a comprehensive analysis of OARs challenging.

**Table 1:**
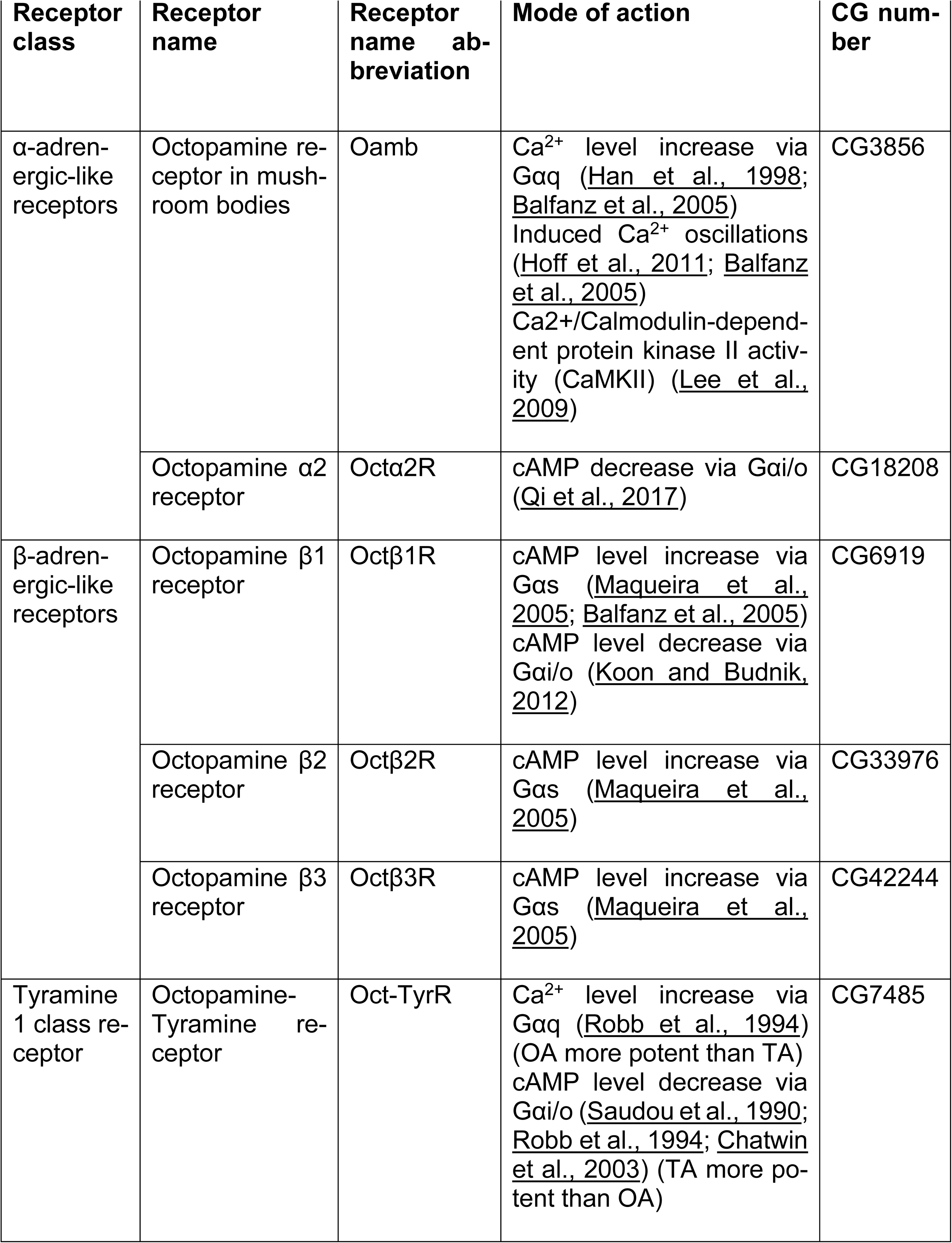
Overview of octopamine receptors. List of all octopamine receptors with receptor class, receptor name, abbreviation, mode of action (G-protein downstream signaling upon octopamine binding) and CG number of the gene. OA: octopamine; TA: tyramine.

Previous data of OA and OARs focused mostly on only OA, individual OARs, or incomplete sets of OARs. Here, we compiled a complete collection of mutant lines to apply the same assays for OA and each OAR. Not only is this the first complete set for all known OARs, but they were also all generated using the same genetic tool – the Trojan Exon developed by Diao et al. (2025). The Trojan Exon lines were generated from the MiMIC collection (*Minos-*mediated integration cassette; Venken et al., 2011) by inserting the Trojan Exon cassette at the MiMIC docking site via recombina-tion-mediated cassette exchange. The Trojan Exon is described as a “coding intron” since it is inserted into an intronic region and contains a splice acceptor and a splice donor, thereby masking it as an exon. Moreover, the cassette encodes a Gal4 for gene-specific targeting, which is translated separately from the endogenous protein. This integration event produces a truncated protein, the length of which is determined by the insertion site within the gene. Such truncation can compromise the functionality of the endogenous protein, while concurrently enabling precise visualization of its native expression pattern via a Gal4-dependent reporter.

With this set of Trojan Exon OA and OAR lines we observed broad labelling of each OAR in the central (CNS), and the peripheral nervous system (PNS), and Octβ2R specific signal in somatic muscles. Bulk RNA-sequencing analysis allowed us to identify further yet unknown expression locations, such as larval lymph gland, pupal brain, eye and salivary gland, and adult glia, olfactory projection neurons and neuronal subsets in the protocerebral bridge. For the first time, we generated an anatomical map of Octα2R in the larva. We observed large overlapping expression patterns of different OARs in various tissues and cell populations throughout the body, from larva to adult. Using the same genetic tools, we found that larval hatching ability and larval survival are impaired in *Octβ1R*, *Octβ3R* and *Oct-TyrR* mutants. Pupal mortality is increased in *Oct-TyrR* larvae. These results suggest that the involvement of OA during development is greater than previously thought. We compared our results with findings from previous studies and, together with a molecular evaluation of the Trojan Exon OA and OAR lines, provide a complete and validated set of mutants to analyze the OA system even beyond this study.

## Materials and Methods

### Fly strains

Fly strains were reared on standardized cornmeal medium at 25 °C, 65 % relative humidity and in a 14:10 h light:dark cycle. Trojan Exon octopamine and octopamine receptor mutant lines were obtained from the BDSC: Oamb (RRID:BDSC_67506), Octα2R (RRID:BDSC_67636), Octβ2R (RRID:BDSC_67511) (the first three lines carried the TM3, Sb[1] Ser[1] balancer with no visible marker in larvae; Oamb and Octβ2R were re-balanced over TM6B, Hu, Tb [1] using *w*;Bl/CyO;𝑇𝑀2/𝑇𝑀6𝐵*); moreover, Octβ1R (RRID:BDSC_76677), Octβ3R (RRID:BDSC_76680), Oct-TyrR (RRID:BDSC_77735) and Tβh (RRID:BDSC_76655). Control lines were CantonS (WT-CS) and yellow white (*y* w\**). Only homozygous individuals were used for experiments. Each experiment was performed at 25 °C.

### Oviposition

Freshly hatched or max. one day old female flies of each genotype were paired with WT-CS male flies in a 1:1 ratio to ensure a high copulation probability and maximize the yield of laid eggs. After a mating period of 72 h, individual female flies were placed on oviposition plates (Sarstedt, 82.1135.500, Ø 35 mm) filled with 2.5 % agarose (w/v) (VWR-Life Science, CAS 9012-36-6), 3 % sucrose (w/v) (Roth, CAS 57-50-1) and 30 % store bought apple juice (Sachsen Obst, 100 % fruit). The oviposition plates were topped with yeast paste. 24 h later, the number of laid eggs was counted.

### Viability

The oviposition assay was repeated with female and male flies of the same genotype and up to 4 female flies per oviposition plate. The increased number of females ensured to have a proper number of eggs and later larvae to assess larval and pupal mortality. Female flies were allowed to lay eggs for 24 h and after another 24 h the number of hatched larvae was counted. From these plates, 30 hatched larvae were then transferred to standard food vials and monitored closely. The number of pupae and emerging flies was quantified three times a day over a period of 16 days. Hatching rate describes the number of hatched larvae per total number of eggs. Larval mortality is the number of larvae that did not enter the pupal stage in percentage. Pupal mortality is the number of flies that did not hatch from the pupa in percentage.

### Locomotion assay

The protocol used was modified from Bhatt and Neckameyer, 2013. Third instar for-aging stage larvae were washed with tap water to remove residual food and transferred to a petri dish (Sarstedt, 82.1472, Ø 92 mm) filled with 2.5 % agarose (w/v). After 30 s of acclimatization, single larvae were filmed for 75 s at a rate of 20 fps with an area scan camera (Basler, 106909, lens: 25 mm) and the software Pylon viewer (v 6.3.0; Basler). For a total duration of 1 min each peristaltic wave was counted for 15 larvae per genotype.

### Feeding assay

Feeding behavior was assessed via the rate of mouth hook contractions of third instar foraging stage larvae on a petri dish filled with 2.5 % agarose (w/v) and topped with 5 ml of 2 % activated bakeŕs yeast solution. Yeast solution prevented larvae from moving around ensuring that the recorded mouth hook contractions were a result of feeding behavior. Before the experiment, larvae were washed with tap water to remove residual food. Larvae were allowed to adjust to the new surrounding for 30 s and afterwards were filmed for 75 s with the same setup and software as for the locomotion assay. Each mouth hook contraction over a period of 1 min was counted of 15 individual larvae per genotype.

### Olfactory and Gustatory Preference tests

Experiments were performed as described in Apostolopoulou et al., 2013. For the gus-tatory preference test, petri dishes were filled with a thin layer of pure 2.5 % agarose (w/v) on one half and 2.5 % agarose (w/v) mixed with the appetitive stimulus (fructose (Roth, 57-48-7; 2 M) or arabinose (Sigma Aldrich, 10323-20-3; 2 M)) on the other half. The middle of the petri dish was marked as a 10 mm wide stripe separating sugar and pure agarose side. In this area, 30 larvae were initially placed and were given 5 min of free motion before being counted. Three groups emerge: larvae on the sugar side (#sugar), larvae on the pure agarose side (#agarose) and larvae in the middle. The total amount of larvae resulted from the number of larvae in each group (#total). The preference index was calculated as follows: #sugar minus #agarose divided by #total. Olfactory preference tests were performed on a petri dish filled with 2.5 % agarose (w/v) and one empty custom-made Teflon container (4.5 mm diameter) opposite of a second container filled with 10 µl of the odors amyl acetate (Sigma Aldrich, 628-63-7; diluted 1:50 in paraffin oil (Sigma Aldrich, 628-63-7)) or 1-octanol (Sigma Aldrich, 111-87-5; undiluted). The odor containers were closed with a perforated lid. 30 larvae were transferred to the middle of the petri dish and were given 5 min of free motion before being counted. Again, three groups emerge: larvae located on the odor side (#odor), larvae on the side with an empty container (#empty container) and larvae in the middle (10 mm wide stripe in the middle of the petri dish). The total amount of larvae resulted from the number of larvae in each group (#total). Preference index is calculated as follows: #odor minus #empty container divided by #total. Positive preference indices represent an attraction towards a gustatory or olfactory stimuli.

### Anatomical expression analysis

Trojan Exon lines were crossed with *20xUAS-6xmcherry/CyO; nsyb-LexA,LexAop-GFP* to investigate native fluorescence signal omitting the need of immunolabelling. 1^st^ instar larvae were collected in phosphate-buffered saline (PBS; Sigma Aldrich, P4417) and starved for 1 h. Afterwards, they were placed in 4 % bleach for 10 min, rinsed twice with H_2_O and incubated another 5 min in H_2_O. Larvae were fixated in 4 % paraformaldehyde in PBS (PFA; Sigma Aldrich, F8775) followed for 15 min at room temperature and 15 min on ice. A washing step included incubation in 3 % Triton-X in PBS (PBT; Sigma Aldrich, X100) for 15 min, repeated for three times. Finally, larvae were washed three times in PBS for 15 min. Specimens were embedded in VECTASHIELD® PLUS (Vector Laboratories, H-2000).

3^rd^ instar larvae were collected in PBS and starved for 2 h. Larvae were then heat fixed by immersion in 70 °C water for 1 min and subsequent cooling in cold water. After drying, samples were mounted in Vectashield and imaged immediately.

Brains of 3^rd^ instar wandering stage larvae were dissected in PBS on ice and fixated in 4 % PFA for 25 min under vacuum at room temperature. Afterwards, brains were washed several times in 3 % PBT and several times in PBS. Brains were mounted on poly-L-lysine (Sigma Aldrich, P1524-25MG) coated cover slips and embedded in Vectashield. Specimens were scanned at a Confocal laser scanning microscope LSM800 by Zeiss with ZEN 2.3 software and processed for publication with Image J.

### Gene models for Trojan Exon lines

Gene sequences, transcripts and annotations for *Tβh* and octopamine receptor genes were obtained via the Flybase Sequence Downloader (Öztürk-Çolak et al., 2024; release: FB2019_05). Each transcript of the respective genes listed in FlyBase is shown in the gene models in Figure 6. Conserved domain predictions were performed with NCBI conserved domain search (Wang et al., 2023) using the nucleotide sequence of each isoform of the genes. The Trojan Exon Cassette Sequence was acquired from the supplementary information from Diao et al., 2015. To annotate the sequence of the Trojan Exon, we followed Diao et al., 2015, supplementing this with data from previous publications (Venken et al., 2011; Diao and White, 2012). Location of the transgenic insertion in the gene can be found on their respective entries on Flybase. Insertion and visualization of the Trojan Exon construct sequence into the receptor gene sequences were done using Benchling [Biology Software] (2020-2025), retrieved from https://benchling.com.

### Molecular evaluation – PCR and qualitative RT-PCR

Genomic DNA was isolated using the kit NucleoSpin® Tissue; DNA, RNA, and protein purification; Genomic DNA from Tissue (Macherey-Nagel, 740952.250) following the manufacturers protocol. RNA was isolated using TRIzol™ Reagent (Invitrogen, 15596026) following the manufacturers protocol. The second step comprised DNase digestion (Roche, 04 716 728 001) followed by RNA purification using the RNeasy® MinElute® Cleanup kit (Qiagen, 74204) which were performed as stated in the manuals. The RevertAid First Strand cDNA Synthesis kit (Thermo Scientific, K1621) was used for cDNA synthesis (performed as in manufacturer protocol). Primer used for a PCR resulting in a product overlapping genomic region and Trojan Exon are as follows: Trojan Exon 5‘-TGCAGGCTATCCATCTTCAGG-3‘, *Oamb* 5‘-CGATAAAC-GCTGAGGAAGCAC-3‘, *Octα2R* 5‘-TGGTGTGATGTGTGTGGAGT-3‘, *Octβ1R* 5‘-GAGGAAGAGCCGGGAAAGTG-3‘, *Octβ2R* 5‘-GTAGCCAGGAGCAAAACAGA-3‘, *Octβ3R* 5‘-TCCTTTCAAAAGCGGTGCGA-3‘, *Oct-TyrR* 5‘-GTTTAA-GCCGATTGAGCCAC-3‘. Primer located in the *Tβh* genomic region surrounding the Trojan Exon were as follows: *Tβh* forward 5‘-GGAAGGAGAGCGTCTGAGAT-3‘ and *Tβh* reverse 5‘-TCACTGACGACAACGAACGA-3‘. Primer for PCR at 3‘ site of Trojan Exon in *Octβ2R*: *Octβ2R* genomic region 5‘-GCCGCCTTTCATTCACCATT-3‘ and in Trojan Exon 5‘-TACAACGCCTTCGGCATCAC-3‘. Primer for PCR with *Octβ2R* mRNA as template were: *Octβ2R* genomic region 5‘-CGGCGCAGGAT-TAAGTGACCA-3‘ and in Trojan Exon 5‘-TGCAGGCTATCCATCTTCAGG-3‘. *Taq* PCR Master Mix (QIAGEN, 201443) was used to amplify target regions according to the manufacturer protocol. PCR products were separated during gel electrophoresis (1 % agarose gel (Sigma Aldrich, A9539), 1xTBE buffer, MassRuler DNA Ladder Mix (Thermo Scientific, SM0403)). Gel extraction of amplified products was performed with QIAquick Gel Extraction Kit (QIAGEN, 28706) and were sequenced by Microsynth Seqlab®. Sequences for primer design and evaluation of PCR products were retrieved from the online database FlyBase (FB2021_06) (Larkin et al., 2021). For sequence management Benchling [Biology Software]. (2021) was used (https://benchling.com). Primer were designed using Primer Basic local alignment search tool (BLAST) by National Center for Biotechnology Information (Ye et al., 2012; https://www.ncbi.nlm.nih.gov/tools/primer-blast/). Sequenced PCR products were compared to the core nucleotide database and nucleotide collection by National Center for Biotechnology Information (Altschul et al., 1990; https://blast.ncbi.nlm.nih.gov/Blast.cgi?PROGRAM=blastn&PAGE_TYPE=Blast-Search&LINK_LOC=blasthome).

### Quantitative real-time RT-PCR

Total RNA from L3 larvae was isolated using QIAzol (Qiagen, 79306) with following DNA digestion using RNase-Free DNase Set (Qiagen, 79256) and purification using RNeasy Mini Kit (Qiagen, 74104), all steps were performed according to the manufacturers protocol. 250 µg RNA was reverse transcribed using SuperScript™ III First-Strand Synthesis SuperMix for qRT-PCR (Invitrogen, 11752050). Gene expression was measured by quantitative real-time RT-PCR and the TaqMan methodology (Dm02139710_g1 for *Octß2R* and Dm01825538_m1 for *Oct-TyrR*). Fluorescence was detected on an ABI PRISM 7500 sequence detector (Applied Biosystems, Foster City, CA, USA) according to the manufacturer’s instructions (Applied Biosystems). mRNA expression was calculated relative to the mRNA expression of *Rpl32* (Applied Biosystems; assay Dm02151827_g1 for *Rpl32*) and to *y^1^w\** control larvae.

### Bulk RNA-seq analysis

Raw sequencing reads were obtained from the European Nucleotide Archive (https://www.ebi.ac.uk). The RNA-seq runs used in this study are listed in Supplementary Table 8. Quality control of raw FASTQ files was performed using FastQC (v0.11.9), and reads were pseudo-aligned to the reference transcriptome (Drosophila melanogaster, genome assembly BDGP6.54, Ensembl release 113) using Salmon (v1.10.2) in quasi-mapping mode with sequence-specific and GC bias correction ena-bled (–seqBias, –gcBias). Transcript-level expression estimates were imported into R (v4.3.3) using the importIsoformExpression function from the IsoformSwitchAnalyzeR package (v2.2.0). Gene-level expression values were derived by summing transcript-level TPMs for each gene. Genes with low expression were filtered prior to down-stream analyses. Specifically, genes were retained only if they showed a gene-level expression ≥ 5 TPM in at least 3 samples. For comparative and visualization purposes, gene expression values were summarized as median TPM per tissue and developmental stage. TPM values were log-transformed where indicated for improved visualization.

## Statistical analysis

GraphPad Prism 8.0 (Graphpad Software Inc.) was used for statistical analysis and visualization. First, normal distribution of data points was assessed with the Shapiro-Wilk test. If a normal distribution was assumed, a one-way ANOVA followed by a Tukey honestly significant difference post hoc test for pairwise comparison between more than two groups were applied. If assumptions of normal distribution were not met, data was analyzed with a Kruskal-Wallis test and Dunńs multiple pairwise com-parison for more than two groups. Experimental groups were compared to the genetic background line, *y* w*.* The Wilcoxon signed-rank test was chosen to compare groups against chance level. Significance level was set at 0.05. Box plots are shown with 25 % and 75 % quantiles as box boundaries while whiskers display the complete data set (min to max values). The median is indicated as middle line and each dot represents a data point. Bar plots show the mean and standard deviation of the data set. Additionally, each dot represents a data point.

## Results

In order to use the Trojan Exon OA and OAR lines for the analysis of the OA system, we investigated whether known phenotypes could be reproduced. We tested for I) the prominent role of Tβh, Oamb and Octβ2R in female sterility (Monastirioti et al., 1996; Lee et al., 2003; Monastirioti, 2003; Lim et al., 2014; Rezaval et al., 2014; Deady and Sun, 2015; Li et al., 2015), II) impaired pupation formation in *Octβ3R* mutants (Ohhara et al., 2015) and III) disrupted larval locomotion in *Tβh, Octβ1R*, *Octβ2R* and *Oct-TyrR* larvae (Saraswati et al., 2004; Fox et al., 2006; Koon and Budnik, 2012; Selcho et al., 2012). We integrated these tests in the A) developmental assays, comprising female fecundity, larval hatching as well as larval and pupal survival (Figure 1); B) behavioral assays, comprising chemosensation, locomotion and feeding behavior (Figure 2); and C) anatomical mapping, comprising the central and peripheral nervous system as well as non-neuronal tissue (Figure 3, 4). A molecular verification of this mutant set supported the evaluation of the Trojan Exon OA and OAR lines (Figure 6). In addition to the experiments with OA and OAR mutants, gene expression analysis, based on bulk RNA-sequencing data, revealed unknown expression locations and showed quantitative expression differences in different development stages and various tissues and cells (Figure 7).

**Figure 1:**
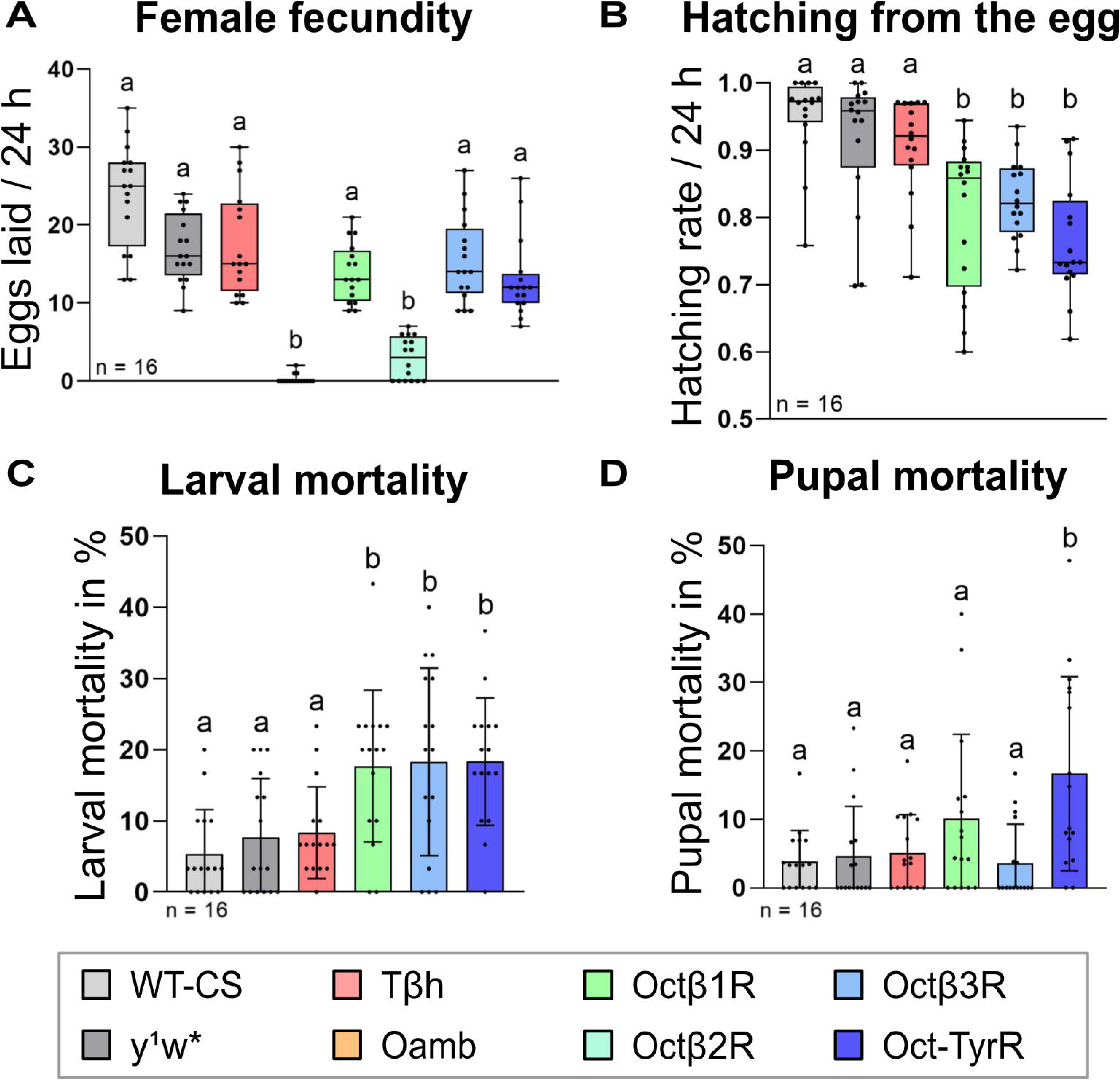
Trojan Exon octopamine receptor mutant lines have an overall reduced survival capacity. **(A)** Female fecundity was assessed by allowing mated 3-to 4-day old single female flies to lay eggs for 24 h. Any defects in male reproductive capacity were excluded by letting the females mate with WT males. The number of laid eggs was diminished only in Trojan *Oamb* (p < 0.0001) and *Octβ2R* (p < 0.0001) females compared to the genetic control *y^1^w\**. **(B)** In a second approach with homozygous animals, the ability of larvae to hatch from the egg was analyzed. Females laid eggs for 24 h and were then removed from the assay plate. After another 24 h, the number of hatched and unhatched eggs was counted. The hatching rate was calculated as the number of hatched larvae per total number of laid eggs. The number of hatched larvae of Trojan *Octβ1R* (p = 0.0081), *Octβ3R* (p = 0.0158) and *Oct-TyrR* (p = 0.0002) was significantly reduced. Trojan *Oamb* and *Octβ2R* had to be excluded since homozygous females did not lay enough eggs to compare with the control. Hatched 1^st^ instar larvae were used to determine larval and pupal mortality in c) and d). **(C)** Larval survival was assessed by transferring 30 1^st^ instar larvae to a standard food vial and counting the number of emerging pupae. Larval mortality was calculated as the number of larvae that did not pupate (in %). Trojan *Octβ1R* (p = 0.0246), *Octβ3R* (p = 0.0388) and *Oct-TyrR* (p = 0.0121) mutants showed an increased larval mortality. **(D)** Pupal survival was based on the number of flies that were able to hatch from the pupa. Pupal mortality was calculated as the number of adult flies that did not hatch (in %). Trojan *Oct-TyrR* (p = 0.0069) mutants showed an increased pupal mortality.

**Figure 2:**
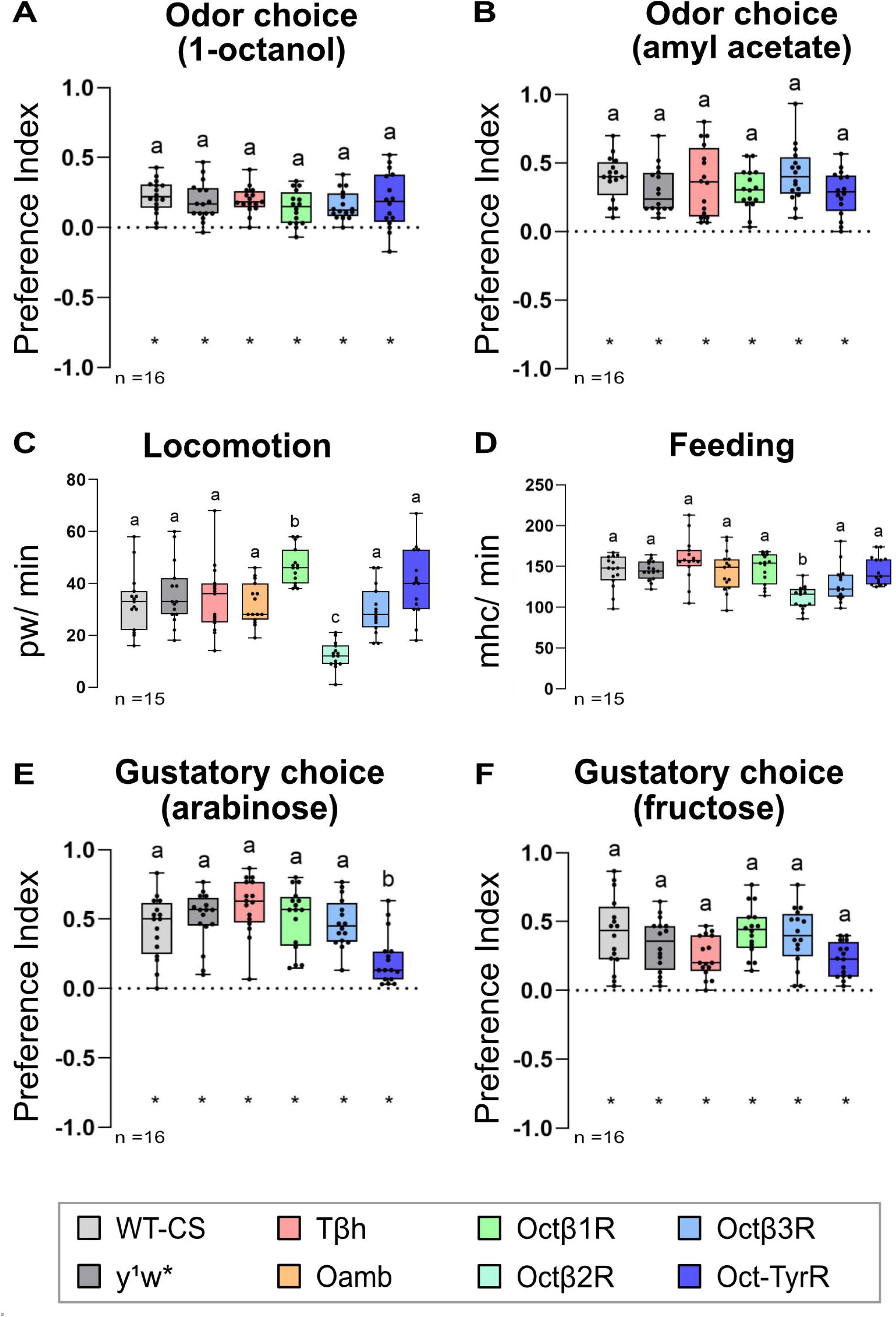
Locomotion is impaired only in Trojan *Octβ1R* and *Octβ2R*, while chemosensory behavior is not affected. Odor and taste preference tests were con-ducted with each 30 larvae per data point for which one preference score was calcu-lated. To determine locomotion and feeding rate, one larva was used per data point. **(A)** In a choice assay, larvae’s preference for the odor 1-octanol (undiluted) over an empty container after 5 min exposure is shown. Each Trojan Exon octopamine recep-tor line had a similar preference to the odor as the control. **(B)** In a choice assay, larvae’s preference for the odor amyl acetate (1:50 dilution in paraffin oil) over an empty container after 5 min exposure is shown. Each Trojan Exon octopamine recep-tor line had a similar preference to the odor as the control. **(C)** Locomotion behavior was analyzed by quantifying the peristaltic waves per minute (pw/min) of a moving larva on an agarose plate without any food source. Trojan *Octβ1R* showed significantly more pw/min (p = 0.0257) while *Octβ2R* showed significantly less pw/min (p < 0.0001). **(D)** Mouth hook retractions can be utilized as an indirect read out of feeding. Surround-ing the larva with a rich food environment makes it possible to distinguish between mouth hook retractions used for crawling and for feeding. Eliminating the need to move forward, in this assay larval mouth hook retractions were used to determine the ability of food uptake. Trojan Exon octopamine receptor mutants did not differ from the con-trol in their feeding rate. Only Trojan *Octβ2R* showed a significantly decreased number of mouth hook retractions per minute (mhc/min; p = 0.0008). **(E)** The preference for the sugar arabinose (2 M) over neutral agarose after 5 min of food uptake was tested. Arabinose has a sweet taste but is not nutritious and has no caloric value for the larva. Only Trojan *Oct-TyrR* showed a reduced preference (p = 0.0036) yet preferred this sugar over plain agarose. **(F)** The preference for the sugar fructose (2 M) over neutral agarose after 5 min of food uptake was tested. Fructose is a sweet and nutritious sugar.

**Figure 3:**
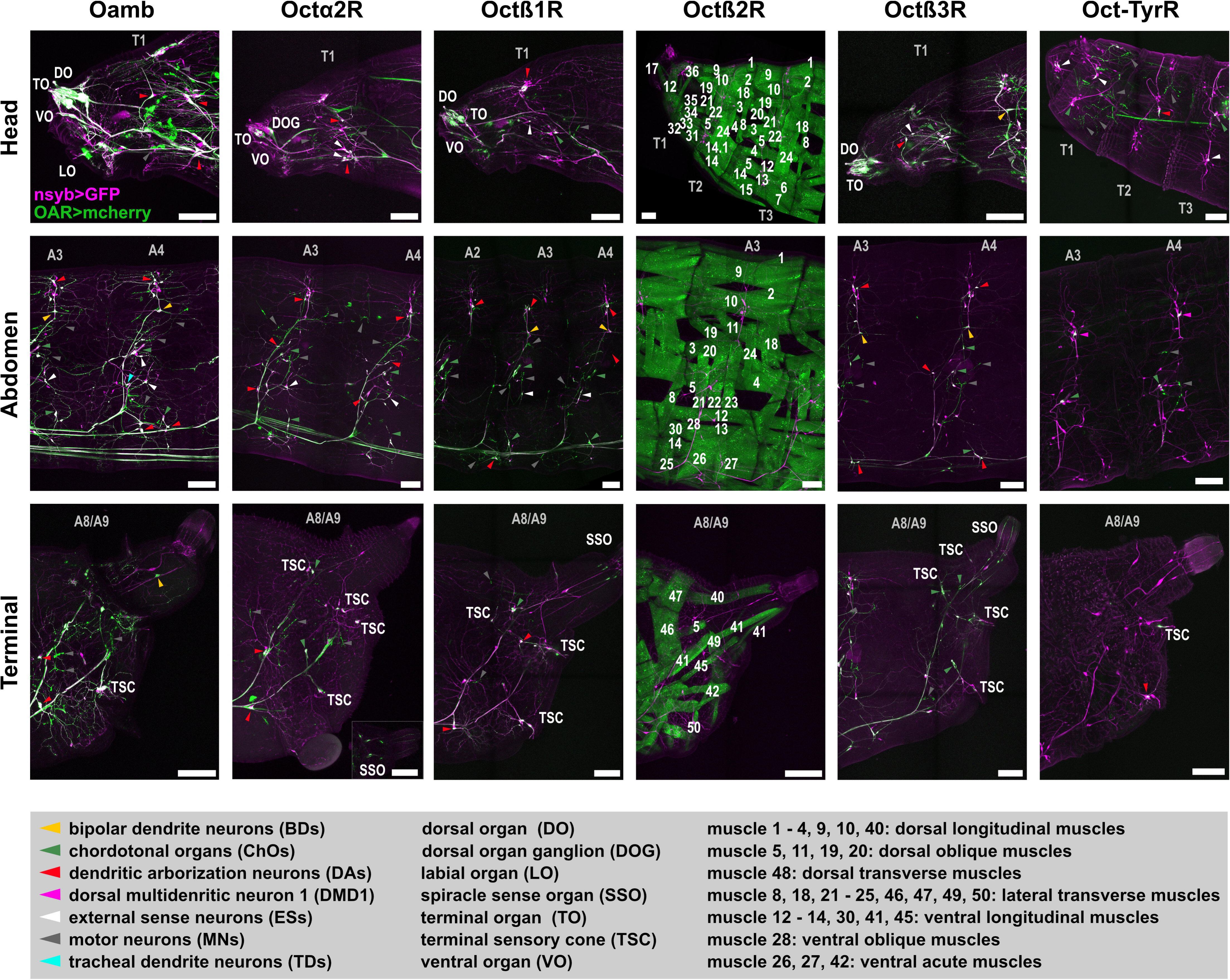
Expression pattern of each octopamine receptor throughout the body of Trojan Exon lines. Using native fluorescence, the expression pattern of each oc-topamine receptor (green via mcherry) and neuronal Synaptobrevin, as a reference channel (magenta via GFP), is shown. For each Trojan Exon line, the head (pseudo-cephalon: PCE; and up to thoracic segment 3: T3), abdominal segment 3 (and A4) and abdominal segment A8/9 is shown. In Trojan *Octα2R*, signal in SSO is shown in an extra inlet due to technical reasons. In Trojan *Octβ2R*, extensive muscle labelling was observed. Additional muscle labelling of *Octβ2R* is shown in Supplementary Fig-ure 2. Expression pattern in 1^st^ instar larvae is shown in Supplementary Figure 3. Scale bar: 100 µm.

**Figure 4:**
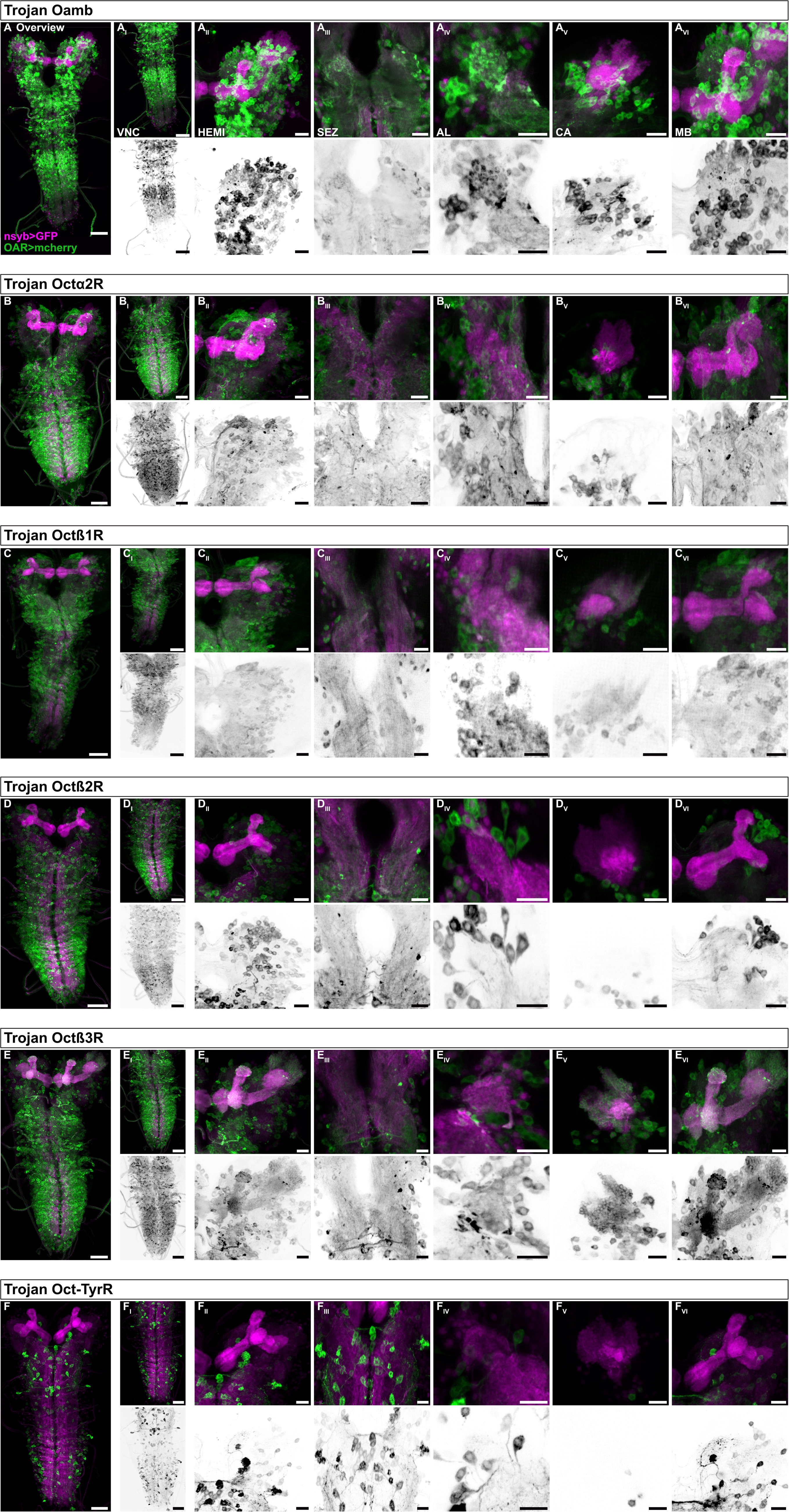
Expression pattern of each octopamine receptor in the central nervous system of Trojan Exon lines. Using native fluorescence, the expression pattern of each octopamine receptor (green via mcherry) and neuronal Synaptobrevin, as a ref-erence channel (magenta via GFP), is shown. For all lines the first panel shows an overview of the central nervous system. Close ups of the ventral nerve cord (VNC), one hemisphere (HEMI), the subesophageal zone (SEZ), the antennal lobe (AL), the calyx (CA) and the mushroom body (MB) are shown from left to right. Below each panel is a grayscale image only of the mcherry channel to show the innervation pat-terns without the reference channel. Additional images of cell bodies in the superior medial protocerebrum are shown in Supplementary Figure 1. Scale bar: 50 µm (Over-view and VNC panel), 20 µm (HEMI, SEZ, AL, CA, MB panels).

**Figure 5:**
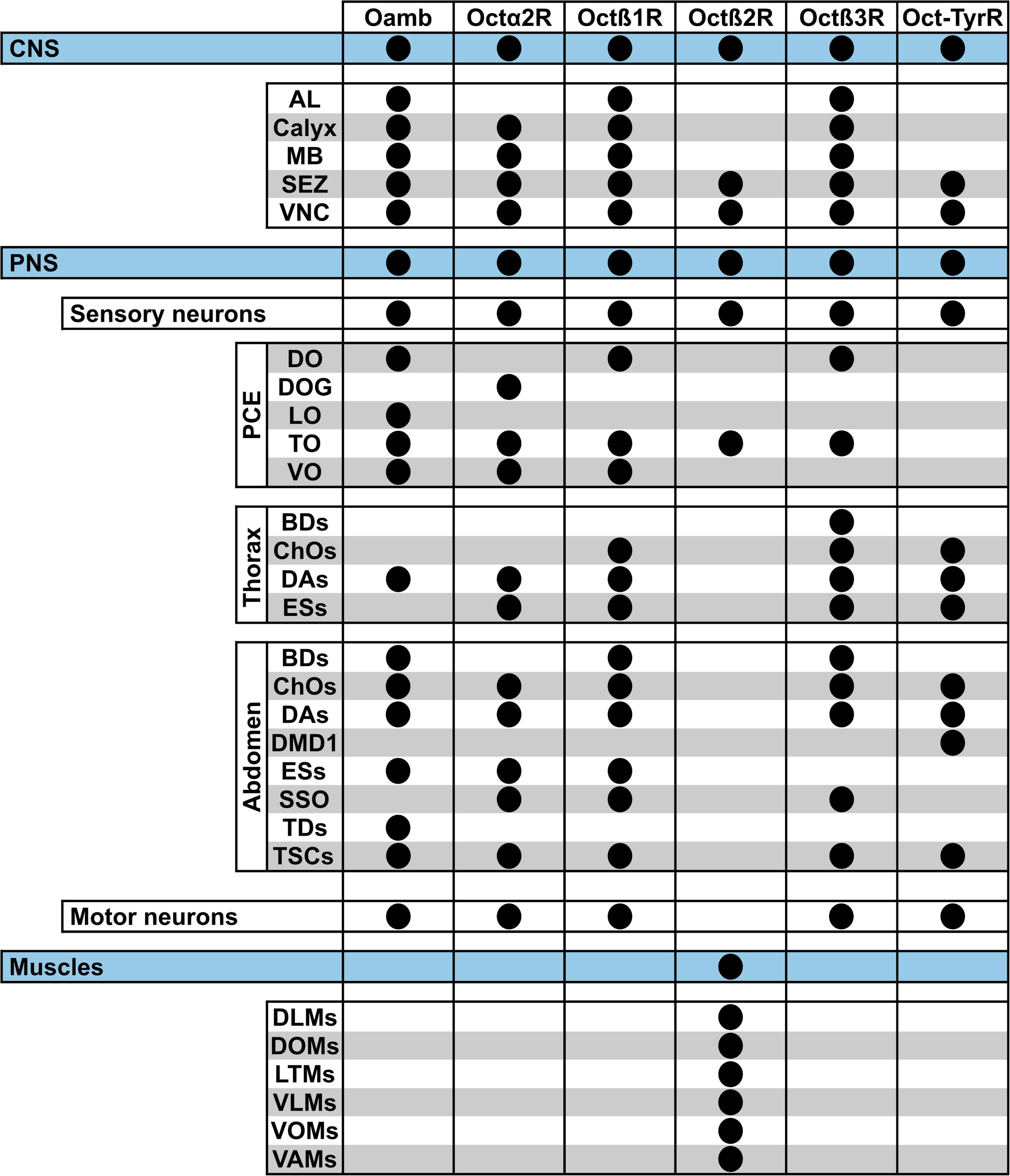
Overview of expression patterns of Trojan Exon octopamine receptor lines throughout the body. Summary of expression in the central nervous system (CNS), peripheral nervous system (PNS, split in sensory neurons and motor neurons) and somatic muscles of the body-wall of each Trojan Exon octopamine receptor line. AL: Antennal lobe; BDs: bipolar dendritic neuron; ChOs: chordotonal organs; DAs: dendritic arborization neurons; DLM: dorsal longitudinal muscle; DMD1: dorsal multi-dendritic neuron 1; DO: dorsal organ; DOG: dorsal organ ganglion; DOM: dorsal oblique muscle; ESs: external sensory neuron; LO: labial organ; LTM: lateral trans-verse muscle; MB: Mushroom body; SEZ: Subesophageal zone; SSO: spiracle sense organ; TDs: tracheal dendrite neuron; TO: terminal organ; TSCs: terminal sensory cones; VAM: ventral acute muscle; VLM: ventral longitudinal muscle; VNC: Ventral nerve cord; VO: ventral organ; VOM: ventral oblique muscle.

**Figure 6:**
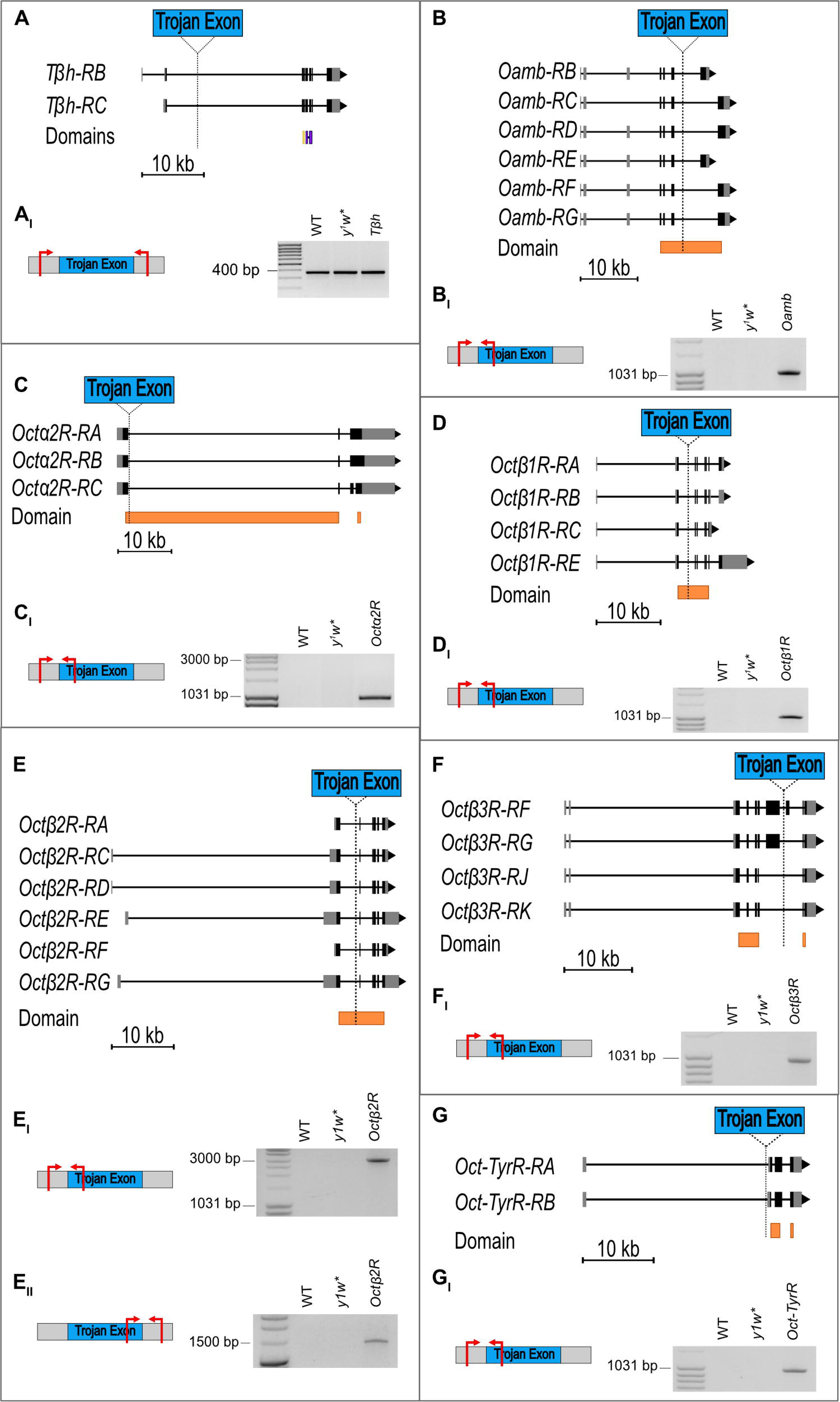
Evaluation of the correct insertion of the Trojan Exon in the mutant lines. (A)-(G) A schematic of the genes *Tβh, Oamb, Octα2R, Octβ1R, Octβ2R, Octβ3R* and *Oct-TyrR*. For each gene, the structure of each isoform with 5‘UTR (grey), Exons (black), Introns (space between Exons) and 3‘UTR (grey) is depicted. In the bottom, the predicted conserved domains and their location in the gene is marked. The insertion location of the Trojan Exon is indicated in the isoforms (dashed line). **(A_I_)-(G_I_)** On the left, a schematic shows the location of the primer (red arrows) used for PCR corresponding to the Trojan Exon (blue) in the gene (grey). On the right, vis-ualization of the amplified product during PCR. For (A_I_), first, a PCR with one primer in the gene and one in the Trojan Exon (same strategy as for the octopamine recep-tors) showed no amplified products (data not shown). In a second approach, a product of the same size was observed in the WT, *y^1^w\** and Trojan *Tβh*. Sequencing revealed that the three amplicons were identical (Supplementary Figure 4). For (B_I_)-(G_I_) and (E_II_), no product was amplified in the WT and *y^1^w\** control since the Trojan Exon, in which one primer is binding, is not present. For each Trojan Exon mutant line, a prod-uct was received and sequenced which confirmed the insert to be present and at the correct location (Supplementary Figure 5). Only for Trojan *Octβ2R*, the amplicon was of bigger size than expected (around 2000 bp; E_I_). **(E_II_)** Due to that, a second PCR was conducted to check for the 3‘ side of the Trojan Exon. Again, additional sequences were found (around 130 bp). Sequencing confirmed the insertion of additional se-quences at the 5‘ (E) and 3‘ (E_II_) side of the Trojan Exon (Supplementary Figure 6, 7), although in a BLAST analysis those sequences were not found to be coding (Supple-mentary Table 5, 6).

**Figure 7:**
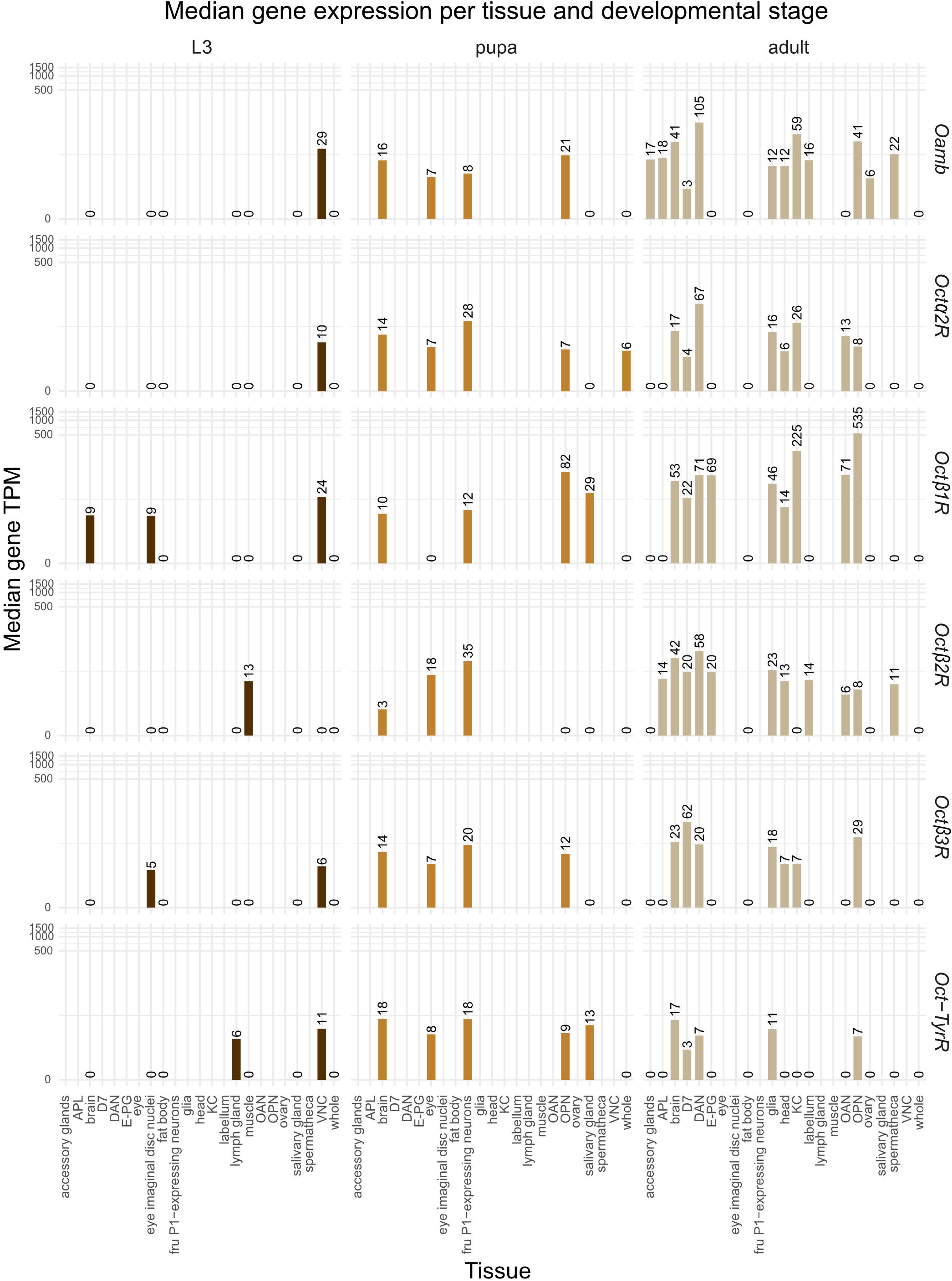
Median gene expression across tissues and developmental stages. Median transcript abundance (TPM) was calculated for all analyzed genes in each tissue and developmental stage (L3 larva, pupa including prepupa, and adult). Tissue–stage combinations with no data are left blank. Median TPM values were log-trans-formed for visualization. Sample numbers per tissue and stage are as follows: L3 larva: brain (hemispheres and VNC; n = 56), eye imaginal disc nuclei (n = 3), fat body (n = 35), lymph gland (n = 3), muscle (n = 14), salivary gland (n = 17), ventral nerve cord (VNC; n = 8), whole body (n = 183). Pupa (including prepupa): brain (n = 6), eye (n = 18), fru P1-expressing neurons (n = 16; only males), olfactory projection neurons (OPN; n = 3), salivary gland (n = 13), whole body (n = 76). Adult: olfactory projection neurons (OPN; n = 3), anterior paired lateral neuron (APL; n = 5), brain (n = 74), Δ7 neurons (D7; n = 4), dopaminergic neurons (DAN; n = 14), E-PG neurons (n = 4), fat body (n = 12), glia (n = 7), head (n = 201), Kenyon cells (KC; n = 9), labellum (n = 3), octopaminergic neurons (OAN; n = 154), ovary (n = 81), spermatheca (n = 6), whole body (n = 444).

### Octopamine receptors contribute to the animal’s viability – from egg to adult

First, we performed a general assessment of the animal’s viability over the different developmental stages. Embryonic mortality was examined based on the ability of the 1^st^ instar larva to hatch from the egg. Larval and pupal mortality was used to evaluate survival rates. Additionally, female sterility was analyzed applying an egg-laying test, in which the number of eggs laid by individual flies within 24 h was counted. To exclude potential effects arising from male reproductive system defects (as has been reported for Oamb (Zhou et al., 2012; Luo et al., 2014)), mutant females were mated with WT males. A significantly reduced number of eggs was observed for Trojan *Oamb* and *Octβ2R* compared to the genetic control. In contrast, no reduction was seen for Trojan *Tβh*, *Octβ1R*, *Octβ3R* and *Oct-TyrR* (Figure 1A). Due to the limited number of experimental animals resulting from the small number of eggs laid (Figure 1A) and the low percentage of homozygous animals (Supplementary Table 1), further analysis of larval development in Trojan *Oamb* and *Octβ2R* was not possible. For all other groups, we assessed the ability of mutant larvae to hatch from the egg and calculated a hatching rate, which is defined as the number of hatched larvae divided by the total number of eggs. This time, females were mated with males of the same genotype to analyze only homozygous mutant larvae. The hatching rate was found to be reduced in Trojan *Octβ1R*, *Octβ3R* and *Oct-TyrR* compared to control animals while no reduction was observed in *Tβh* (Figure 1B). Subsequently, the survival of larvae and pupae was monitored. Larval mortality was assessed as the percentage of larvae that did not enter the pupal stage. A significant increase was observed in Trojan *Octβ1R*, *Octβ3R* and *Oct-TyrR* (Figure 1C). Pupal mortality, which indicates the percentage of animals unable to hatch from the pupa, showed that only Trojan *Oct-TyrR* had a reduced number of hatched flies, reflected in an increased pupal mortality (Figure 1D). Raw data and statistical analysis for all data presented in Figure 1 can be found in Supplemen-tary able 2. So, on the one hand, the Trojan *Oamb* and *Octβ2R* lines reproduced the well-known female sterility phenotype; on the other hand, for the first time, it was shown that Octβ1R and Oct-TyrR are involved in larval hatching and larval survival. Even the survival of pupae was impaired in Trojan *Oct-TyrR*. We also observed a decreased survival rate of larvae in Trojan *Octβ3R*, which could be related to the reported impairment of pupation (Ohhara et al., 2015). Nevertheless, for the first time we showed that Octβ3R is involved in larval hatching.

### Analysis of larval foraging behavior

In addition to physiological mechanisms, changes in behavior can also explain the decreased viability of Trojan Exon mutants observed in Figure 1. Therefore, we aimed to analyze the role of the OARs in one such behavior – foraging. During the larval stage, energy requirements are high due to a body mass increase by about 200 times (Bakker, 1959). Consequently, foraging plays an essential role in the life of larvae, ensuring their survival and progression into metamorphosis (Bakker, 1961; Ohnishi, 1979). In this study, we subdivide the larval foraging behavior in I) the perception of olfactory cues of a potential food source (Figure 2A, b), II) locomotion on pure agarose (Figure 2C), III) the ability of food uptake (Figure 2D) and IV) the gustatory preference for food sources that are generally indicative of high energy supply (Figure 2E, F). A first encounter with a potential food source can be olfactory cues, these help the larva to navigate in their environment and initiate and/or maintain contact with a substrate (reviewed in Cobb, 1999; shown in Gomez-Marin et al., 2011; Gershow et al., 2012). Here, we tested the preference for the two odors amyl acetate and 1-octanol, which can be associated with a food source in an olfactory associative learning and memory assay (reviewed in Gerber and Stocker, 2007; shown in Selcho et al., 2009; Pauls et al., 2010; Schleyer et al., 2011; Apostolopoulou et al., 2013; Weber et al., 2023a; We-ber et al., 2023b). We used the same experimental setup and odor concentrations as in Apostolopoulou et al., 2013). Larvae from each Trojan Exon line showed a preference for the two odors, similar to control animals, suggesting that olfactory perception for these odors and the tested concentrations is unaffected (Figure 2A, B). It is important to note that due to the small number of laid eggs, which resulted in a limited number of experimental animals, Trojan *Oamb* and *Octβ2R* could not be included in these mass assay experiments.

After perceiving an appetitive odor cue, the crawling behavior is directed more specifically towards a potential food source (Fishilevich et al., 2005; Louis et al., 2008). In a locomotion assay, we evaluated the crawling ability by quantifying the peristaltic waves (pw) as a read-out of rhythmic motor patterns (Gjorgjieva et al., 2013; Liu et al., 2023). Tested animals were neither exposed to external stimuli nor treated in any other way. While Trojan *Octβ1R* exhibited an increased number of pw/min, Trojan *Octβ2R* showed significantly less pw/min than the control animals (Figure 2C). Given that these assays analyze individual larvae, it was possible to include the relatively small number of surviving animals of Trojan *Oamb* and *Octβ2R*.

The next assay focused on assessing the feeding behavior of the larvae. In contrast to crawling behavior, feeding primarily relies on contractions of the cephalopharyngeal muscles, which are separate from the somatic segment muscles in the abdomen (Schoofs et al., 2010). However, mouth hook contractions (mhc) are also required for crawling behavior. In the feeding assay, larvae were placed in a nutrient-rich environment that eliminated the need to search for food. Mouth hook contractions serve as a straightforward indicator of feeding behavior (Bhatt and Neckameyer, 2013). Most Tro-jan Exon lines showed no change in feeding rate, only *Octβ2R* larvae had a significantly reduced number of mhc per minute compared to control animals (Figure 2D). Gustatory preferences can indicate whether a larva is able to identify beneficial sources of nutrition, such as sweet and high caloric food (Rohwedder et al., 2012). In a choice assay, larvae were presented sugar on the one side and tasteless, non-nutritious agarose on the other side. We used two different sugars, I) fructose which is sweet and has a high caloric value, while II) arabinose is only sweet but non-nutritious. All Trojan Exon lines preferred the sugars over the neutral agarose (Figure 2E, F). However, Trojan *Oct-TyrR* showed a significantly reduced preference for arabinose. Please note that due to the small number of laid eggs, which resulted in a limited number of experimental animals, Trojan *Oamb* and *Octβ2R* were not included in these mass assay experiments. Raw data and statistical analysis for all data in Figure 2 can be found in Supplementary Table 3.

### Expression pattern of Trojan Exon OAR lines in the larva

Utilizing the simultaneous driven Gal4 expression in the Trojan Exon lines, we anatomically mapped the expression of each OAR in neuronal and non-neuronal cells. Two protocols were applied, one to evaluate the respective expression throughout the body (Figure 3) and the other in the CNS (Figure 4) of 3^rd^ instar larvae. We map only unambiguous signals, mostly at the body surface, but labelling of deep-seated tissues cannot be excluded. In most Trojan Exon lines, we observed extensive labelling in the CNS which made it difficult to determine the identity of individual cells.

Trojan *Oamb* showed labelling in the dorsal organ (DO), labial organ (LO), terminal organ (TO) and ventral organ (VO) in the pseudocephalon (PCE) (Figure 3). Each segment, from thoracic segment 1 (T1) to abdominal segment 8/9 (A8/9), featured signal in dendritic arborization neurons (DAs) as well as motor neurons (MNs). Additional labelling was found in bipolar dendrite neurons (BDs), chordotonal organs (ChOs), external sense neurons (ESs) and tracheal dendrite neurons (TDs) in the abdomen, as well as in the terminal sensory cones (TSC) in A8/9. Oamb was observed to be the only OAR expressing in TDs. The CNS showed broad labelling from the hemispheres (HEMI, Figure 4A_II_) to the subesophageal zone (SEZ, Figure 4A_III_) and the ventral nerve cord (VNC, Figure 4A_I_) but it gets weaker in the posterior VNC ganglia that represents abdominal segments seven and eight (Figure 4A, A_I_). A cluster of cell bodies that project to the antennal lobe (AL) expressed Oamb (Figure 4A_IV_) as well as a few Kenyon cells (KC) in the calyx (CA, Figure 4A_V_) of the mushroom body (MB, Figure 4A_VI_).

In Trojan *Octα2R*, both TO and VO labelling can be found in the PCE (Figure 3). While the DO itself is not labelled, signals were detected in a group of sensory neurons belonging to the dorso-lateral group of the TO, which have their cell bodies within the associated ganglion (DOG). DAs, ESs and MNs showed Octα2R expression both in thoracic and abdominal segments. ChOs were solely labelled in the abdomen. The specialized sensory structures of the fused terminal abdominal segments A8/A9, TSC and spiracle sense organ (SSO), showed Octα2R expression, too. An extensive signal was observed in the VNC of the CNS, which was less strong in the hemispheres, although homogeneously visualized cell bodies surrounding specialized structures such as the AL and MB (Figure 4B, B_I-VI_). Only single KC, likely embryonic born (Pauls et al., 2010), were found to express Octα2R (Figure 4B_V_). Two distinct bilateral cell clusters were found in the superior medial protocerebrum, with somata in the pars intercerebralis and projections extending along the midline of each hemisphere (in more detail in Supplementary Figure 1). This pattern corresponds to neurosecretory cells that release insulin (insulin producing cells (IPCs); compare Rulifson et al., 2002; Cao et al., 2014). However, further analysis is needed to confirm the identity of this cell cluster.

Trojan *Octβ1R* showed expression in DO, TO and VO in the PCE, and in ChOs, DAs, ESs and MNs in each thoracic and abdominal segment (Figure 3). Additionally, BDs were labelled in the abdominal segments, and TSCs and SSO were observed to express Octβ1R. In the CNS, we found many cell bodies labelled, from the hemispheres to the VNC, mostly surrounding the AL and MB (Figure 4C, C_I-VI_). But a few KC showed weak signal (Figure 4C_V_). Similar to Trojan *Octα2R*, a cell cluster in the superior medial protocerebrum of each hemisphere may resemble IPCs (Figure 4C_II_).

Extensive Octβ2R signals can be found in somatic muscles of the body-wall across each thoracic and abdominal segment (Figure 3). In the PCE, two intersegmental muscles showed labelling: muscle 12, which runs from dorsal T1 to ventral PCE, and muscle 17, which extends from laterofrontal PCE to dorsal T2. In the prothorax (T1), we identified dorsal longitudinal muscle 36 and ventral longitudinal muscles 31-35, which are associated with the mouthparts (Hooper, 1986). In the mesothorax (T2), signal was observed in dorsal longitudinal (muscle 1-4, 9, 10), dorsal oblique (muscle 5 and 19; muscle 20 is probably covered by muscle 19), lateral transverse (muscle 8, 18, 21, 22, 24; muscle 23 is probably covered by muscle 22 and 24) and ventral longitudinal muscle (muscle 14). Only dorsal oblique muscle 30 appears to be not labelled. In the metathorax (T3), we identified dorsal longitudinal (muscle 1-4, 9, 10), dorsal oblique (muscle 5, 19, 20), lateral transverse (muscle 8, 18, 21, 22, 24; muscle 23 is probably covered by muscle 22 and 24), ventral longitudinal (muscle 6, 7, 12, 13, 14) and ventral oblique muscles (muscle 15). Only ventral acute muscle 26 and dorsal oblique muscle 30 showed no Octβ2R signal. Each muscle of the body-wall in abdominal segments A1-7 expressed Octβ2R in the Trojan Exon line and can be summarized as follows since the muscle pattern is identical for each segment: dorsal longitudinal (muscle 1-4, 9, 10), dorsal oblique (muscle 5, 11, 19, 20), lateral transverse (muscle 8, 18, 21-25), ventral longitudinal (muscle 6, 7, 12-14, 30), ventral oblique (muscle 15-17, 28) and ventral acute (muscle 26, 27, 29). Due to their position, labelling of muscles 6, 7, 15, 16, 17 and 29 is shown in Supplementary Figure 2A. Muscle 25 is the only muscle not present in A1 (Zarin et al., 2019) and hence was not found during the expression analysis. In segment A8/9, dorsal longitudinal muscle 40, dorsal transverse muscle 48 (Supplementary Figure 2B), lateral transverse muscles 46, 47, 49 and 50, ventral longitudinal muscle 41 and 45 as well as ventral acute muscle 42 were identified. These muscles can be grouped in I) associated with the terminal spiracle (muscle 40, 41 and 42) and II) associated with the anal division (muscle 45, 46, 47, 49 and 50) depending on their position (Wipfler et al., 2013). Only one muscle in A8/9 was not expressing Octβ2R, the ring-shaped muscle 43 which surrounds the terminal spiracles. According to (Wipfler et al., 2013), we found a muscle labelled as number 5, although it was not described in detail. Due to the uncertainty of their position, the identity of two additional muscles (44 and 51) was not determined. Muscles 44 and 51 have been previously described but not visualized (Wipfler et al., 2013). Muscle identity was determined based on the work of Wipfler et al., 2013 for PCE and A8/9, Hooper, 1986 for T1-3, and Zarin et al., 2019 for A1-7. Additionally, we identified a signal in at least one sensory neuron in the TO (Supplementary Figure 2C). While additional expression may also occur in other sensory neurons, motor neurons, or non-neuronal cells, the very strong signal in the somatic muscles could be overshadowing these other signals. The CNS of Trojan *Octβ2R* showed broad signal in the VNC, though it was less pronounced in the hemispheres (Figure 4D). We found paired cell bodies along the mid-line of the SEZ (Figure 4D_III_), single cell bodies around the AL (Figure 4D_IV_) and a cell cluster lateral to the vertical lobe of the MB, but no labelled KC (Figure 4D_V,VI_).

In Trojan *Octβ3R*, signals were observed in both the DO and TO within the PCE (Figure 3). Octβ3R expression was detected in the thoracic and abdominal segments in certain neuron types, including BDs, ChOs, DAs, ESs (only thorax) and MNs. In segment A8/9, ChOs and MNs as well as SSO and TSCs were labelled. In the CNS, a broad expression signal encompassed all segmental ganglia within the VNC, with less pronounced signal in the hemispheres of the brain (Figure 4E). The SEZ showed cell bodies along the midline and lateral towards the VNC (Figure 4E_III_). While we observed a signal in only a few cells in and around the AL (Figure 4E_IV_), many KC in the CA and the lobes of the MB were distinctively labelled (Figure 4E_V__, VI_). Additional cell bodies in the hemispheres, around the MB expressed Octβ3R (Figure 4E _VI_).

In thoracic and abdominal segments, we observed ChOs, DAs, ESs and MNs in Trojan *Oct-TyrR* larvae (Figure 3). Signal in the dorsal multidendritic neuron 1 (DMD1) was observed in the abdomen, and other OARs were not expressed in this particular neuron type. Similar to most OARs, Oct-TyrR showed expression in TSCs in segment A8/9. The CNS showed the most specific expression pattern among the Trojan Exon OAR lines. Cell bodies in the abdominal segments were positioned laterally, while the thoracic segments have additional cell bodies along the midline (Figure 4F). In the SEZ, we found cell bodies specifically near the esophagus (Figure 4F_III_). The hemispheres showed only little Oct-TyrR signal and no distinct labelling of the AL or MB (Figure 4F_IV, V, VI_).

A summary of the expression locations throughout the entire body and the CNS can be found in Figure 5. Additional whole-body scans of first instar larvae show similar, yet less pronounced expression patterns compared to the third instar larvae (Supplementary Figure 3). It remains unclear whether these differences are due to developmental stages or technical factors.

### Molecular analysis reveals absence of Trojan Exon in Tβh line

*Tβh* mutant lines are known to exhibit severe egg laying impairments as well as larval locomotor deficits (Monastirioti, 2003; Saraswati et al., 2004; Selcho et al., 2012), prominent behavioral phenotypes that were not observed in Trojan *Tβh*. Moreover, during expression pattern analysis, no signal was observed in the brain, which contradicted an earlier study that found extensive labelling applying immunohistochemistry (Selcho et al., 2012). In Trojan *Tβh*, the Trojan Exon is located in the first half of the *Tβh* gene locus (based on FlyBase entry ID FBti0195387) resulting in a premature truncation of the protein (Figure 6A). The conserved domains DOMON (dopamine beta-monooxygenase N-terminal; yellow) and the Copper type II ascorbate-dependent monooxygenase (blue) were predicted to be located downstream of the Trojan Exon insertion site (Supplementary Table 4) and are crucial for the enzymatic function of Tβh (Figure 6A). It can therefore be assumed that the truncated Trojan *Tβh* protein is non-functional, which is why we expected to find previously known phenotypes of *Tβh* mutants. Thus, we performed targeted PCR to analyze the Trojan Exon on the genomic level and found that the Trojan *Tβh* line lost the cassette. According to the published insertion site of the Trojan Exon in the *Tβh* gene locus, MI04010, we used a primer pair that surrounded the *Tβh* MI04010 docking site. In case of the presence of the Trojan Exon, we expected an amplified product of around 4 kb. But, as shown in Figure 6A_I_, the size of the amplicon was similar to that of the WT and the *y^1^w\** controls. Sequencing of the amplified PCR product revealed no differences between Trojan *Tβh* and the WT *Tβh* gene locus of the reference genome acquired from FlyBase (Supplementary Figure 4). We repeated the targeted PCR with a different copy of the Trojan *Tβh* fly line, ordered from the Bloomington Drosophila Stock Center (BDSC) and received identical results (data not shown).

Based on these results, we continued analyzing the Trojan Exon OAR lines to ensure the correct insertion of the Trojan Exon. This time, a different primer strategy was applied: based on the MiMIC docking sites (*Oamb*: MI12417, *Octα2R*: MI10227, *Octβ1R*: MI05807, *Octβ2R:* MI13416, *Octβ3R*: MI06217, *Oct-TyrR*: MI03485), one primer was located in the genomic region and one in the Trojan Exon (see schematic in Figure 6 for each OAR). Due to the size of the amplified products and sequencing results (Supplementary Figure 5), we conclude that the location of the Trojan Exon in the *Oamb*, *Octα2*, *Octβ1*, *Octβ2, Octβ3* and *Oct-Tyr* receptor lines is identical with the corresponding MiMIC docking sites. Only for Trojan *Octβ2R*, the amplicon size was larger than expected (expected: 1072 bp, observed: ∼ 3000 bp; Figure 6E_I_). A second PCR at the 3‘end of the Trojan Exon in the *Octβ2R* gene locus showed a similar result (expected: 1366 bp, observed: ∼1500 bp; Figure 6E_II_). Sequencing revealed that the additional parts were not endogenous to the *Octβ2R* gene and BLAST analysis showed no distinct origin (Supplementary Figure 6, 7 and Supplementary Table 5, 6). We further analyzed if the additional sequences upstream of the 5‘end of the Trojan Exon were transcribed and could affect the formation of the Trojan Exon. A schematic (not to scale) shows the unknown additional parts (∼ 2000 bp) right upstream of the Trojan Exon at the genomic level (Supplementary Figure 8A_I_). In a qualitative RT-PCR, primers were designed to bind in the upstream Exon 7 and in the Trojan Exon. If the inserted sequences were excluded from transcription, an amplicon of 532 bp was expected. The observed PCR product, visualized below the schematic, migrated between 500 and 600 bp. Subsequent sequencing of this fragment confirmed the absence of the additional sequences in the *Octβ2R* mRNA of the Trojan *Octβ2R* line. (Supplementary Figure 9). Moreover, quantitative real-time RT-PCR was performed to analyze *Octβ2R* mRNA level. We applied the TaqMan methodology with a probe binding downstream of the inserted Trojan Exon (Supplementary Figure 8A). Only a fraction (∼6 %) of *Octβ2R* mRNA was detected compared to the control, showing the decreased expression of this OAR in the Trojan Exon mutant (Supplementary Figure 8C).

In each tested line, the Trojan Exon is inserted into the first half of the respective OAR gene (Figure 6), suggesting that the Trojan Exon induced protein truncation leads to the loss of major structural parts of the receptor. Conserved domain prediction analysis revealed that the conserved transmembrane domains of the GPCRs are affected (Figure 6; Supplementary Table 4), suggesting that the OARs are non-functional and thus that the Trojan Exon OAR lines are suitable mutants for investigating the OAR system. Only for *Oct-TyrR*, the insert location of the Trojan Exon is located in an intron in the non-coding 5‘UTR. Since the Trojan Exon must be inserted into an intron between two coding exons, the intended effect of truncating the Oct-TyrR protein and thereby disrupting its functionality is questionable. But, quantitative real-time RT-PCR showed that no *Oct-TyrR* transcript was detectable in the Trojan Exon mutant indicating that the Oct-TyrR protein is absent (Supplementary Figure 8B, C). Further, due to the insert location in an intron in the 5‘UTR, the expression pattern obtained during anatomical mapping may not correspond to the endogenous pattern of Oct-TyrR.

All in all, insertion sites were confirmed in most Trojan Exon OARs making these lines suitable genetic tools for interfering with the OA system. Even though the insertion site of the Trojan Exon in *Octβ2R* is compromised, the mutant seems to be valid based on our mRNA analysis. For Trojan *Oct-TyrR*, we found that the targeted protein disruption takes place even if it may not have been in the intended manner. But due to the insert location in an intron in the 5‘UTR, Gal4 induced expression may not correspond to the endogenous Oct-TyrR pattern. Only Trojan *Tβh* cannot be used for further studies because it lacks the Trojan Exon.

### Gene expression data

All results obtained so far are based on experiments with Trojan Exon mutants. In a different approach, we utilized bulk RNA-sequencing data retrieved from the European Nucleotide Archive to focus on OAR expression location and quantity in L3 larva, pupa and adult, each in different tissues and cell populations (Figure 7).

In L3, expression of Oamb, Octα2R, Octβ3R and Oct-TyrR was more pronounced in the VNC than the hemispheres, in contrast to Octβ1R that showed a high abundance throughout the brain (hemispheres and VNC). Increased levels of Octβ1R and Octβ3R were observed in eye imaginal disc nuclei, while Octβ2R showed increased expression in L3 muscle. A yet unknown expression location was found for Oct-TyrR in the lymph gland. In the pupa, every OAR had elevated transcript level in the brain and in a neuronal subset, the fru P1-expressing neurons of the male. The pupal eye showed expression of each OAR except Octβ1R, and olfactory projection neurons (OPN) were found to express all OARs except Octβ2R at increased levels. High levels of Octβ1R and Oct-TyrR were observed in salivary glands of the prepupa. Throughout the body, Octα2R was the most abundant OAR. In the adult, whole head samples showed elevated transcript levels for each OAR except Oct-TyrR. But in the brain, Oct-TyrR transcript level were increased as well, indicating that head tissue surrounding the brain does not express Oct-TyrR as much as the other OARs. In some neuron populations, every OAR was observed at high abundance, namely: Δ7 neurons (D7) of the protocerebral bridge, dopaminergic neurons (DAN), glia and olfactory projection neurons (OPN). Other neuron populations were found to express some OARs more than others, such as KC (Oamb, Octα2R, Octβ1R and Octβ3R); octopaminergic neurons (OAN) (Octα2R, Octβ1R and Octβ2R); the anterior paired lateral neuron (APL; Oamb and Octβ2R) and the so-called E-PG neurons in the central complex (Octβ1R and Octβ2R). Similarly, non-neuronal tissue was found to express max. two OARs at elevated levels: labellum and spermatheca express the same OARs (Oamb and Octβ2R), and accessory gland and ovary express only Oamb. In addition, due to the assignment of cell identity in RNA-sequencing data, we were able to find so far unknown expression locations in neuronal cells: APL (Oamb, Octβ2R), DAN (Octα2R, Octβ1R, Octβ2R, Octβ3R and Oct-TyrR), Δ7 neurons (Oamb, Octα2R, Octβ1R, Octβ2R, Octβ3R and Oct-TyrR), E-PG neurons (Octβ1R and Octβ2R), glia (Oamb, Octα2R, Octβ1R, Octβ3R and Oct-TyrR), KC (Octα2R), OPN (Oamb, Octα2R, Octβ1R, Octβ2R, Octβ3R and Oct-TyrR), and in non-neuronal cells: labellum (Oamb, Octβ2R). Interestingly, Octβ1R was found at exceptionally high transcript levels in adult OPN (median gene TPM: 535) and in KC (median gene TPM: 225) compared to all other expression levels in the different tissues and cells as well as for other OARs. Only Oamb in DAN (median gene TPM: 105) might be comparable.

Overall, each OAR was expressed in fewer tissues and cells in L3 compared to the pupa and adult. And although the median transcript abundance in general was low in the whole body in each stage (with Octα2R being the only exception in the pupa), expression was elevated in specific tissues and cell populations, varying from L3 to adult.

## Discussion

The Trojan Exon provides the possibility to analyze a functionally disrupted target gene and at the same time localize its gene product by using the Gal4/UAS system. Using this powerful genetic tool, we were able to break down the analysis from the entire OA circuit to the receiver side and focus on each receptor individually. We identified novel phenotypes of OARs in larval hatching and the survival of larvae and pupae. Moreover, we expanded the expression map of OARs to I) cells and tissues which until now were not considered to be a potential place of action of OA, II) to the 1^st^ instar larva which were not reported before and III) to Octα2R which has not been examined in the larva. An overview of the observed expression patterns reported in earlier studies and the results from this study are summarized in Supplementary Table 7. The molecular evaluation of the Trojan Exon OARs not only offers a better understanding of the results presented in this study but also provides a solid basis for future investigations of the OA system using these genetic tools. Likewise, the results show that Trojan Exon lines generally require molecular validation to ensure that the gene of interest is disrupted in the intended manner.

### Tβh

Prominent phenotypes of *Tβh* mutants are impaired female sterility (Monastirioti et al., 1996; Monastirioti, 2003; Rezaval et al., 2014) and larval locomotion (Saraswati et al., 2004; Fox et al., 2006; Selcho et al., 2012), among others. We analyzed the performance of Trojan *Tβh* for these two selected traits and couldńt find any impairments (Figure 1A; Figure 2C). Utilizing the Gal4 expression in Trojan Exon lines, anatomical characterization of Tβh in the Trojan *Tβh* line was not feasible after a cross with several UAS reporters (data not shown). After molecular evaluation, it became clear that the Trojan *Tβh* line had lost the Trojan Exon (Figure 6A_I_). Even re-ordering the line showed the same results. Therefore, the Trojan *Tβh* line available from Bloomington Drosophila Stock Center cannot be used to examine the function of OA in *Drosophila*.

### Oamb

Oamb is known for its prominent role in the female reproductive system, receiving the OA signal required for ovulation. Mutants deprived of Oamb lay significantly fewer eggs (Lee et al., 2003; Deady and Sun, 2015), which was also observed in the Trojan *Oamb* mutant (Figure 1A). The high abundance of Oamb in ovary and spermatheca reflect the female sterility phenotype observed in *Oamb* mutants (Figure 7). The egg laying phenotype, together with molecular evaluation (Figure 6), verifies the Trojan *Oamb* to be a suitable mutant for further studies and supports our findings from expression analyses (Figure 3; Figure 4A). We observed extensive labelling in the larval CNS (utilizing the Trojan Exon mutant; also reported by El-Kholy et al., 2015) with Oamb being more abundant in the VNC than the hemispheres (Figure 7). But the PNS showed also a broad signal: beginning in the PCE, gustatory (TO, VO), olfactory (DO), mechanosensory (DO, LO, TO, VO), thermosensory (DO), chemosensory (oxygen; TO) and osmosensory (TO) organs were identified (for further information on the function of the different organs see Rist and Thum, 2017, and Richter et al., 2023). Tho-racic and abdominal segments were found to have signal in neurons and organs for proprioception (DAs; Ferreira Castro et al., 2020; He et al., 2019; ChOs; Caldwell et al., 2003), innocuous touch (DAs; Hu et al., 2017; Tsubouchi et al., 2012) as well as noxious stimuli such as cold, heat, touch, light or acidity (DAs; Tsubouchi et al., 2012). Recently, a function for Oamb in DAs was found in nociceptive plasticity (Boivin et al.,2026). ESs in the abdomen indicate further involvement in the perception of mechanical and/or chemical stimuli (Jan and Jan, 1993). Labelled TDs, that wrap tracheal branches, and TSCs at A8/9 suggest a role in the respiratory system since these structures are associated with the perception of CO_2_ and/or oxygen (Bodmer and Jan, ^1^98^7^; Hückesfeld et al., 2021; Lu et al., 2024; Richter et al., 2023). Even the neuro-muscular system is affected by Oamb, as expression was found in BD neurons associated with somatic muscles (Bodmer and Jan, 1987) and in motor neurons (also reported by Jetti et al., 2023; Bakshinska et al., 2025). These findings identify Oamb as one of the most broadly distributed OARs in the larval periphery, suggesting a yet unknown function in the sensory system (for further discussion see last paragraph).

### Octα2R

With the Trojan *Octα2R* line, it is now possible to create a first anatomical expression map in the larva (Figure 3; Figure 4B). In the PCE, Octα2R was observed in the DOG, TO and VO suggesting involvement in gustatory, mechanosensory, chemosensory and/or osmosensory processes (see above for further details). Across the body, we found signal in DAs, ESs, and MNs, as well as in ChOs in the abdominal segments; thus, Octα2R can also be associated with proprioception and/or nociception, mech-ano-/chemosensory and the neuromuscular system. At the terminus, Octα2R labelled the TSCs and SSO. Richter et al., 2023 discussed the function of the TSCs and SSO in the positioning of the larva in a food medium to hide from predators and daylight but in the same way avoid suffocation from digging too deep (Kim et al., 2017). Although the exact role of OA in this survival mechanism has yet to be elucidated, the expres-sion of Octα2R may constitute an initial indicator of its potential involvement.

### Octβ1R

Embryonic development comprises 17 stages, with organ differentiation beginning at stage 13 and continuing through to stage 16. During this phase, elevated *Octβ1R* transcript levels were reported in the CNS and PNS (Berkeley Drosophila Genome Project; Tomancak et al., 2002; Tomancak et al., 2007). In stages 15-17, Ohhara et al., 2012 observed increased mRNA levels of *Octβ1R* in the CNS and in the entire embryo. Our research revealed a decreased percentage of hatched eggs in Trojan *Octβ1R* (Figure 1B). Together, these findings suggest a functional involvement of Octβ1R. Furthermore, overall larval mortality is increased in Trojan *Octβ1R* (Figure 1C). In this assay, we did not discriminate between the different larval stages. However, in 3^rd^ instar larvae we observed expression in gustatory (TO, VO), olfactory (DO), mechanosensory (DO, TO, VO), thermosensory (DO), chemosensory (TO) and osmosensory (TO) organs (Figure 3; see section Oamb for further details). Further, Octβ1R function in proprioception (ChOs and DAs), nociception (DAs), mechano-/chemosensation (ESs) or the neuromuscular system (BDs and MNs; MN labelling was also shown by Koon and Budnik, 2012) could have influenced larval survival. Maybe even defects in the TSCs and SSO could have caused detrimental positioning of the larva in the food medium leading to a misjudgment of oxygen availability. However, the broad labelling in the CNS (also shown by Koon and Budnik, 2012; Ohhara et al., 2012; El-Kholy et al., 2015), even in specialized structures such as the AL and MB, should not be overlooked when analyzing the role of Octβ1R in larval mortality (Figure 4C). Specifically, the high abundance of this receptor in the whole CNS and eye imaginal disc nuclei (compared to other OARs) suggests a functional role in the nervous system and in development (Figure 7). Additionally, earlier studies reported this receptor in the Malpighian tubules, the tracheal system (El-Kholy et al., 2015), the salivary glands (Ohhara et al., 2012) and the eye-antennal discs (Ohhara et al., 2012; Koon and Budnik, 2012). Whether it is a defect in a single tissue or a combination of tissues that leads to increased mortality remains to be analyzed.

Larval movement features characteristic peristaltic waves. In *Tβh^nM18^* larvae, a reduction in peristaltic waves during crawling has been correlated with reduced forward movement and shorter distance traveled (Saraswati et al., 2004). In the applied locomotion assay, without any stimuli or treatment, we associated the number of peristaltic waves within 1 min with baseline locomotor activity. Trojan *Octβ1R* larvae were the only tested group with increased baseline locomotor activity (Figure 2C). In contrast, *octβ1r* mutants were shown to have decreased basal locomotor activity in a similar environment (a pure agarose plate without any stimuli) and a fed state (Koon and Budnik, 2012). Apart from the agarose concentration (2.5% vs. 3%), we are not aware of any significant methodological differences. So, Octβ1R function in larval locomotion needs further investigation, ideally using different mutants in the same assays and conditions or even utilizing tissue-or cell-specific manipulation of this receptor for a detailed analysis.

RNA-seq analysis revealed that the OARs are expressed at low level in different tissues and developmental stages, only Octβ1R showed exceptionally high abundance and only in specific cell populations - KC and OPN in adult. Octβ1R expression has been reported in the MB (El-Kholy et al., 2015; McKinney et al., 2020), where it drives aversive learning; while located in OPN, Octβ1R processes appetitive learning (Sabandal et al., 2020).

### Octβ2R

In the Trojan Exon line, we observed two known phenotypes associated with *Octβ2R* mutants. The first is associated with the role of Octβ2R in ovulation, in which mutant females have a deficit in egg laying (Lim et al., 2014; Li et al., 2015; Figure 1A). The second shows a reduction in basal locomotion behavior in 3^rd^ instar larvae (Koon and Budnik, 2012; Figure 2C). Octβ2R was found to be highly abundant in larval muscle tissue (Figure 7) and muscle labelling has also been reported in another promotor line (El-Kholy et al., 2015). However, with the Trojan *Octβ2R* line, individual muscles can be identified from the head to the tail. We found that all 29 somatic muscles of abdominal segment 1 and all 30 somatic muscles of abdominal segments 2-7 in the body-wall express Octβ2R (Figure 3). During crawling, all somatic muscles in the body-wall are activated (Zarin et al., 2019). Thus, Octβ2R seems to play a general role in basal locomotor activity. Moreover, muscles of the head (PCE and T1-3) showed extensive labelling, indicating a further role of Octβ2R in larval movement such as turning or burrowing behavior. For feeding behavior, we found not only labelling of mouthpart associated muscles in the prothorax (Figure 3; Hooper, 1986) but also a reduction of the feeding rate in the Trojan Exon mutant (Figure 2D). Moreover, even the respiratory and excretory system might involve Octβ2R function, since somatic muscles associated with the terminal spiracles and anal division were identified (Figure 3). In addition to the broad expression in somatic muscles throughout the body, Octβ2R was reported to function as an auto-receptor in type II motor neurons and to receive plasticity-mediating OA signaling in type I motor neurons (Koon et al., 2011). The Trojan Exon mutant did not show any labeling of the MNs, though the sensitive structures could have been overlooked by the excessive signal in the muscles. This could have also happened for the earlier reported expression in the fat body, intestine, imaginal disc, Malpighian tubule, midgut, salivary gland and tracheal system, which was found utilizing molecular techniques (RT-PCR and in situ hybridization; Ohhara et al., 2012; El-Kholy et al., 2015). The only other signal in the periphery was observed in the TO, probably because it was not covered by any muscle labeling.

Molecular evaluation of the Trojan *Octβ2R* line showed additional sequences 5‘ and 3‘ of the Trojan Exon (Supplementary Figure 8A_I_). The additional sequences did not appear to be coding regions or parts of the *Octβ2R* gene itself. Qualitative RT-PCR revealed that the size of the *Octβ2R* transcript did not change due to these additional sequences. Further, due to the female sterility, the larval locomotion and feeding phenotype, and the utilization of Gal4 during the expression analysis, the additional sequences do not appear to affect the transcriptional and/or translational processes of the *Octβ2R*-Trojan Exon fragment. Quantitative real-time RT-PCR showed decreased *Octβ2R* mRNA level. Thus, despite these genetic inconsistencies, the Trojan *Octβ2R* is a suitable mutant line to investigate this receptor and its role in the OA system in more detail.

### Octβ3R

A function of Octβ3R emerged as early as the transition from embryo to larva. The hatching rate of 1^st^ instar larvae from the egg was reduced in the Trojan Exon line (Figure 1B). *Octβ3R* mRNA level have been observed to be elevated in embryonic stages 15-17 in the CNS/whole animal (Ohhara et al., 2012). Although, the role of Octβ3R in embryo development has yet to be investigated, the reduced hatching rate and elevated expression levels in the embryonic stage warrant further studies. In L3 larvae, Octβ3R is broadly expressed in both internal and external sensory organs, responding primarily to chemo-and mechanosensory signals. This indicates that Octβ3R plays a functional role in various sensory modalities, including gustation (TO), olfaction (DO), mechanosensation (DO, ESs, TO), thermosensation (DO), chemosensation (ESs, TO), osmosensation (TO) and proprioception (ChOs, DAs). Additionally, it is involved in peripheral neurons related to nociception (DAs) and muscle activity (BDs, MNs) (see Figure 3 and section Oamb for more details). Moreover, Octβ3R was found in TSCs and SSO, which are believed to respond to changes in oxygen levels or to give mechanosensory feedback when the larvae submerge into the substrate. When such an array of information is not properly detected, adjustments to environmental conditions may be affected and thus could affect larval survival (Figure 1C). Additionally, the extensive labelling (Figure 4) and elevated transcript level (Figure 7) in the CNS should not be overlooked, as well as the high abundance in eye imaginal disc nuclei (Figure 7). Though, as summarized in Supplementary Table 7, earlier reports have Octβ3R observed even in further tissues, including different glands, the Malpighian tubule and the midgut.

During our assessment of larval mortality, we focused on counting the number of emerging pupae rather than quantifying the number of deceased larvae. Earlier, it has been reported that Octβ3R plays a crucial role in inducing metamorphosis by regulating ecdysone biosynthesis in the prothoracic gland (Ohhara et al., 2015). However, the pupariation defect was less pronounced in the Trojan *Octβ3R* than in the earlier report. This discrepancy may be attributed to technical differences or variations in the genetic tools employed. Further investigations are necessary to determine the specific time point and location of Octβ3R function that affects larval survival and pupariation.

### Oct-TyrR

Similar to Trojan *Octβ1R* and *Octβ3R*, 1^st^ instar larva hatching ability was reduced in Trojan *Oct-TyrR* (Figure 1B). Unfortunately, data regarding the transcript level of *Oct-TyrR* in the embryo is not available. Given that both OA and TA are potential ligands, the role of TA in embryo development and larval hatching should also be considered. Beyond that, later stages of larval and pupal development showed reduced survival rates (Figure 1C, D). The expression of Oct-TyrR in ChOs, DAs, DMD1, ESs and TSCs suggests a potential involvement in proprioception (ChOs, DAs and DMD1 (Heckscher et al., 2015)), as well as mechano-and chemosensation and oxygen perception (ESs and TSCs, Figure 3; see sections above for further details). This corresponds with an earlier reported phenotype in chemotaxis behavior and the startle response in *honoka* larvae, a different *Oct-TyrR* mutant (Ma et al., 2016). Moreover, this receptor was found throughout the body, including the CNS, dorsal aorta, lymph gland, Malpighian tubule, posterior spiracles, salivary gland and the tracheal system (Stowers, 2011; El-Kholy et al., 2015; Figure 4F; Figure 7). It remains unclear whether a functional role in the sensory system, the different tissues, or both contributes to the increased larval mortality. Further, we identified a potential role of Oct-TyrR in pupal survival based on increased pupal mortality of Trojan *Oct-TyrR* mutants (Figure 1D). Interestingly, RNA-sequencing analysis revealed that transcript *Oct-TyrR* level were elevated not only in the pupal CNS and eye but also in the salivary glands (Figure 7). While other OARs showed similar results, it remains to be elucidated why one OAR has an impact on pupal survival and others have not. But, specifically the high abundance of *Oct-TyrR* in salivary glands of the prepupa, which is a major secretory organ that undergoes autophagy during the early phases of metamorphosis, suggests a function of OA in development.

A role for Oct-TyrR in locomotion has first been described in *honoka* mutants that showed a reduction of excitatory junctional potential (EJP) amplitudes in body-wall muscles (Kutsukake et al., 2000; Nagaya et al., 2002). Later, expression has been found in the CNS, MNs and at the neuromuscular junction (NMJ; Stowers, 2011; Fig-ure 3). Our analysis of locomotion behavior in Trojan *Oct-TyrR* showed no impairment compared to the control (Figure 2C) which is in line with previous data of *honoka* (Schützler et al., 2019). Although, we also showed a hyperactivity phenotype in *honoka* (Selcho et al., 2012). A comparison of these two studies is difficult, since not only the controls were different but also the tracking method itself (automated (Schützler et al., 2019) vs. manual (Selcho et al., 2012)). Further research is needed to clarify the role of Oct-TyrR in larval locomotion behavior.

### The OAR system

OA plays a role in various mechanisms in *Drosophila melanogaster*, starting from the embryonic stage and continuing through larval development to adulthood. A body of literature already identified several of its receptors being involved in the same biological mechanism, such as Oamb, Octβ1R, Octβ2R and Oct-TyrR in larval locomotion (Koon et al., 2011; Koon and Budnik, 2012; Selcho et al., 2012; Bakshinska et al., 2025), Oamb, Octα2R, Octβ1R, Octβ2R, Octβ3R and Oct-TyrR in larval and adult starvation response (Koon et al., 2011;Erion et al., 2012; Koon and Budnik, 2012; Zhang et al., 2013; Luo et al., 2014; Damrau et al., 2018; Nakagawa et al., 2022), Oamb, Octβ1R and Octβ2R in adult olfactory learning and memory (Burke et al., 2012; Kim et al., 2013; Huetteroth et al., 2015; Yang et al., 2016; Sabandal et al., 2020), Oamb and Octα2R in adult arousal (Kula-Eversole et al., 2010), Oamb, Octβ1R, Octβ2R, Octβ3R and Oct-TyrR in adult sleep (Crocker et al., 2010; Deng et al., 2019 but see also Reyes et al., 2026), or Oamb and Octβ2R in ovulation (Lee et al., 2003; Deady and Sun, 2015; Lim et al., 2014; Li et al., 2015). Here, we observed an overlap of three OARs (Octβ1R, Octβ3R and Oct-TyrR) being involved in embryo development and/or larval hatching as well as larval survival. Additionally, together with previous data, we observed a broad overlap in the expression locations of the various OARs (Supplementary Table 7). The larval CNS showed specifically extensive labelling of most of the OARs and, although we did not identify individual cells, it seems likely that expression patterns of different OARs overlap (Figure 4, 5). Especially specialized structures such as the AL (Oamb, Octβ1R and Octβ3R), Calyx/MB (Oamb, Octα2R, Octβ1R and Octβ3R) and potentially IPCs (Octα2R and Octβ1R) were observed to be labelled by more than one OAR (Figure 4, 5). In the PNS, we found overlapping expression in BDs (Oamb, Octβ1R and Octβ3R), ChOs (Oamb, Octα2R, Octβ1R, Octβ3R and Oct-TyrR), DAs (Oamb, Octα2R, Octβ1R, Octβ3R and Oct-TyrR), DO (Oamb, Octβ1R and Octβ3R), ESs (Oamb, Octα2R, Octβ1R, Octβ3R and Oct-TyrR), SSO (Octα2R, Octβ1R and Octβ3R), TO (Oamb, Octα2R, Octβ1R, Octβ2R and Octβ3R), TSCs (Oamb, Octα2R, Octβ1R, Octβ3R and Oct-TyrR) and VO (Oamb, Octα2R and Octβ1R). Even an imaginal disc showed a high abundance of two OARs (Octβ1R and Octβ3R; Figure 7). Moreover, although Octβ2R does not share most of the expression locations with the other OARs, muscles associated with the mouthparts in the PCE suggest involvement in feeding related behaviors, similarly to OARs found in DO, TO and VO. On the other hand, muscles associated with the posterior spiracles have been discussed as contributing to larval positioning in the food medium while digging, thereby playing a crucial role in respiration (Dombrovski et al., 2017), similarly to OARs in SSO and TSCs. In the pupa and adult, RNA-sequencing analysis revealed that several tissues/cell populations express multiple OARs at high abundance: the pupal eye (all OARs except Octβ1R), neuronal subsets, e.g. pupal fru P1-expressing neurons (all OARs) and OPN (all OARs except Octβ2R), and the prepupal salivary gland (Octβ1R and Oct-TyrR). Adult neuronal subsets include D7, DAN, glia, OPN (all OARs), KC (Oamb, Octα2R, Octβ1R and Octβ3R), OAN (Octα2R, Octβ1R and Octβ2R), APL (Oamb and Octβ2R), E-PG (Octβ1R and Octβ2R; Figure 7).

One experimental approach to understanding the functional implications of the overlapping OAR expression was to apply chemosensory assays. The sensory system in the larval PCE showed particularly broad expression of OARs in the DO, TO and VO. However, behavior tests do not suggest direct involvement of OA. The similarities in OAR expression patterns within the same functional unit for olfactory or gustatory perception may indicate that other OARs compensate for the absence of receptors in single OAR mutants. This is particularly plausible since the downstream signaling is conserved in GPCRs and some OARs exhibit similar G-protein coupling (see Table 1). OA is involved in various mechanisms crucial for survival, a redundant system in which a receptor is able to take over the function of another, is less prone to interference. For instance, in ovulation it has been found that ectopically expressed Octβ1R in the oviduct epithelium substitutes Octβ2R and restores ovulation in *octβ2r* females (Lim et al., 2014). However, it remains to be clarified whether there is a compensatory effect in the chemosensory system - and if so, if it occurs at the level of the functional unit or within individual cells.

On the other hand, intersecting OAR expression may rather reflect the modulatory functioning of OA, as it is found for example in the locomotory system. Muscles of the body-wall are innervated by OANs (Monastirioti et al., 1995; Hoang and Chiba, 2001; Koon et al., 2011; Selcho et al., 2012). OA signaling contributes to synaptic plasticity at the NMJ of adjacent MNs via Oamb, Octβ1R and Octβ2R (Koon et al., 2011; Koon and Budnik, 2012; Bakshinska et al., 2025; Blaum et al., 2025). Here, we observed that five out of six OARs are expressed in MNs, adding Octα2R, Octβ3R and Oct-TyrR to the list of potential OARs involved in synaptic plasticity (Figure 3, 5). While Oamb and Octβ2R have a stimulating effect on synapse potentiation, Octβ1R has an inhibi-tory effect (Koon et al., 2011; Koon and Budnik, 2012; Bakshinska et al., 2025; Blaum et al., 2025). These varying effects can be explained with different G-protein coupling and their subsequent downstream signaling pathways (Table 1). But how excitatory and inhibitory OARs can be expressed in the same cell needs further analysis. How-ever, most of the OARs have been linked to cAMP signaling. Interestingly, cAMP sig-nals have been found to be spatially confined to different compartments within MNs (Maiellaro et al., 2016). Such local signaling indicates that different OARs with different downstream signaling may be expressed within the same cell, located at different sites. But if this is the case and if this takes place also in other neuronal and non-neuronal cells remains to be elucidated. A combination of different protein tags, such as Venus (Kondo et al., 2020), at different OARs could provide insight into subcellular receptor localization in individual cells. Further, selectively manipulating individual OARs in single cells, that express multiple OARs, could shed light on whether the OA system is based on redundancy or modulation - or perhaps both - varying between cells.

## Author contributions

Conceptualization: A.G., I.C., and A.S.T; Methodology: A.G., V.R., F.R., J.B., and A.S.T; Investigation: A.G., V.R., F.R., M.H., R.B., J.K. and A.S.T; Writing: A.G., V.R., F.R., J.B., P.K., P.F.S., I.C., and A.S.T; Supervision: A.S.T.

## Acknowledgements

We thank Dennis Pauls, Wolfgang Blenau, Bert Klagges, Tilman Triphan and Wolf Hütteroth for their discussions and comments.

## References

Altschul, S.F., Gish, W., Miller, W., Myers, E.W., and Lipman, D.J. (1990). Basic local alignment search tool. Journal of Molecular Biology 215(3), 403–410. doi: 10.1016/S0022-2836(05)80360-2.

Andrews, J.C., Fernandez, M.P., Yu, Q., Leary, G.P., Leung, A.K., Kavanaugh, M.P., et al. (2014). Octopamine Neuromodulation Regulates Gr32a-Linked Aggression and Courtship Pathways in Drosophila Males. PLoS Genet 10(5), e1004356. doi: 10.1371/journal.pgen.1004356.

Apostolopoulou, A.A., Widmann, A., Rohwedder, A., Pfitzenmaier, J.E., and Thum, A.S. (2013). Appetitive Associative Olfactory Learning in <em>Drosophila<em>Larvae. Journal of Visualized Experiments (72). doi: 10.3791/4334-v.

Babski, H., Codianni, M., and Bhandawat, V. (2024). Octopaminergic descending neurons in Drosophila: Connectivity, tonic activity and relation to locomotion. Heliyon 10(9), e29952. doi: 10.1016/j.heliyon.2024.e29952.

Bakker, K. (1959). Feeding Period, Growth, and Pupation in Larvae of *Drosophila melanogaster*. Entomologia Experimentalis et Applicata 2(3), 171–186. doi: 10.1111/j.1570-7458.1959.tb00432.x.

Bakker, K. (1961). An Analysis of Factors Which Determine Success in Competition for Food Among Larvae of *Drosophila Melanogaster*. Archives Néerlandaises de Zoologie 14(2), 200–281. doi: 10.1163/036551661X00061.

Bakshinska, D., Liu, W.Y., Schultz, R., Stowers, R.S., Hoagland, A., Cypranowska, C., et al. (2025). Synapse-specific catecholaminergic modulation of neuronal glutamate release. Proc. Natl. Acad. Sci. U.S.A. 122(1), e2420496121. doi: 10.1073/pnas.2420496121.

Balfanz, S., Strunker, T., Frings, S., and Baumann, A. (2005). A family of octapamine receptors that specifically induce cyclic AMP production or Ca2+ release in *Drosophila melanogaster*. J Neurochem 93(2), 440–451. doi: 10.1111/j.1471-4159.2005.03034.x.

Berger, M., Fraatz, M., Auweiler, K., Dorn, K., El Khadrawe, T., and Scholz, H. (2024). Octopamine integrates the status of internal energy supply into the formation of food-related memories. Elife 12. doi: 10.7554/eLife.88247.

Bhatt, P.K., and Neckameyer, W.S. (2013). Functional analysis of the larval feeding circuit in *Drosophila*. J Vis Exp (81), e51062. doi: 10.3791/51062.

Blaum, N., Ghelani, T., Gotz, T.W.B., Chronister, K.S., Bengochea, M., Ceresnova, L., et al. (2025). Monoamine-induced diacylglycerol signaling rapidly accumulates Unc13 in nanoclusters for fast presynaptic potentiation. Proc. Natl. Acad. Sci. U.S.A. 122(34), e2514151122. doi: 10.1073/pnas.2514151122.

Bodmer, R., and Jan, Y.N. (1987). Morphological characterization of the embryonic peripheral neurons in *Drosophila*. Roux’s Arch Dev Biol 196, 69–77. doi: 10.1007/BF00402027.

Boivin, J.C., Zhao, Y.Q., Zhu, J., Dakin, J.T., Ning, J., and Ohyama, T. (2026). A positive feedback loop between sensory and octopaminergic neurons underlies nociceptive plasticity in Drosophila larvae. PLoS Genet 22(4), e1012122. doi: 10.1371/journal.pgen.1012122.

Branch, A., Zhang, Y., and Shen, P. (2017). Genetic and Neurobiological Analyses of the Noradrenergic-like System in Vulnerability to Sugar Overconsumption Using a Drosophila Model. Sci Rep 7(1), 17642. doi: 10.1038/s41598-017-17760-w.

Brembs, B., Christiansen, F., Pfluger, H.J., and Duch, C. (2007). Flight initiation and maintenance deficits in flies with genetically altered biogenic amine levels. J Neurosci 27(41), 11122–11131. doi: 10.1523/JNEUROSCI.2704-07.2007.

Burke, C.J., Huetteroth, W., Owald, D., Perisse, E., Krashes, M.J., Das, G., et al. (2012). Layered reward signalling through octopamine and dopamine in *Drosophila*. Nature 492(7429), 433–437. doi: 10.1038/nature11614.

Busch, S., Selcho, M., Ito, K., and Tanimoto, H. (2009). A map of octopaminergic neurons in the Drosophila brain. Journal of Comparative Neurology 513(6), 643–667. doi: 10.1002/cne.21966.

Busch, S., and Tanimoto, H. (2010). Cellular configuration of single octopamine neurons in Drosophila. J Comp Neurol 518(12), 2355–2364. doi: 10.1002/cne.22337.

Caldwell, J.C., Miller, M.M., Wing, S., Soll, D.R., and Eberl, D.F. (2003). Dynamic analysis of larval locomotion in *Drosophila* chordotonal organ mutants. Proc. Natl. Acad. Sci. U.S.A. 100(26), 16053–16058. doi: 10.1073/pnas.2535546100.

Cao, J., Ni, J., Ma, W., Shiu, V., Milla, L.A., Park, S., et al. (2014). Insight into insulin secretion from transcriptome and genetic analysis of insulin-producing cells of *Drosophila*. Genetics 197(1), 175–192. doi: 10.1534/genetics.113.160663.

Certel, S., Savella, M.G., Schlegel, D.C.F., and Kravitz, E.A. (2007). Modulation of *Drosophila* male behavioral choice. Proc. Natl. Acad. Sci. U.S.A. 104(11), 4706–4711. doi: 10.1073/pnas.0700328104.

Certel, S.J., Leung, A., Lin, C.Y., Perez, P., Chiang, A.S., and Kravitz, E.A. (2010). Octopamine neuromodulatory effects on a social behavior decision-making network in *Drosophila* males. PLoS One 5(10), e13248. doi: 10.1371/journal.pone.0013248.

Chatwin, H.M., Rudling, J.E., Patel, D., Reale, V., and Evans, P.D. (2003). Site-directed mutagenesis studies on the *Drosophila* octopamine/tyramine receptor. Insect Biochem Mol Biol 33(2), 173–184. doi: 10.1016/s0965-1748(02)00188-1.

Classen, G., and Scholz, H. (2018). Octopamine Shifts the Behavioral Response From Indecision to Approach or Aversion in Drosophila melanogaster. Front Behav Neurosci 12, 131. doi: 10.3389/fnbeh.2018.00131.

Cobb, M. (1999). What and how do maggots smell? Biological Reviews 74(4), 425–459. doi: 10.1017/S0006323199005393.

Crocker, A., and Sehgal, A. (2008). Octopamine regulates sleep in *Drosophila* through protein kinase A-dependent mechanisms. J Neurosci 28(38), 9377–9385. doi: 10.1523/JNEUROSCI.3072-08a.2008.

Crocker, A., Shahidullah, M., Levitan, I.B., and Sehgal, A. (2010). Identification of a neural circuit that underlies the effects of octopamine on sleep:wake behavior. Neuron 65(5), 670–681. doi: 10.1016/j.neuron.2010.01.032.

Damrau, C., Toshima, N., Tanimura, T., Brembs, B., and Colomb, J. (2018). Octopamine and Tyramine Contribute Separately to the Counter-Regulatory Response to Sugar Deficit in Drosophila. Front Syst Neurosci 11, 100. doi: 10.3389/fnsys.2017.00100.

David, J.-C., and Coulon, J.-F. (1985). Octopamine in invertebrates and vertebrates. A review. Progress in Neurobiology 24, 141–185. doi: 10.1016/0301-0082(85)90009-7.

Deady, L.D., and Sun, J. (2015). A Follicle Rupture Assay Reveals an Essential Role for Follicular Adrenergic Signaling in *Drosophila* Ovulation. PLoS Genet 11(10), e1005604. doi: 10.1371/journal.pgen.1005604.

Deng, B., Li, Q., Liu, X., Cao, Y., Li, B., Qian, Y., et al. (2019). Chemoconnectomics: Mapping Chemical Transmission in *Drosophila*. Neuron 101(5), 876–893 e874. doi: 10.1016/j.neuron.2019.01.045.

Diao, F., Ironfield, H., Luan, H., Diao, F., Shropshire, W.C., Ewer, J., et al. (2015). Plug-and-play genetic access to *Drosophila* cell types using exchangeable exon cassettes. Cell Rep 10(8), 1410–1421. doi: 10.1016/j.celrep.2015.01.059.

Diao, F., and White, B.H. (2012). A novel approach for directing transgene expression in *Drosophila*: T2A-Gal4 in-frame fusion. Genetics 190(3), 1139–1144. doi: 10.1534/genetics.111.136291.

Dombrovski, M., Poussard, L., Moalem, K., Kmecova, L., Hogan, N., Schott, E., et al. (2017). Cooperative Behavior Emerges among Drosophila Larvae. Curr Biol 27(18), 2821–2826 e2822. doi: 10.1016/j.cub.2017.07.054.

El-Kholy, S., Stephano, F., Li, Y., Bhandari, A., Fink, C., and Roeder, T. (2015). Expression analysis of octopamine and tyramine receptors in *Drosophila*. Cell Tissue Res 361(3), 669–684. doi: 10.1007/s00441-015-2137-4.

El-Kholy, S.E., Afifi, B., El-Husseiny, I., and Seif, A. (2022). Octopamine signaling via OctalphaR is essential for a well-orchestrated climbing performance of adult Drosophila melanogaster. Sci Rep 12(1), 14024. doi: 10.1038/s41598-022-18203-x.

Erion, R., DiAngelo, J.R., Crocker, A., and Sehgal, A. (2012). Interaction between sleep and metabolism in Drosophila with altered octopamine signaling. J Biol Chem 287(39), 32406–32414. doi: 10.1074/jbc.M112.360875.

Farooqui, T. (2012). Review of octopamine in insect nervous systems. Open Access Insect Physiology. doi: 10.2147/oaip.S20911.

Ferreira Castro, A., Baltruschat, L., Sturner, T., Bahrami, A., Jedlicka, P., Tavosanis, G., et al. (2020). Achieving functional neuronal dendrite structure through sequential stochastic growth and retraction. eLife 9, e60920. doi: 10.7554/eLife.60920.

Fishilevich, E., Domingos, A.I., Asahina, K., Naef, F., Vosshall, L.B., and Louis, M. (2005). Chemotaxis behavior mediated by single larval olfactory neurons in Drosophila. Curr Biol 15(23), 2086–2096. doi: 10.1016/j.cub.2005.11.016.

Fox, L.E., Soll, D.R., and Wu, C.F. (2006). Coordination and modulation of locomotion pattern generators in *Drosophila* larvae: effects of altered biogenic amine levels by the tyramine beta hydroxlyase mutation. J Neurosci 26(5), 1486–1498. doi: 10.1523/JNEUROSCI.4749-05.2006.

Franke, U.S., Großjohann, A., Aurich, S., Kohler, I., Lamberty, M., Granato, S., et al. (2026). Selective octopaminergic tuning of mushroom body circuits during memory formation. Proc Natl Acad Sci U S A 123(16), e2517403123. doi: 10.1073/pnas.2517403123.

Gerber, B., and Stocker, R.F. (2007). The *Drosophila* larva as a model for studying chemosensation and chemosensory learning: a review. Chem Senses 32(1), 65–89. doi: 10.1093/chemse/bjl030.

Gershow, M., Berck, M., Mathew, D., Luo, L., Kane, E.A., Carlson, J.R., et al. (2012). Controlling airborne cues to study small animal navigation. Nat Methods 9(3), 290–296. doi: 10.1038/nmeth.1853.

Gjorgjieva, J., Berni, J., Evers, J.F., and Eglen, S.J. (2013). Neural circuits for peristaltic wave propagation in crawling Drosophila larvae: analysis and modeling. Front Comput Neurosci 7, 24. doi: 10.3389/fncom.2013.00024.

Gomez-Marin, A., Stephens, G.J., and Louis, M. (2011). Active sampling and decision making in *Drosophila* chemotaxis. Nat Commun 2, 441. doi: 10.1038/ncomms1455.

Han, K.-A., Millar, N.S., and Davis, R.L. (1998). A Novel Octopamine Receptor with Preferential Expression in *Drosophila* Mushroom Bodies. J Neurosci 18(10), 3650–3658. doi: 10.1523/JNEUROSCI.18-10-03650.1998.

He, L., Gulyanon, S., Mihovilovic Skanata, M., Karagyozov, D., Heckscher, E.S., Krieg, M., et al. (2019). Direction Selectivity in *Drosophila* Proprioceptors Requires the Mechanosensory Channel Tmc. Curr Biol 29(6), 945–956 e943. doi: 10.1016/j.cub.2019.02.025.

Heckscher, E.S., Zarin, A.A., Faumont, S., Clark, M.Q., Manning, L., Fushiki, A., et al. (2015). Even-Skipped^+^ Interneurons Are Core Components of a Sensorimotor Circuit that Maintains Left-Right Symmetric Muscle Contraction Amplitude. Neuron 88(2), 314–329. doi: 10.1016/j.neuron.2015.09.009.

Hoang, B., and Chiba, A. (2001). Single-cell analysis of *Drosophila* larval neuromuscular synapses. Dev Biol 229(1), 55–70. doi: 10.1006/dbio.2000.9983.

Hoff, M., Balfanz, S., Ehling, P., Gensch, T., and Baumann, A. (2011). A single amino acid residue controls Ca^2+^ signaling by an octopamine receptor from *Drosophila melanogaster*. FASEB J 25(7), 2484–2491. doi: 10.1096/fj.11-180703.

Hooper, J.E. (1986). Homeotic gene function in the muscles of *Drosophila* larvae. The EMBO Journal 5(9), 2321–3229. doi: 10.1002/j.1460-2075.1986.tb04500.x.

Hoyer, S.C., Eckart, A., Herrel, A., Zars, T., Fischer, S.A., Hardie, S.L., et al. (2008). Octopamine in male aggression of *Drosophila*. Curr Biol 18(3), 159–167. doi: 10.1016/j.cub.2007.12.052.

Hu, C., Petersen, M., Hoyer, N., Spitzweck, B., Tenedini, F., Wang, D., et al. (2017). Sensory integration and neuromodulatory feedback facilitate *Drosophila* mechanonociceptive behavior. Nat Neurosci 20(8), 1085–1095. doi: 10.1038/nn.4580.

Huang, Y., Ainsley, J.A., Reijmers, L.G., and Jackson, F.R. (2013). Translational profiling of clock cells reveals circadianly synchronized protein synthesis. PLoS Biol 11(11), e1001703. doi: 10.1371/journal.pbio.1001703.

Hückesfeld, S., Schlegel, P., Miroschnikow, A., Schoofs, A., Zinke, I., Haubrich, A.N., et al. (2021). Unveiling the sensory and interneuronal pathways of the neuroendocrine connectome in *Drosophila*. eLife 10(:e65745). doi: 10.7554/eLife.65745.

Huetteroth, W., Perisse, E., Lin, S., Klappenbach, M., Burke, C., and Waddell, S. (2015). Sweet taste and nutrient value subdivide rewarding dopaminergic neurons in *Drosophila*. Curr Biol 25(6), 751–758. doi: 10.1016/j.cub.2015.01.036.

Jan, Y.N., and Jan, L.Y. (1993). ““The peripheral nervous system”,” in The Development of Drosophila melanogaster, ed. M. Bate. Cold Spring Harbor Laboratory Press), 1207–1244.

Jetti, S.K., Crane, A.B., Akbergenova, Y., Aponte-Santiago, N.A., Cunningham, K.L., Whittaker, C.A., et al. (2023). Molecular logic of synaptic diversity between *Drosophila* tonic and phasic motoneurons. Neuron 111(22), 3554–3569 e3557. doi: 10.1016/j.neuron.2023.07.019.

Johnson, E., Ringo, J., and Dowse, H. (1997). Modulation of Drosophila heartbeat by neurotransmitters. J Comp Physiol B 167, 89–97. doi: 10.1007/s003600050051.

Kim, D., Alvarez, M., Lechuga, L.M., and Louis, M. (2017). Species-specific modulation of food-search behavior by respiration and chemosensation in Drosophila larvae. Elife 6. doi: 10.7554/eLife.27057.

Kim, Y.C., Lee, H.G., Lim, J., and Han, K.A. (2013). Appetitive learning requires the alpha1-like octopamine receptor OAMB in the *Drosophila* mushroom body neurons. J Neurosci 33(4), 1672–1677. doi: 10.1523/JNEUROSCI.3042-12.2013.

Kondo, S., Takahashi, T., Yamagata, N., Imanishi, Y., Katow, H., Hiramatsu, S., et al. (2020). Neurochemical Organization of the *Drosophila* Brain Visualized by Endogenously Tagged Neurotransmitter Receptors. Cell Rep 30(1), 284–297 e285. doi: 10.1016/j.celrep.2019.12.018.

Koon, A.C., Ashley, J., Barria, R., DasGupta, S., Brain, R., Waddell, S., et al. (2011). Autoregulatory and paracrine control of synaptic and behavioral plasticity by octopaminergic signaling. Nat Neurosci 14(2), 190–199. doi: 10.1038/nn.2716.

Koon, A.C., and Budnik, V. (2012). Inhibitory control of synaptic and behavioral plasticity by octopaminergic signaling. J Neurosci 32(18), 6312–6322. doi: 10.1523/JNEUROSCI.6517-11.2012.

Kula-Eversole, E., Nagoshi, E., Shang, Y., Rodriguez, J., Allada, R., and Rosbash, M. (2010). Surprising gene expression patterns within and between PDF-containing circadian neurons in Drosophila. Proc Natl Acad Sci U S A 107(30), 13497–13502. doi: 10.1073/pnas.1002081107.

Kutsukake, M., Komatsu, A., Yamamoto, D., and Ishiwa-Chigusa, S. (2000). A tyramine receptor gene mutation causes a defective olfactory behavior in *Drosophila melanogaster*. Gene 245(1), 31–42. doi: 10.1016/S0378-1119(99)00569-7.

Larkin, A., Marygold, S.J., Antonazzo, G., Attrill, H., Dos Santos, G., Garapati, P.V., et al. (2021). FlyBase: updates to the *Drosophila melanogaster* knowledge base. Nucleic Acids Res 49(D1), D899–D907. doi: 10.1093/nar/gkaa1026.

LeDue, E.E., Mann, K., Koch, E., Chu, B., Dakin, R., and Gordon, M.D. (2016). Starvation-Induced Depotentiation of Bitter Taste in Drosophila. Curr Biol 26(21), 2854–2861. doi: 10.1016/j.cub.2016.08.028.

Lee, H.-G., Seong, C.-S., Kim, Y.-C., Davis, R.L., and Han, K.-A. (2003). Octopamine receptor OAMB is required for ovulation in *Drosophila melanogaster*. Dev Biol 264(1), 179–190. doi: 10.1016/j.ydbio.2003.07.018.

Lee, H.G., Rohila, S., and Han, K.A. (2009). The octopamine receptor OAMB mediates ovulation via Ca2+/calmodulin-dependent protein kinase II in the *Drosophila* oviduct epithelium. PLoS One 4(3), e4716. doi: 10.1371/journal.pone.0004716.

Li, K., Tsukasa, Y., Kurio, M., Maeta, K., Tsumadori, A., Baba, S., et al. (2023). Belly roll, a GPI-anchored Ly6 protein, regulates *Drosophila melanogaster* escape behaviors by modulating the excitability of nociceptive peptidergic interneurons. eLife 12, e83856. doi: 10.7554/eLife.83856.

Li, Y., Fink, C., El-Kholy, S., and Roeder, T. (2015). The octopamine receptor octß2R is essential for ovulation and fertilization in the fruit fly *Drosophila melanogaster*. Arch Insect Biochem Physiol 88(3), 168–178. doi: 10.1002/arch.21211.

Li, Y., Hoffmann, J., Li, Y., Stephano, F., Bruchhaus, I., Fink, C., et al. (2016). Octopamine controls starvation resistance, life span and metabolic traits in Drosophila. Sci Rep 6, 35359. doi: 10.1038/srep35359.

Lim, J., Sabandal, P.R., Fernandez, A., Sabandal, J.M., Lee, H.G., Evans, P., et al. (2014). The octopamine receptor Octbeta2R regulates ovulation in *Drosophila melanogaster*. PLoS One 9(8), e104441. doi: 10.1371/journal.pone.0104441.

Liu, Y., Hasegawa, E., Nose, A., Zwart, M.F., and Kohsaka, H. (2023). Synchronous multi-segmental activity between metachronal waves controls locomotion speed in Drosophila larvae. Elife 12. doi: 10.7554/eLife.83328.

Louis, M., Huber, T., Benton, R., Sakmar, T.P., and Vosshall, L.B. (2008). Bilateral olfactory sensory input enhances chemotaxis behavior. Nat Neurosci 11(2), 187–199. doi: 10.1038/nn2031.

Lu, S., Qian, C.S., and Grueber, W.B. (2024). Mechanisms of gas sensing by internal sensory neurons in *Drosophila* larvae. bioRxiv [Preprint]. Available at: https://www.biorxiv.org/content/10.1101/2024.01.20.576342v1 (Accessed September 26, 2025)

Luo, J., Lushchak, O.V., Goergen, P., Williams, M.J., and Nassel, D.R. (2014). *Drosophila* insulin-producing cells are differentially modulated by serotonin and octopamine receptors and affect social behavior. PLoS One 9(6), e99732. doi: 10.1371/journal.pone.0099732.

Ma, Z., Stork, T., Bergles, D.E., and Freeman, M.R. (2016). Neuromodulators signal through astrocytes to alter neural circuit activity and behaviour. Nature 539(7629), 428–432. doi: 10.1038/nature20145.

Maiellaro, I., Lohse, M.J., Kittel, R.J., and Calebiro, D. (2016). cAMP Signals in *Drosophila* Motor Neurons Are Confined to Single Synaptic Boutons. Cell Rep 17(5), 1238–1246. doi: 10.1016/j.celrep.2016.09.090.

Manjila, S.B., Kuruvilla, M., Ferveur, J.F., Sane, S.P., and Hasan, G. (2019). Extended Flight Bouts Require Disinhibition from GABAergic Mushroom Body Neurons. Curr Biol 29(2), 283–293 e285. doi: 10.1016/j.cub.2018.11.070.

Maqueira, B., Chatwin, H., and Evans, P.D. (2005). Identification and characterization of a novel family of *Drosophila* beta-adrenergic-like octopamine G-protein coupled receptors. J Neurochem 94(2), 547–560. doi: 10.1111/j.1471-4159.2005.03251.x.

McKinney, H.M., Sherer, L.M., Williams, J.L., Certel, S.J., and Stowers, R.S. (2020). Characterization of *Drosophila* octopamine receptor neuronal expression using MiMIC-converted Gal4 lines. J Comp Neurol 528(13), 2174–2194. doi: 10.1002/cne.24883.

Mezheritskiy, M.I., Vorontsov, D.D., Dyakonova, V.E., and Zakharov, I.S. (2024). Behavioral Functions of Octopamine in Adult Insects under Stressful Conditions. Biology Bulletin Reviews 14(5), 535–547. doi: 10.1134/s2079086424700014.

Monastirioti, M. (2003). Distinct octopamine cell population residing in the CNS abdominal ganglion controls ovulation in *Drosophila melanogaster*. Dev Biol 264, 38–49. doi: 10.1016/S0012-1606(03)00459-7.

Monastirioti, M., Gorczyca, M., Rapus, J., Eckert, M., White, K., and Budnik, V. (1995). Octopamine immunoreactivity in the fruit fly *Drosophila melanogaster*. J Comp Neurol 356(2), 275–287. doi: 10.1002/cne.903560210.

Monastirioti, M., Linn, C.E., and White, K. (1996). Characterization of *Drosophila* Tyramine B-Hydroxylase gene and isolation of mutant flies lacking Octopamine. J Neurosci 16(12), 3900–3911. doi: 10.1523/JNEUROSCI.16-12-03900.1996.

Nagaya, Y., Kutsukake, M., Chigusa, S.I., and Komatsu, A. (2002). A trace amine, tyramine, functions as a neuromodulator in *Drosophila melanogaster*. Neuroscience Letters 329(3), 324–328. doi: 10.1016/S0304-3940(02)00596-7.

Nakagawa, H., Maehara, S., Kume, K., Ohta, H., and Tomita, J. (2022). Biological functions of alpha2-adrenergic-like octopamine receptor in *Drosophila melanogaster*. Genes Brain Behav 21(6), e12807. doi: 10.1111/gbb.12807.

Ohhara, Y., Kayashima, Y., Hayashi, Y., Kobayashi, S., and Yamakawa-Kobayashi, K. (2012). Expression of β-adrenergic-like Octopamine Receptors during *Drosophila* Development. Zoological Science 29(2), 83–89. doi: 10.2108/zsj.29.83.

Ohhara, Y., Shimada-Niwa, Y., Niwa, R., Kayashima, Y., Hayashi, Y., Akagi, K., et al. (2015). Autocrine regulation of ecdysone synthesis by beta3-octopamine receptor in the prothoracic gland is essential for *Drosophila* metamorphosis. Proc. Natl. Acad. Sci. U.S.A. 112(5), 1452–1457. doi: 10.1073/pnas.1414966112.

Ohnishi, S. (1979). Relationship between larval feeding behavior and viability in *Drosophila melanogaster* and *D. simulans*. Behavior Genetics 9(2), 129–134. doi: 10.1007/BF01074332.

Öztürk-Çolak, A., Marygold, S.J., Antonazzo, G., Attrill, H., Goutte-Gattat, D., Jenkins, V.K., et al. (2024). FlyBase: updates to the *Drosophila* genes and genomes database. Genetics 227(1), iyad211. doi: 10.1093/genetics/iyad211.

Pauls, D., Blechschmidt, C., Frantzmann, F., El Jundi, B., and Selcho, M. (2018). A comprehensive anatomical map of the peripheral octopaminergic/tyraminergic system of *Drosophila melanogaster*. Sci Rep 8(1), 15314. doi: 10.1038/s41598-018-33686-3.

Pauls, D., Selcho, M., Gendre, N., Stocker, R.F., and Thum, A.S. (2010). *Drosophila* larvae establish appetitive olfactory memories via mushroom body neurons of embryonic origin. J Neurosci 30(32), 10655–10666. doi: 10.1523/JNEUROSCI.1281-10.2010.

Pauls, D., Selcho, M., Raderscheidt, J., Amatobi, K.M., Fekete, A., Krischke, M., et al. (2021). Endocrine signals fine-tune daily activity patterns in Drosophila. Curr Biol 31(18), 4076–4087 e4075. doi: 10.1016/j.cub.2021.07.002.

Pauls, D., von Essen, A., Lyutova, R., van Giesen, L., Rosner, R., Wegener, C., et al. (2015). Potency of transgenic effectors for neurogenetic manipulation in Drosophila larvae. Genetics 199(1), 25–37. doi: 10.1534/genetics.114.172023.

Qi, Y.X., Xu, G., Gu, G.X., Mao, F., Ye, G.Y., Liu, W., et al. (2017). A new *Drosophila* octopamine receptor responds to serotonin. Insect Biochem Mol Biol 90, 61–70. doi: 10.1016/j.ibmb.2017.09.010.

Reyes, M., Lee, Y.S., Ali, M.M., Sundaramurthi, P., Dhungana, N., Nguyen, A., et al. (2026). Octopamine regulates neural circuits in the mushroom body and central complex, influencing sleep and arousal. iScience 29(5), 115564. doi: 10.1016/j.isci.2026.115564.

Rezaval, C., Nojima, T., Neville, M.C., Lin, A.C., and Goodwin, S.F. (2014). Sexually dimorphic octopaminergic neurons modulate female postmating behaviors in *Drosophila*. Curr Biol 24(7), 725–730. doi: 10.1016/j.cub.2013.12.051.

Richter, V., Rist, A., Kislinger, G., Laumann, M., Schoofs, A., Miroschnikow, A., et al. (2023). Morphology and ultrastructure of external sense organs of *Drosophila* larvae. eLife 12, RP91155. doi: 10.7554/eLife.91155.1.

Rist, A., and Thum, A.S. (2017). A map of sensilla and neurons in the taste system of *Drosophila* larvae. J Comp Neurol 525(18), 3865–3889. doi: 10.1002/cne.24308.

Robb, S., Cheek, T.R., Hannan, F.L., Hall, L.M., Midgley, J.M., and Evans, P.D. (1994). Agonist-specific coupling of a cloned *Drosophila* octopamine/tyramine receptor to multiple second messenger systems. The EMBO Journal 13, 1325–1330. doi: 10.1002/j.1460-2075.1994.tb06385.x.

Roeder, T. (2005). Tyramine and octopamine: ruling behavior and metabolism. Annu Rev Entomol 50, 447–477. doi: 10.1146/annurev.ento.50.071803.130404.

Roeder, T. (2020). The control of metabolic traits by octopamine and tyramine in invertebrates. J Exp Biol 223(Pt 7). doi: 10.1242/jeb.194282.

Rohwedder, A., Pfitzenmaier, J.E., Ramsperger, N., Apostolopoulou, A.A., Widmann, A., and Thum, A.S. (2012). Nutritional value-dependent and nutritional value-independent effects on *Drosophila melanogaster* larval behavior. Chem Senses 37(8), 711–721. doi: 10.1093/chemse/bjs055.

Rulifson, E.J., Kim, S.K., and Nusse, R. (2002). Ablation of Insulin-Producing Neurons in Flies: Growth and Diabetic Phenotypes. Science 296(5570), 1118–1120. doi: 10.1126/science.1070058.

Sabandal, J.M., Sabandal, P.R., Kim, Y.C., and Han, K.A. (2020). Concerted Actions of Octopamine and Dopamine Receptors Drive Olfactory Learning. J Neurosci 40(21), 4240–4250. doi: 10.1523/JNEUROSCI.1756-19.2020.

Saraswati, S., Fox, L.E., Soll, D.R., and Wu, C.F. (2004). Tyramine and octopamine have opposite effects on the locomotion of *Drosophila* larvae. J Neurobiol 58(4), 425–441. doi: 10.1002/neu.10298.

Saudou, F., Amlaiky, N., Plassat, J.-L., Borrelli, E., and Hen, R. (1990). Cloning and characterization of a *Drosophila* tyramine receptor. The EMBO Journal 9(11), 3611–3617.

Sayin, S., De Backer, J.F., Siju, K.P., Wosniack, M.E., Lewis, L.P., Frisch, L.M., et al. (2019). A Neural Circuit Arbitrates between Persistence and Withdrawal in Hungry Drosophila. Neuron 104(3), 544–558 e546. doi: 10.1016/j.neuron.2019.07.028.

Schleyer, M., Saumweber, T., Nahrendorf, W., Fischer, B., von Alpen, D., Pauls, D., et al. (2011). A behavior-based circuit model of how outcome expectations organize learned behavior in larval *Drosophila*. Learn Mem 18(10), 639–653. doi: 10.1101/lm.2163411.

Schneider, A., Ruppert, M., Hendrich, O., Giang, T., Ogueta, M., Hampel, S., et al. (2012). Neuronal basis of innate olfactory attraction to ethanol in *Drosophila*. PLoS One 7(12), e52007. doi: 10.1371/journal.pone.0052007.

Schoofs, A., Niederegger, S., van Ooyen, A., Heinzel, H.G., and Spiess, R. (2010). The brain can eat: establishing the existence of a central pattern generator for feeding in third instar larvae of *Drosophila virilis* and *Drosophila melanogaster*. J Insect Physiol 56(7), 695–705. doi: 10.1016/j.jinsphys.2009.12.008.

Schretter, C.E., Vielmetter, J., Bartos, I., Marka, Z., Marka, S., Argade, S., et al. (2018). A gut microbial factor modulates locomotor behaviour in Drosophila. Nature 563(7731), 402–406. doi: 10.1038/s41586-018-0634-9.

Schützler, N., Girwert, C., Hugli, I., Mohana, G., Roignant, J.Y., Ryglewski, S., et al. (2019). Tyramine action on motoneuron excitability and adaptable tyramine/octopamine ratios adjust Drosophila locomotion to nutritional state. Proc Natl Acad Sci U S A 116(9), 3805–3810. doi: 10.1073/pnas.1813554116.

Schwaerzel, M., Monastirioti, M., Scholz, H., Friggi-Grelin, F., Birman, S., and Heisenberg, M. (2003). Dopamine and Octopamine Differentiate between Aversive and Appetitive Olfactory Memories in *Drosophila*. J Neurosci 23(33), 10495–10502. doi: 10.1523/JNEUROSCI.23-33-10495.2003.

Selcho, M. (2024). Octopamine in the mushroom body circuitry for learning and memory. Learn Mem 31(5). doi: 10.1101/lm.053839.123.

Selcho, M., Pauls, D., El Jundi, B., Stocker, R.F., and Thum, A.S. (2012). The role of octopamine and tyramine in *Drosophila* larval locomotion. J Comp Neurol 520(16), 3764–3785. doi: 10.1002/cne.23152.

Selcho, M., Pauls, D., Han, K.A., Stocker, R.F., and Thum, A.S. (2009). The role of dopamine in *Drosophila* larval classical olfactory conditioning. PLoS One 4(6), e5897. doi: 10.1371/journal.pone.0005897.

Selcho, M., Pauls, D., Huser, A., Stocker, R.F., and Thum, A.S. (2014). Characterization of the octopaminergic and tyraminergic neurons in the central brain of *Drosophila* larvae. J Comp Neurol 522(15), 3485–3500. doi: 10.1002/cne.23616.

Shang, Y., Haynes, P., Pirez, N., Harrington, K.I., Guo, F., Pollack, J., et al. (2011). Imaging analysis of clock neurons reveals light buffers the wake-promoting effect of dopamine. Nat Neurosci 14(7), 889–895. doi: 10.1038/nn.2860.

Stowers, R.S. (2011). An efficient method for recombineering GAL4 and QF drivers. Fly 5(4), 371–378. doi: 10.4161/fly.5.4.17560.

Sujkowski, A., Ramesh, D., Brockmann, A., and Wessells, R. (2017). Octopamine Drives Endurance Exercise Adaptations in Drosophila. Cell Rep 21(7), 1809–1823. doi: 10.1016/j.celrep.2017.10.065.

Sujkowski, A., and Wessells, R. (2018). Using Drosophila to Understand Biochemical and Behavioral Responses to Exercise. Exerc Sport Sci Rev 46(2), 112–120. doi: 10.1249/JES.0000000000000139.

Suver, M.P., Mamiya, A., and Dickinson, M.H. (2012). Octopamine neurons mediate flight-induced modulation of visual processing in Drosophila. Curr Biol 22(24), 2294–2302. doi: 10.1016/j.cub.2012.10.034.

Szuperak, M., Churgin, M.A., Borja, A.J., Raizen, D.M., Fang-Yen, C., and Kayser, M.S. (2018). A sleep state in Drosophila larvae required for neural stem cell proliferation. Elife 7. doi: 10.7554/eLife.33220.

Tomancak, P., Beaton, A., Weiszmann, R., Kwan, E., Shu, S.Q., Lewis, S.E., et al. (2002). Systematic determination of patterns of gene expression during *Drosophila* embryogenesis. Genome Biol 3(research0088.1). doi: 10.1186/gb-2002-3-12-research0088.

Tomancak, P., Berman, B.P., Beaton, A., Weiszmann, R., Kwan, E., Hartenstein, V., et al. (2007). Global analysis of patterns of gene expression during *Drosophila* embryogenesis. Genome Biol 8(7), R145. doi: 10.1186/gb-2007-8-7-r145.

Tsubouchi, A., Caldwell, J.C., and Tracey, W.D. (2012). Dendritic filopodia, Ripped Pocket, NOMPC, and NMDARs contribute to the sense of touch in *Drosophila* larvae. Curr Biol 22(22), 2124–2134. doi: 10.1016/j.cub.2012.09.019.

van Breugel, F., Suver, M.P., and Dickinson, M.H. (2014). Octopaminergic modulation of the visual flight speed regulator of Drosophila. J Exp Biol 217(Pt 10), 1737–1744. doi: 10.1242/jeb.098665.

Venken, K.J., Schulze, K.L., Haelterman, N.A., Pan, H., He, Y., Evans-Holm, M., et al. (2011). MiMIC: a highly versatile transposon insertion resource for engineering *Drosophila melanogaster* genes. Nat Methods 8(9), 737–743. doi: 10.1038/nmeth.1662.

Wang, J., Chitsaz, F., Derbyshire, M.K., Gonzales, N.R., Gwadz, M., Lu, S., et al. (2023). The conserved domain database in 2023. Nucleic Acids Res 51(D1), D384–D388. doi: 10.1093/nar/gkac1096.

Wang, Q.P., Lin, Y.Q., Zhang, L., Wilson, Y.A., Oyston, L.J., Cotterell, J., et al. (2016). Sucralose Promotes Food Intake through NPY and a Neuronal Fasting Response. Cell Metab 24(1), 75–90. doi: 10.1016/j.cmet.2016.06.010.

Weber, D., Richter, V., Rohwedder, A., Großjohann, A., and Thum, A.S. (2023a). The Analysis of Aversive Olfactory-Taste Learning and Memory in Drosophila Larvae. Cold Spring Harb Protoc 2023(3), 108050-pdb prot. doi: 10.1101/pdb.prot108050.

Weber, D., Richter, V., Rohwedder, A., Großjohann, A., and Thum, A.S. (2023b). Learning and Memory in Drosophila Larvae. Cold Spring Harb Protoc 2023(3), 107863-pdb top. doi: 10.1101/pdb.top107863.

Wipfler, B., Schneeberg, K., Loffler, A., Hunefeld, F., Meier, R., and Beutel, R.G. (2013). The skeletomuscular system of the larva of *Drosophila melanogaster* (Drosophilidae, Diptera): a contribution to the morphology of a model organism. Arthropod Struct Dev 42(1), 47–68. doi: 10.1016/j.asd.2012.09.005.

Yang, C.H., Shih, M.F., Chang, C.C., Chiang, M.H., Shih, H.W., Tsai, Y.L., et al. (2016). Additive Expression of Consolidated Memory through *Drosophila* Mushroom Body Subsets. PLoS Genet 12(5), e1006061. doi: 10.1371/journal.pgen.1006061.

Yang, Z., Yu, Y., Zhang, V., Tian, Y., Qi, W., and Wang, L. (2015). Octopamine mediates starvation-induced hyperactivity in adult *Drosophila*. Proc. Natl. Acad. Sci. U.S.A. 112(16), 5219–5224. doi: 10.1073/pnas.1417838112.

Yarali, A., and Gerber, B. (2010). A Neurogenetic Dissociation between Punishment-, Reward-, and Relief-Learning in *Drosophila*. Front Behav Neurosci 4, 189. doi: 10.3389/fnbeh.2010.00189.

Ye, J., Coulouris, G., Zaretskaya, I., Cutcutache, I., Rozen, S., and Madden, T.L. (2012). Primer-BLAST: A tool to design target-specific primers for polymerase chain reaction. BMC Bioinformatics 13(1), 134. doi: 10.1186/1471-2105-13-134.

Youn, H., Kirkhart, C., Chia, J., and Scott, K. (2018). A subset of octopaminergic neurons that promotes feeding initiation in Drosophila melanogaster. PLoS One 13(6), e0198362. doi: 10.1371/journal.pone.0198362.

Yu, Y., Huang, R., Ye, J., Zhang, V., Wu, C., Cheng, G., et al. (2016). Regulation of starvation-induced hyperactivity by insulin and glucagon signaling in adult *Drosophila*. eLife 5. doi: 10.7554/eLife.15693.

Zarin, A.A., Mark, B., Cardona, A., Litwin-Kumar, A., and Doe, C.Q. (2019). A multilayer circuit architecture for the generation of distinct locomotor behaviors in *Drosophila*. eLife 8. doi: 10.7554/eLife.51781.

Zhang, T., Branch, A., and Shen, P. (2013). Octopamine-mediated circuit mechanism underlying controlled appetite for palatable food in Drosophila. Proc Natl Acad Sci U S A 110(38), 15431–15436. doi: 10.1073/pnas.1308816110.

Zhou, C., Huang, H., Kim, S.M., Lin, H., Meng, X., Han, K.A., et al. (2012). Molecular genetic analysis of sexual rejection: roles of octopamine and its receptor OAMB in *Drosophila* courtship conditioning. J Neurosci 32(41), 14281–14287. doi: 10.1523/JNEUROSCI.0517-12.2012.

Zhou, C., Rao, Y., and Rao, Y. (2008). A subset of octopaminergic neurons are important for *Drosophila* aggression. Nat Neurosci 11(9), 1059–1067. doi: 10.1038/nn.2164.

