## Supplementary Figures for "Octopamine receptors at a glance: from expression and anatomical maps to their role in development and behavior in the *Drosophila melanogaster* larva"

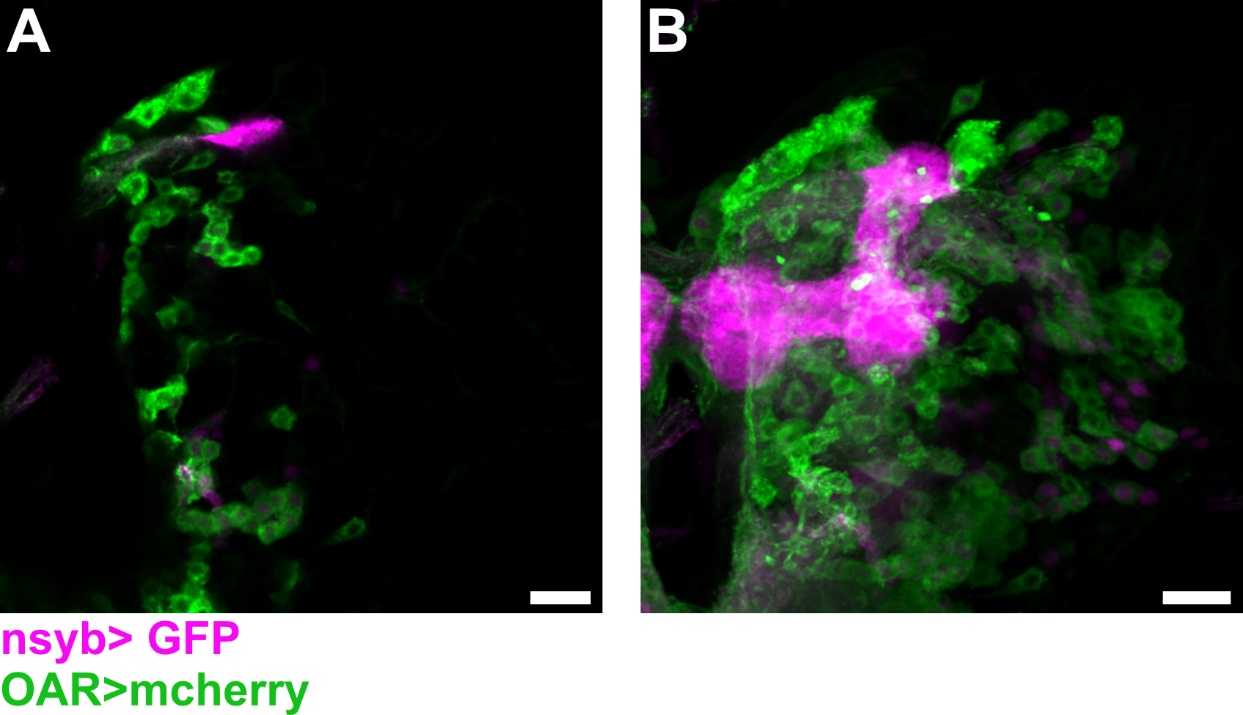

**Supplementary Figure 1: Cell cluster in superior medial protocerebrum with (A) somata in the pars intercerebralis and (B) projections along the midline of each hemisphere.** Using native fluorescence, the expression pattern of Octα2R (green via mcherry) and neuronal Synaptobrevin, as a reference channel (magenta via GFP), is shown. This cell cluster may correspond to insulin producing cells (compare Rulifson et al., 2002; Cao et al., 2014); however, further investigation is needed to confirm cell identity. Scale bar: 20 µm.

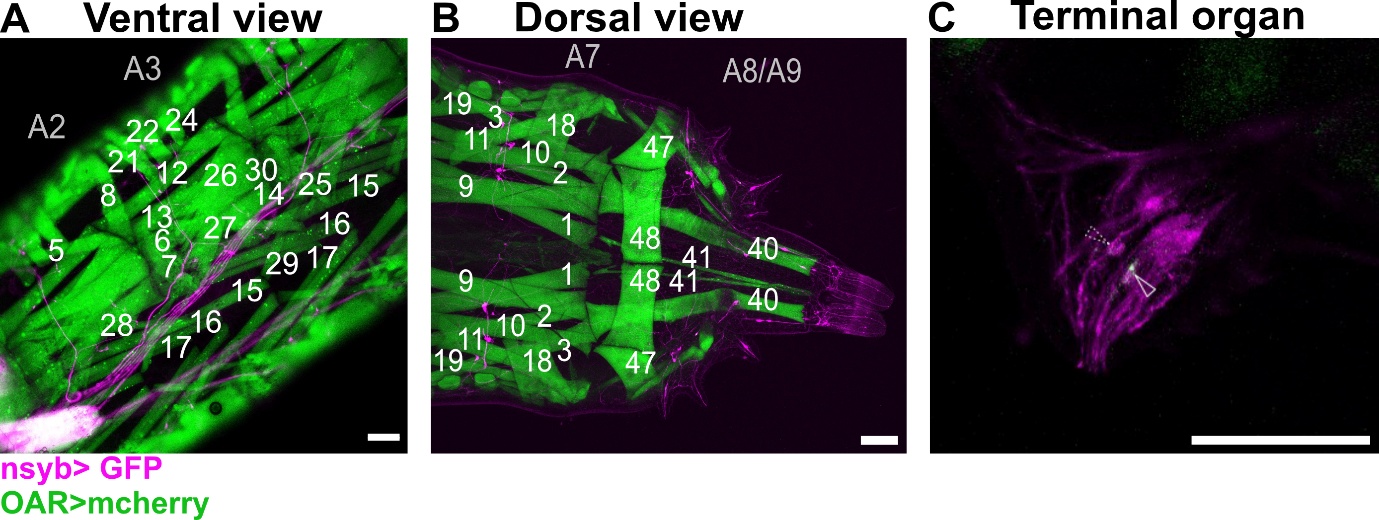

**Supplementary Figure 2: Trojan *Octβ2R* expression analysis throughout the body of 3^rd^ instar larva.** Using native fluorescence, the expression pattern of Octβ2R (green via mcherry) and neuronal Synaptobrevin, as a reference channel (magenta via GFP), is shown. **(A)** Ventral view of the larva shows ventral longitudinal (muscle 6, 7), ventral oblique (muscle 15, 16, 17) and ventral acute (muscle 29) muscles. **(B)** Dorsal view of the terminal shows dorsal transverse muscle 48. **(C)** Octβ2R signal was identified in at least one sensory neuron in the terminal organ (arrows). Additional muscles are labelled for orientation. A2/3/7/8/9: abdominal segment 2/3/7/8/9. Scale bar: 100 µm.

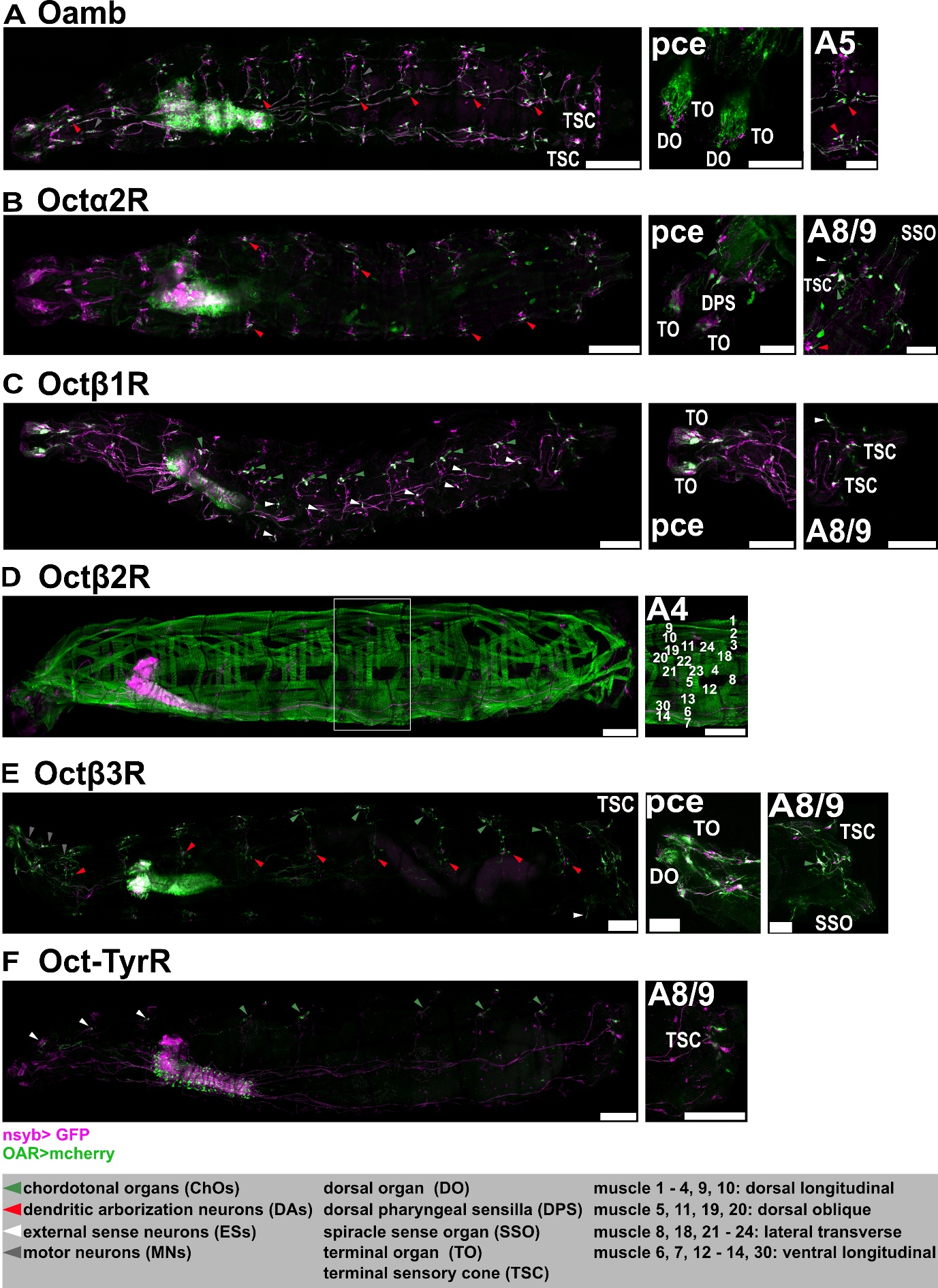

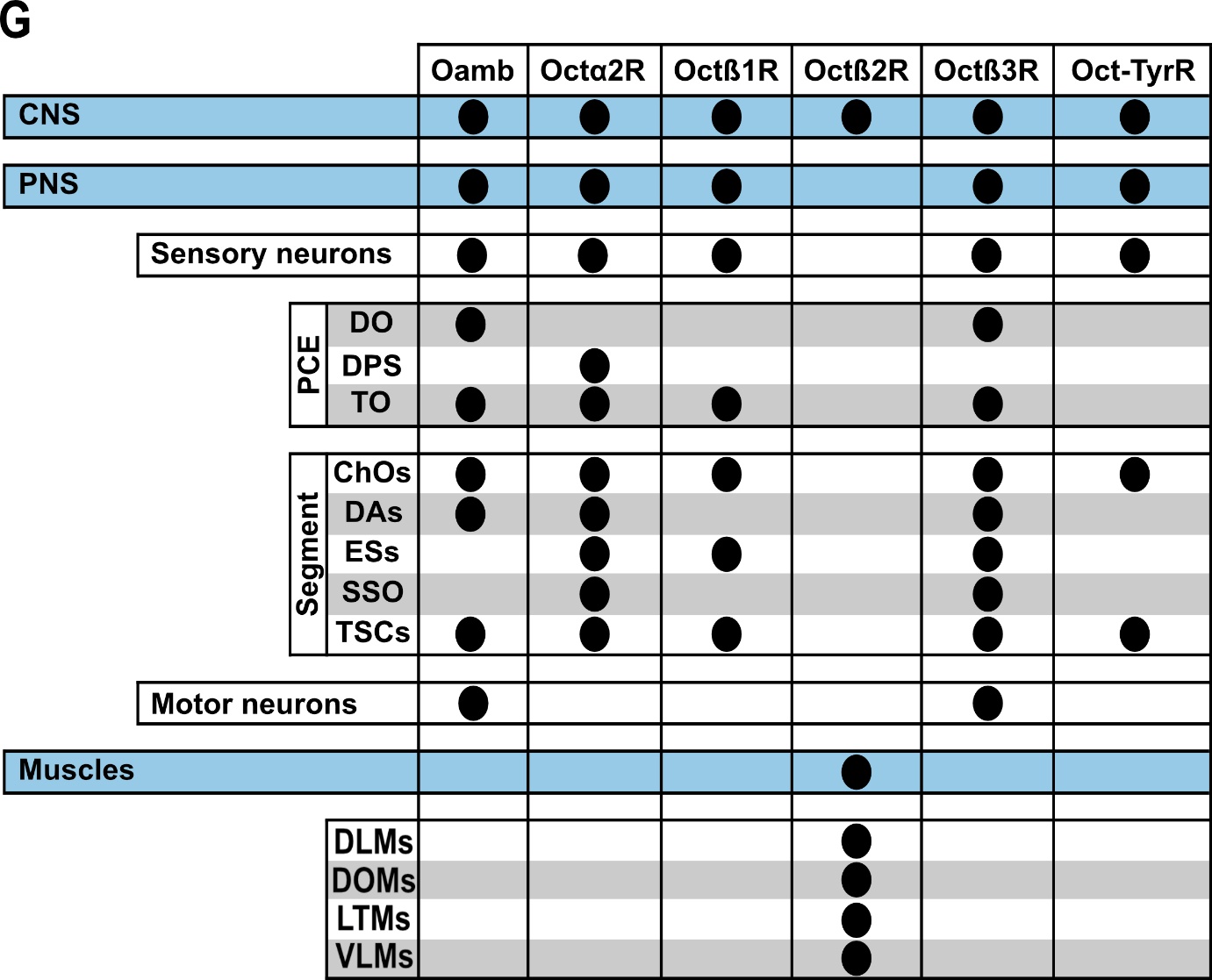

**Supplementary Figure 3: Trojan Exon octopamine receptor expression analysis throughout the body of 1^st^ instar larva.** **(A)-(F)** Using native fluorescence, the expression pattern of each octopamine receptor (green via mcherry) and neuronal Synaptobrevin, as a reference channel (magenta via GFP), is shown. **(G)** The table summarizes the expression locations in the 1^st^ instar larva. PCE: pseudocephalon; A4/5/8/9: abdominal segment 4/5/8/9; DLM: dorsal longitudinal muscle; DOM: dorsal oblique muscle; LTM: lateral transverse muscle; VLM: ventral longitudinal muscle. Scale bar: 100 µm.

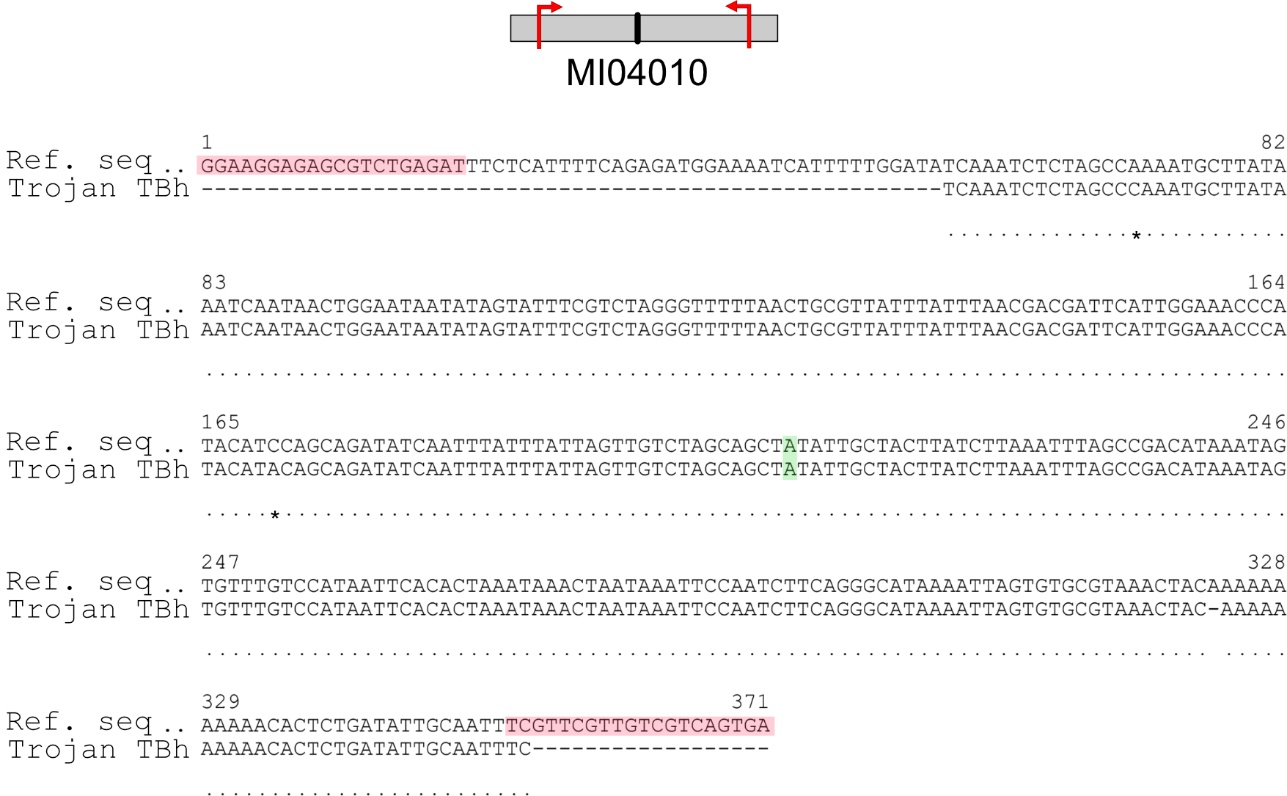
**Supplementary Figure 4: Sequencing result of amplified Trojan Exon insert location in Trojan *Tβh*.** The schematic shows the location of the primer according to the MI04010 docking site which was used to insert the Trojan Exon. One primer is upstream of the docking site, the second primer is downstream. Both are located in the gene region of *Tβh*. Below, the sequenced amplicon is compared with the gene region of *Tβh*. The sequenced amplicon has a size of 296 bp, a bit shorter than the observed product in the gel picture which is due to the sequencing process. The upper row is the reference sequence (Ref. seq), the lower row is the sequenced amplicon (Trojan TBh). Red: primer; green: MI04010 docking site; Dots show same bases, dashes show missing bases, asterisks show dissimilar bases.

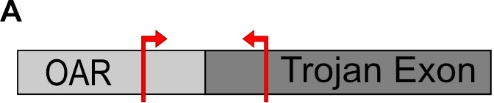

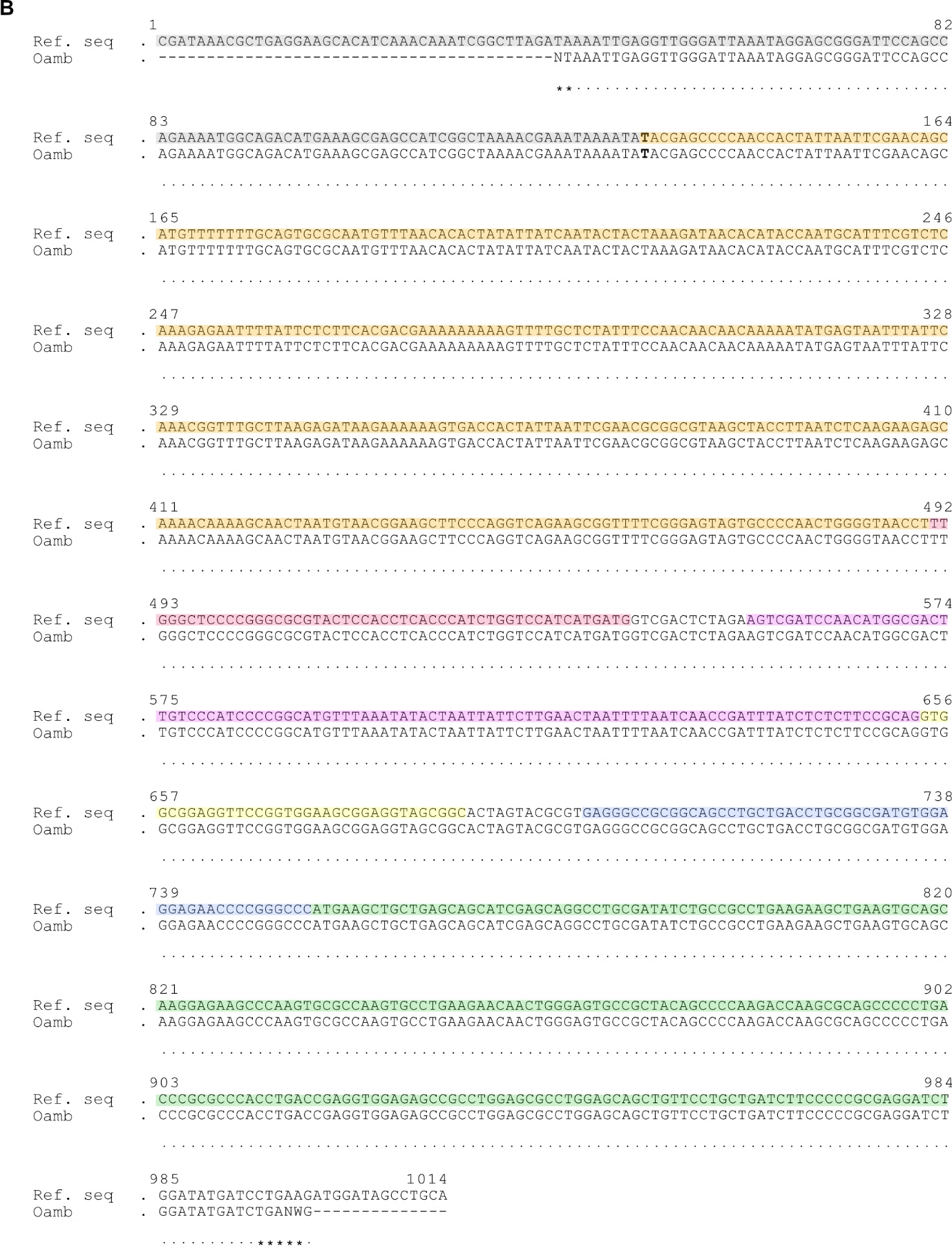

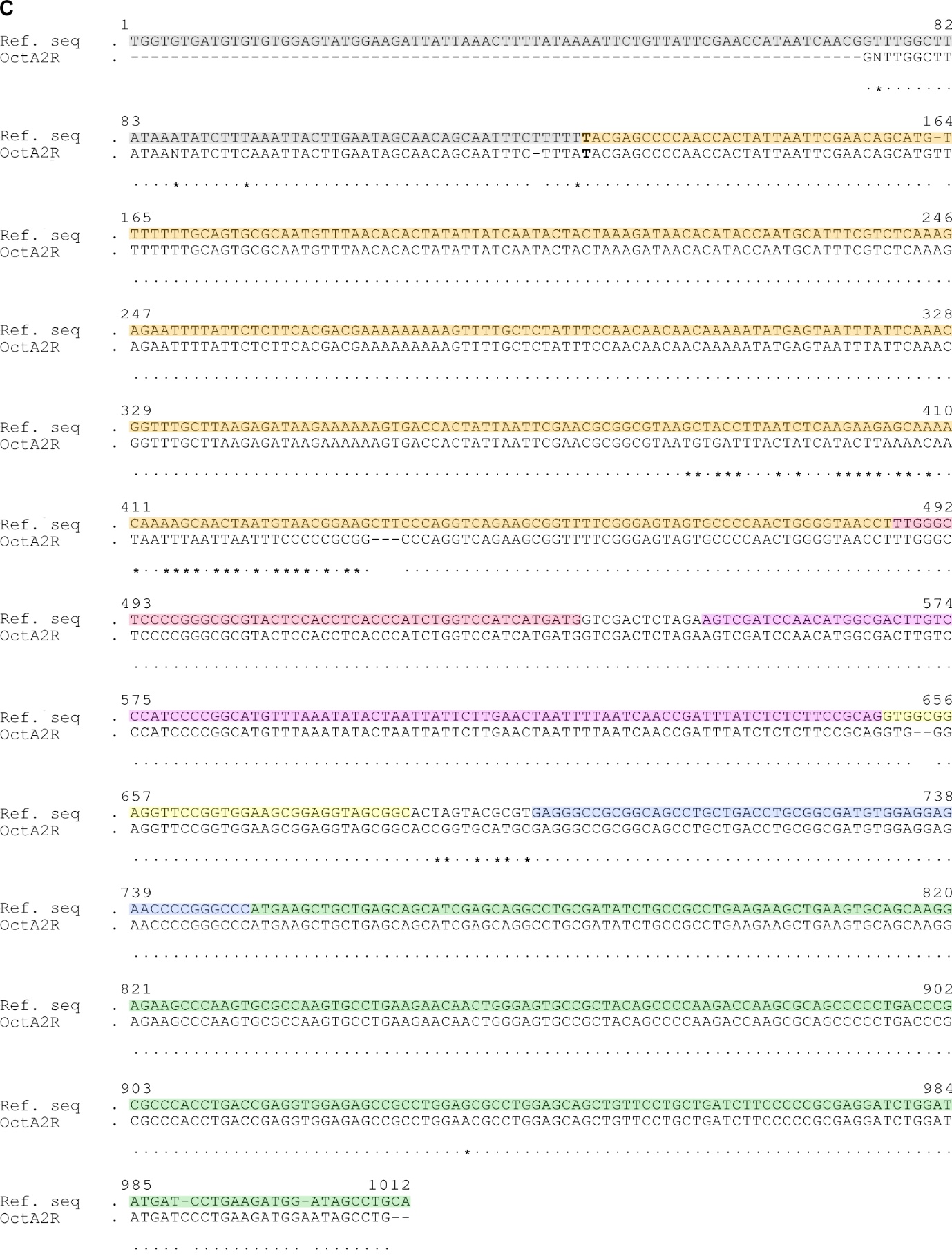

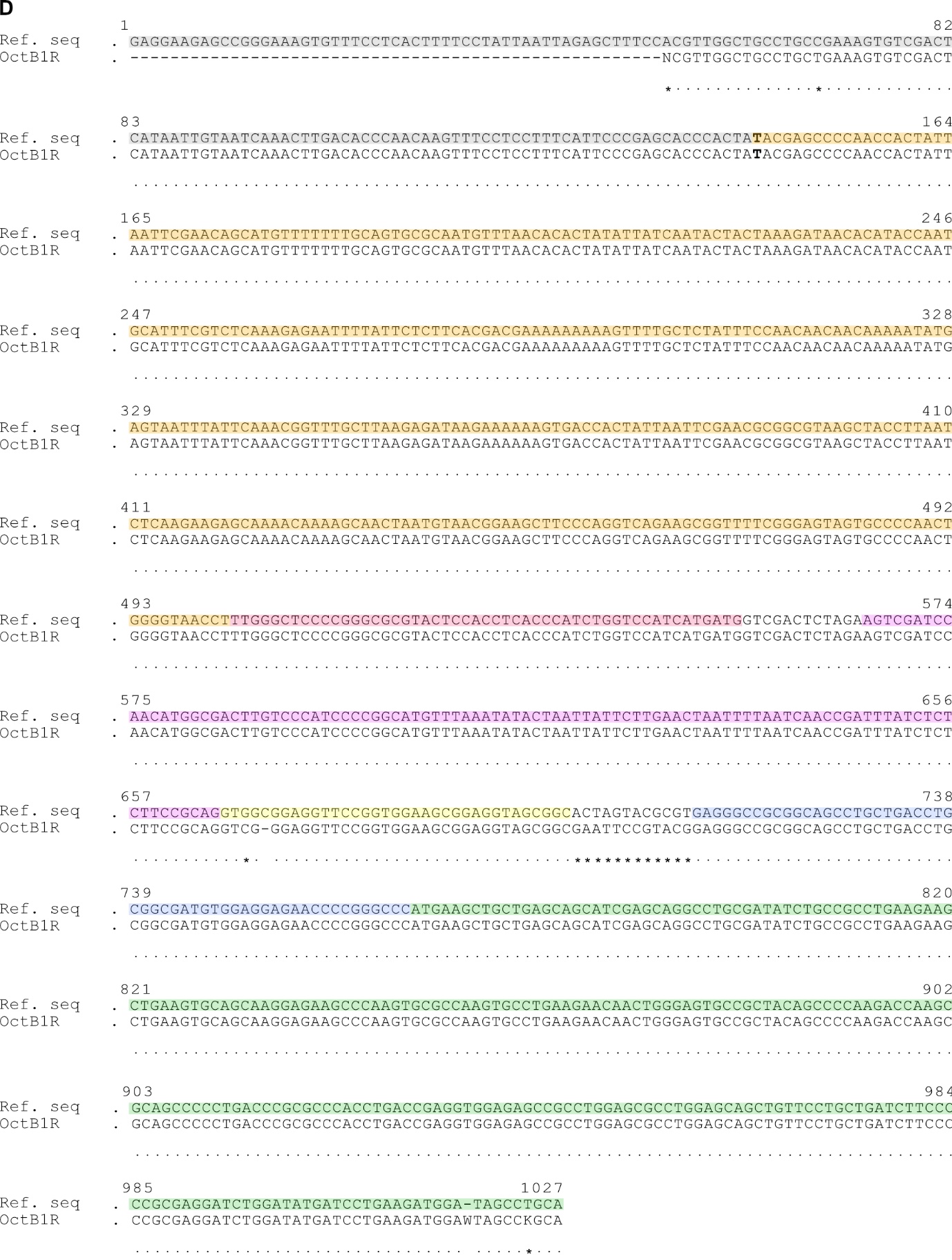

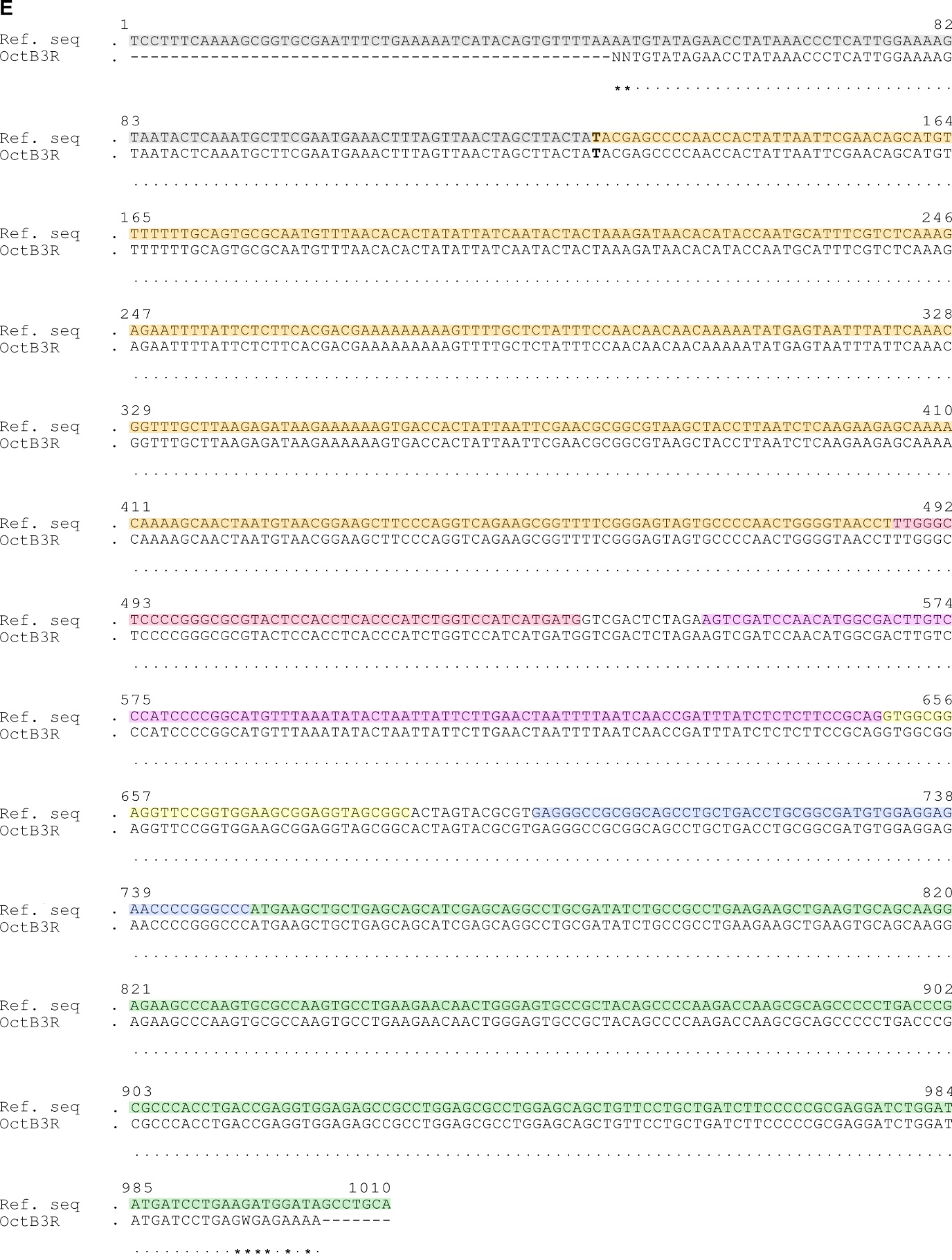

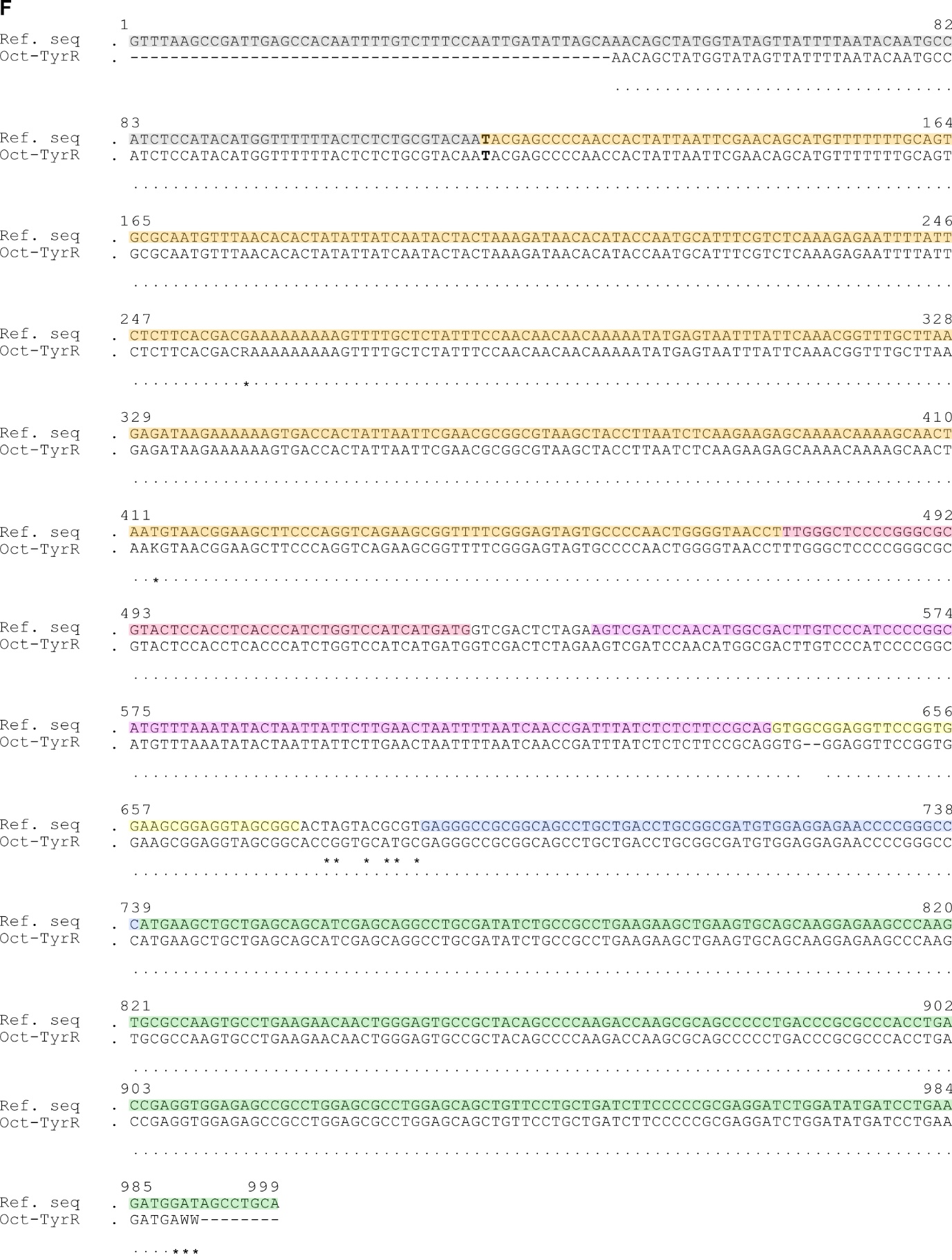

**Supplementary Figure 5: Sequencing results of amplified Trojan Exon insert location in Trojan *Oamb,*** ***Octα2R,*** ***Octβ1R,*** ***Octβ3R* and *Oct-TyrR*.** **(A)-(F)** Verification of the insertion site of the Trojan Exon: the sequencing results of the amplified products depicted in Figure 6B_I_ (*Oamb*), C_I_ (*Octα2R*), D_I_ (*Octβ1R*), F_I_ (*Octβ3R*) and G_I_ (*Oct-TyrR*) are shown. Each reference sequence (Ref. seq; upper row) starts and ends with the primer (red arrows in A) used during the PCR (a schematic is shown in A). For each octopamine receptor, the first primer is located in an Intron (grey), that is upstream of the Trojan Exon, followed by the *Minos* inverted repeat (orange) and the Trojan Exon itself (partially covered with the attB (red), splice acceptor (violet), linker (yellow), T2A (blue) and partial Gal4 (green)). The second primer is located in the Gal4 of the Trojan Exon; hence the Gal4 is sequenced only partially. The reference sequence shows only the region that is amplified using the selected primer, not the entire Intron and Trojan Exon. The lower row is the sequenced amplicon (Trojan Exon octopamine receptor mutant line). Grey: Intron; orange: *Minos* inverted repeat; red: attB; violet: splice acceptor; yellow: linker; blue: T2A; green: Gal4; Bold letters: MiMIC docking site; Dots show same bases, dashes show missing bases, asterisks show dissimilar bases; K: Guanine/Thymine; N: unclear nucleotide; R: Guanine/Adenine; W: Adenine/Thymine.

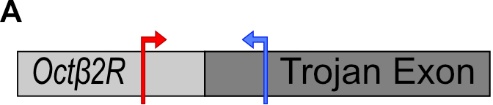

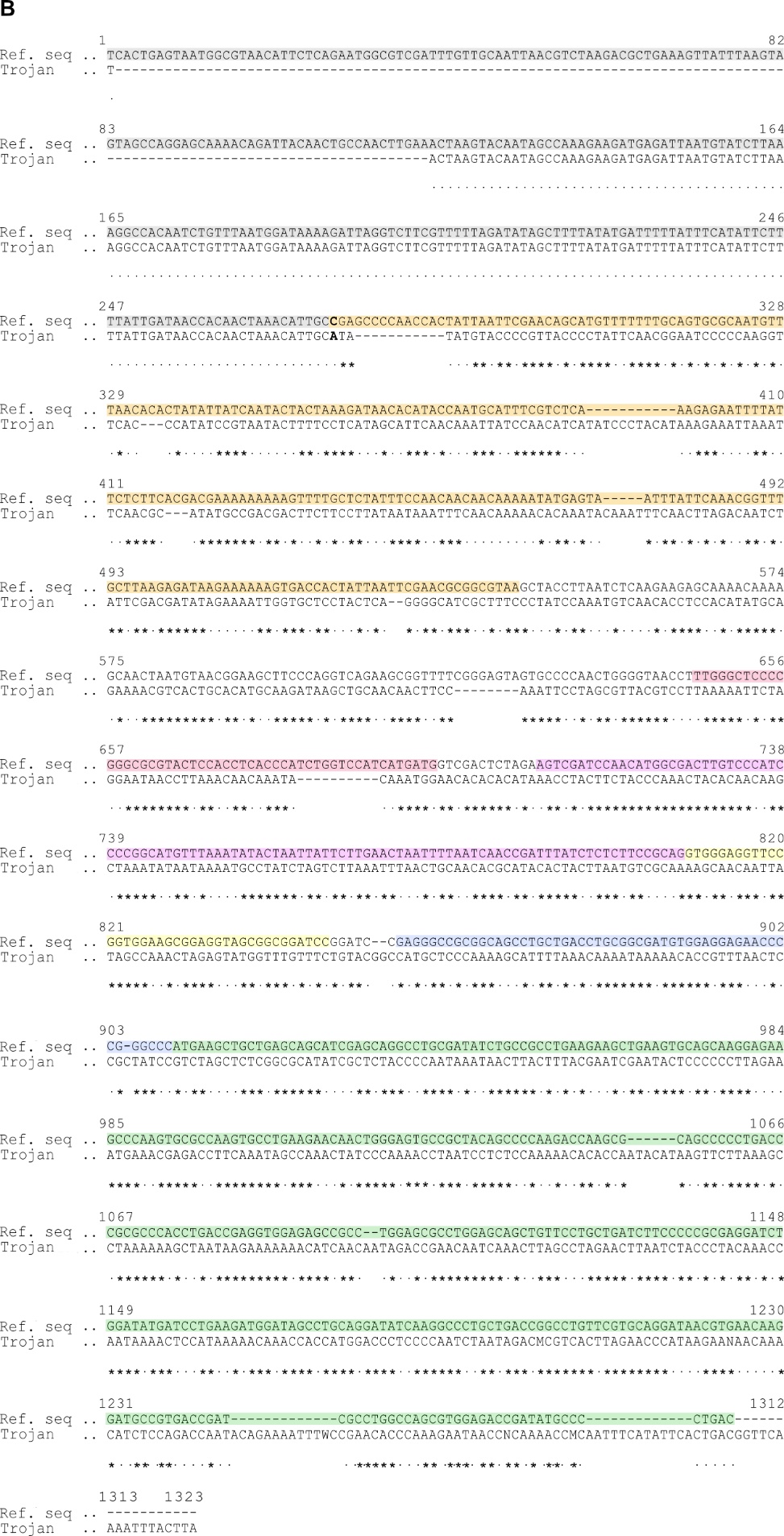

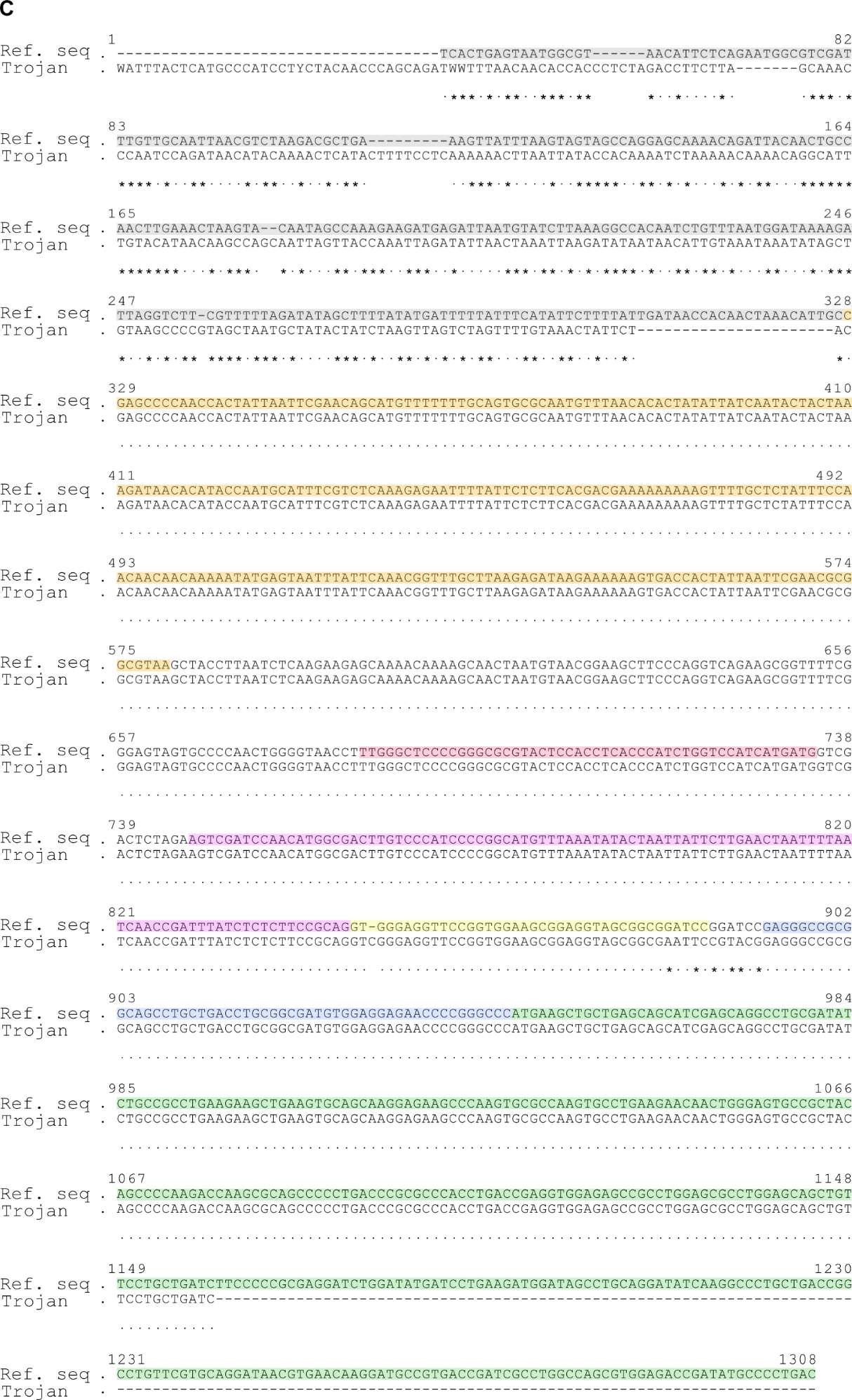

**Supplementary Figure 6: Sequencing result of 5´additional parts of Trojan Exon in Trojan *Octβ2R*.** During PCR analysis a larger fragment than expected was observed (expected: 1072 bp, observed: ~ 3000 bp). Here, the sequencing results of the amplified product depicted in Figure 6_I_ is shown. Due to the large size, the amplicon had to be sequenced in two steps as depicted in schematic **(A)**: first, with a primer located in the genomic region of *Octβ2R* (red arrow, sequencing result shown in B) and second, with a primer located in the Trojan Exon (blue arrow, sequencing result shown in C). For **(B)** and **(C)**, the upper row is the reference sequence (Ref. seq) consisting of Intron 6 (grey), that is upstream of the Trojan Exon, the *Minos* inverted repeat (orange) and the Trojan Exon itself (partially covered with the attB (red), splice acceptor (violet), linker (yellow), T2A (blue) and partial Gal4 (green)). The reference sequence shows only the region that is amplified using the selected primer, not the entire Intron 6 and Trojan Exon. The upper row is the reference sequence (Ref. seq), the lower row is the sequenced amplicon (Trojan). Grey: Intron 6; orange: *Minos* inverted repeat; red: attB; violet: splice acceptor; yellow: linker; blue: T2A; green: Gal4; Bold letters: MiMIC docking site; Dots show same bases, dashes show missing bases, asterisks show dissimilar bases; W: Adenine/Thymine.

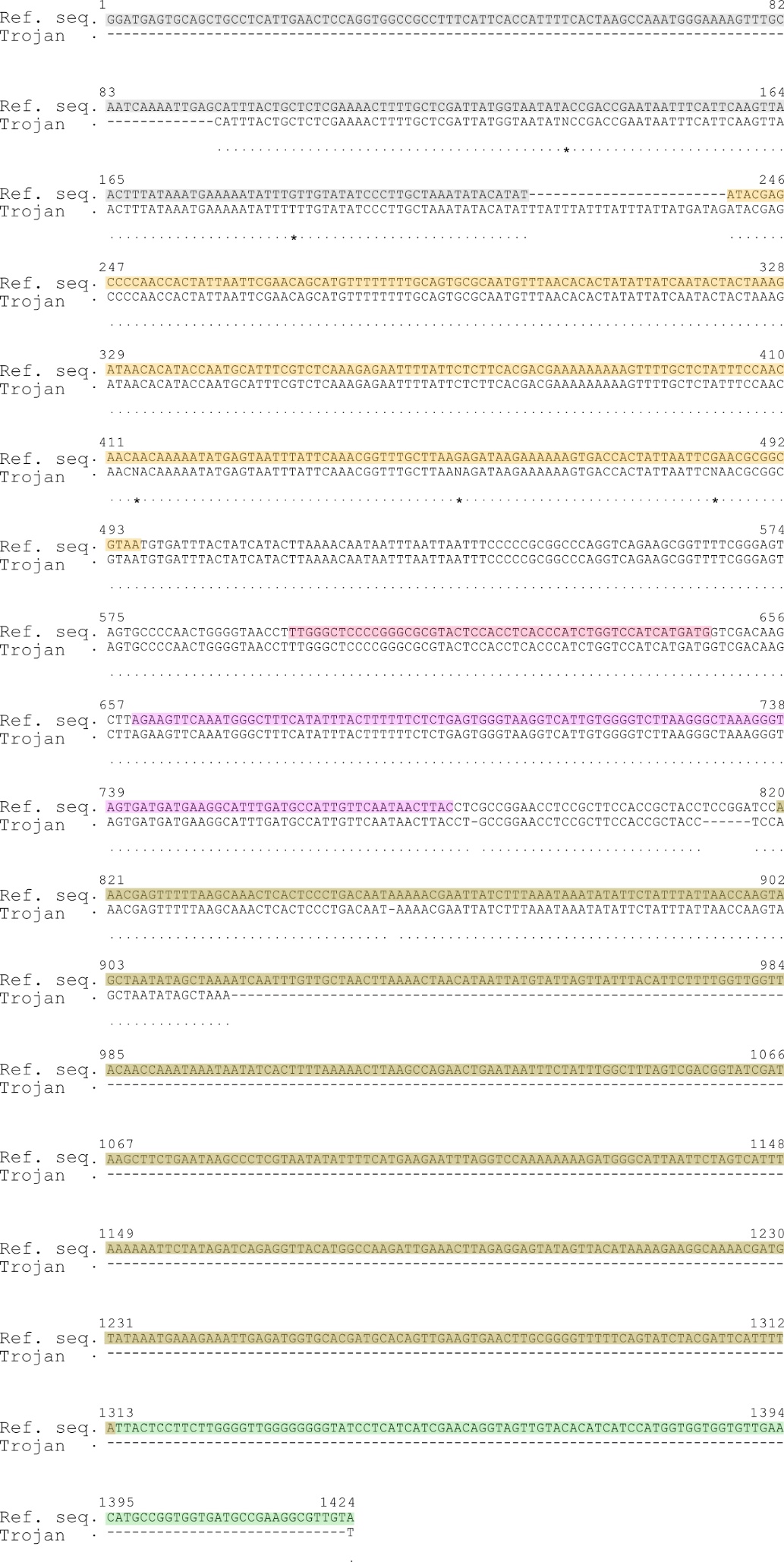

**Supplementary Figure 7: Sequencing result of 3´additional parts of Trojan Exon in Trojan *Octβ2R*.** During PCR analysis a larger fragment than expected was observed (expected: 1366 bp, observed: ~1500 bp). Here, the sequencing results of the amplified product depicted in Figure 4E_II_ is shown. For sequencing, a primer located in the *Octβ2R* genomic region was selected. The upper row is the reference sequence (Ref. seq) consisting of Intron 6 (grey), that is downstream of the Trojan Exon, the *Minos* inverted repeat (orange) and the Trojan Exon itself (partially covered with the attB (red), splice donor (violet), Hsp70 polyA (brown) and partial Gal4 (green)). The reference sequence shows only the region that is amplified using the selected primer, not the entire Intron 6 and Trojan Exon. The upper row is the reference sequence (Ref. seq), the lower row is the sequenced amplicon (Trojan). Grey: Intron 6; orange: *Minos* inverted repeat; red: attB; violet: splice donor; brown: Hsp70 polyA; green: Gal4; Dots show same bases, dashes show missing bases, asterisks show dissimilar bases; N: unclear nucleotide.

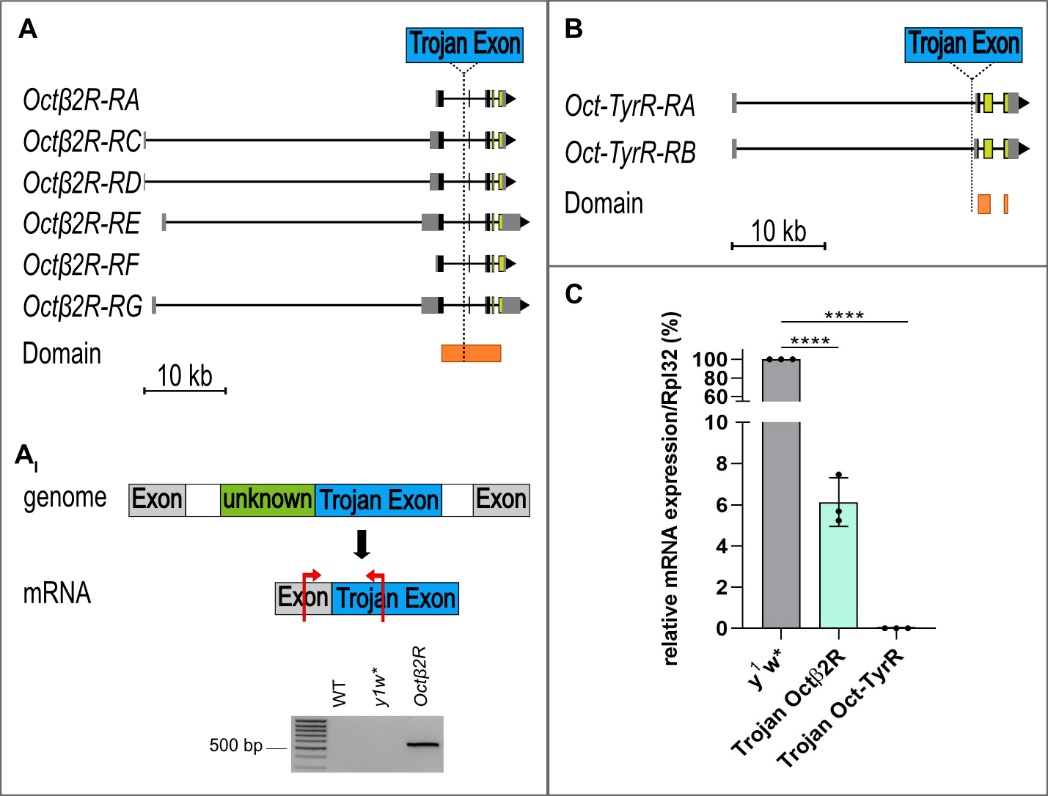

**Supplementary Figure 8: mRNA analysis of Trojan** ***Octβ2R* and *Oct-TyrR*.** Qualitative RT-PCR of Trojan *Octβ2R* showed that additional sequences that were found at the genomic level (Figure 6 E_I_ and E_II_) were not included in the Trojan *Octβ2R* mRNA. In **(A_I_)**, a schematic shows the location of the unknown sequence (green) right upstream of the Trojan Exon (blue) in the genome (schematic not to scale). Below, a schematic shows the location of the primer used during RT-PCR (red arrows). The size of the *Octβ2R* mRNA product in the Trojan *Octβ2R* was between 500 and 600 bp. The expected minimal size was 532 bp which includes the region of the upstream Exon and the Trojan Exon. Product size would have been bigger than the minimal 532 bp if additional sequences were included in the mRNA. Sequencing confirmed that no additional unknown sequences are included in the Trojan *Octβ2R* mRNA (Supplementary Figure 9). **(C)** Moreover, quantitative real-time RT-PCR showed that only a fraction (~6 %) of *Octβ2R* mRNA is present in the Trojan *Octβ2R* (normalized to housekeeping gene *Rpl32* and compared to *y^1^w**). In **(A)** the binding site of the TaqMan probe is depicted (yellow): spanning Exon 1/2 and 3, which is downstream of the inserted Trojan Exon in Intron 6. **(B)** and **(C)** Due to the insertion site of the Trojan Exon in *Oct-TyrR* (5´UTR), quantitative real-time RT-PCR was performed to measure transcript levels of *Oct-TyrR*. **(B)** shows the binding site of the TaqMan probe in the *Oct-TyrR* transcripts (yellow) which is downstream of the Trojan Exon insert location. **(C)** *Oct-TyrR* mRNA levels were undeterminable in the Trojan *Oct-TyrR* during quantitative real-time RT-PCR. Orange: predicted conserved domains for *Octβ2R* and *Oct-TyrR* (see Supplementary Table 5); **** = p < 0.0001.

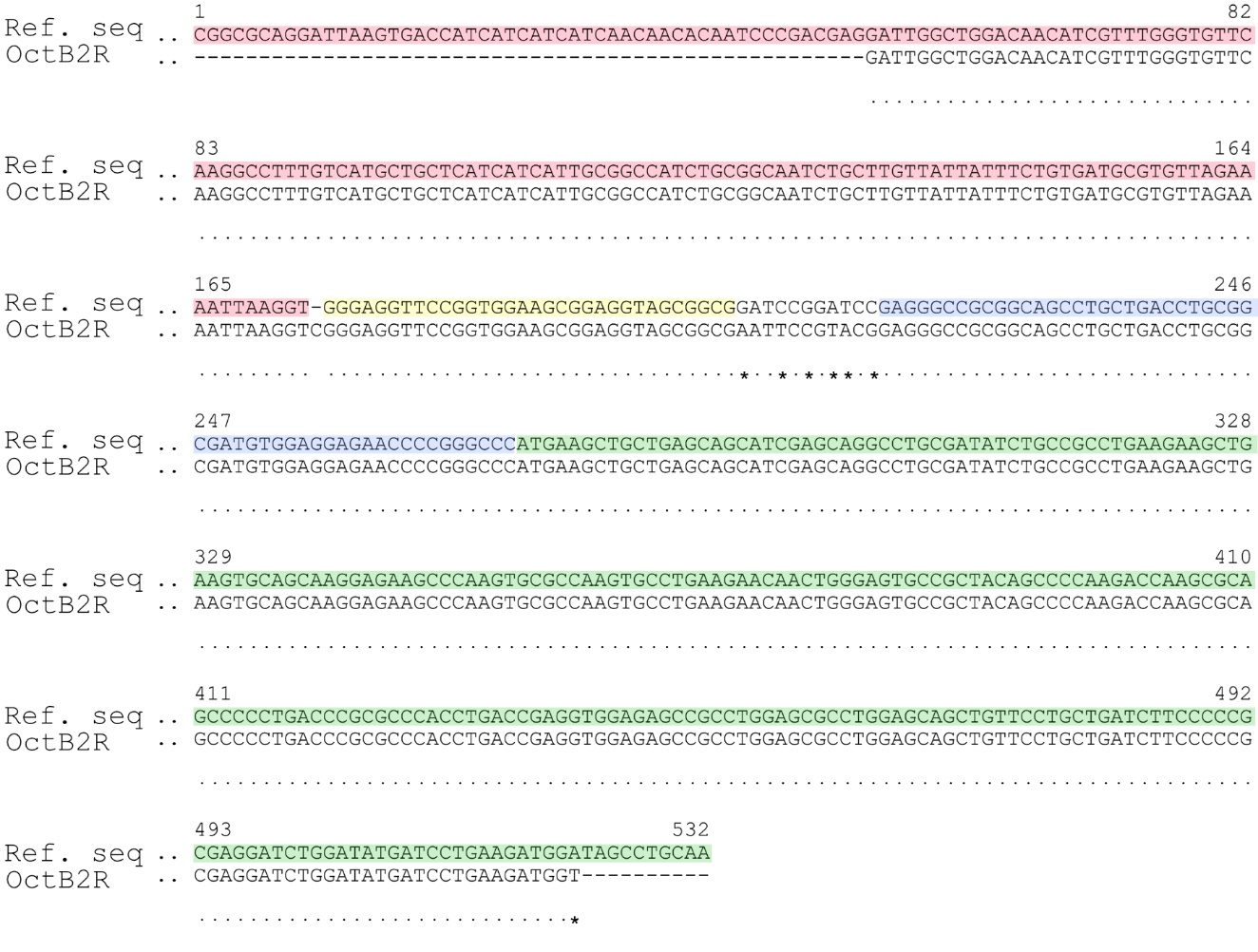

**Supplementary Figure 9: Sequencing result of Trojan *Octβ2R* mRNA to analyze additional regions.** Trojan *Octβ2R* mRNA was analyzed regarding the additional sequences that were found at the genomic level. Here, the sequencing result of the amplified product depicted in Supplementary Figure 8A_I_ is shown. The upper row is the reference sequence (Ref. seq) consisting of Exon 7 (red) that is upstream of the Trojan Exon and the Trojan Exon itself (partially covered with the linker (yellow), T2A (blue) and partial Gal4 (green)). The reference sequence shows only the region that is amplified using the selected primer (not the entire Exon 7) and Trojan Exon. The sequenced amplicon has a size of 470 bp, a bit shorter than the observed product in the gel picture which is due to the sequencing process. The upper row is the reference sequence (Ref. seq), the lower row is the sequenced amplicon of Trojan *Octβ2R* mRNA (OctB2R). Red: Exon 7; yellow: linker (belongs to Trojan Exon); blue: T2A; green: Gal4; Dots show same bases, dashes show missing bases, asterisks show dissimilar bases.
