## Supplementary Table 1 for "Octopamine receptors at a glance: from expression and anatomical maps to their role in development and behavior in the *Drosophila melanogaster* larva"

**Supplementary Table 2: Quantification of homozygous and heterozygous animals in balanced fly lines.** Heterozygous animals in Trojan *Oamb* and *Octβ2R* carried the TM6B, Hu, Tb [1] balancer and were identified using the Tb [1] marker. Heterozygous animals in Trojan *Octα2R* carried the TM3, Sb[1] Ser[1] balancer and were identified using the Sb[1] marker. Table A (Trojan *Oamb*), B (Trojan *Octβ2R*) and C (Trojan *Octα2R*) list the number of homozygous and heterozygous animals as well as the total number of animals. These figures were used to calculate the percentage of homozygous animals.

A: Trojan *Oamb*

| **Homozygote** | **Heterozygote** | **Total** | **Homozygote Animals in %** |
| --- | --- | --- | --- |
| 20 | 94 | 114 | **17.54** |
| 35 | 171 | 206 | **16.99** |
| 24 | 128 | 152 | **15.79** |
| 41 | 278 | 319 | **12.85** |
| 23 | 189 | 212 | **10.85** |
| 28 | 105 | 133 | **21.05** |
| 18 | 141 | 159 | **11.32** |
| 35 | 191 | 226 | **15.49** |
| 21 | 148 | 169 | **12.43** |
| 28 | 223 | 251 | **11.16** |
| 33 | 239 | 272 | **12.13** |
| 18 | 188 | 206 | **8.74** |
| 46 | 417 | 463 | **9.94** |
| 57 | 472 | 529 | **10.78** |
| 40 | 250 | 290 | **13.79** |
| 36 | 305 | 341 | **10.56** |

B: Trojan *Octβ2R*

| **Homozygote** | **Heterozygote** | **Total** | **Homozygote Animals in %** |
| --- | --- | --- | --- |
| 17 | 244 | 261 | **6.51** |
| 15 | 215 | 230 | **6.52** |
| 8 | 156 | 164 | **4.88** |
| 7 | 110 | 117 | **5.98** |
| 15 | 145 | 160 | **9.38** |
| 12 | 161 | 173 | **6.94** |
| 4 | 155 | 159 | **2.52** |
| 17 | 171 | 188 | **9.04** |
| 15 | 172 | 187 | **8.02** |
| 8 | 147 | 155 | **5.16** |
| 19 | 137 | 156 | **12.18** |
| 15 | 171 | 186 | **8.06** |
| 8 | 198 | 206 | **3.88** |
| 6 | 312 | 318 | **1.89** |
| 16 | 306 | 322 | **4.97** |
| 31 | 258 | 289 | **10.73** |

C: Trojan *Octα2R*

| **Homozygote** | **Heterozygote** | **Total** | **Homozygote Animals in %** |
| --- | --- | --- | --- |
| 0 | 64 | 64 | **0.00** |
| 0 | 59 | 59 | **0.00** |
| 0 | 60 | 60 | **0.00** |
| 0 | 94 | 94 | **0.00** |
| 0 | 121 | 121 | **0.00** |
| 0 | 151 | 151 | **0.00** |
| 0 | 58 | 58 | **0.00** |
| 0 | 68 | 68 | **0.00** |
| 0 | 57 | 57 | **0.00** |
| 0 | 63 | 63 | **0.00** |
| 0 | 55 | 55 | **0.00** |
| 0 | 55 | 55 | **0.00** |
| 0 | 110 | 110 | **0.00** |
| 0 | 87 | 87 | **0.00** |
| 0 | 80 | 80 | **0.00** |
| 0 | 141 | 141 | **0.00** |
