## Supplementary Table 4 for "Octopamine receptors at a glance: from expression and anatomical maps to their role in development and behavior in the *Drosophila melanogaster* larva"

**Supplementary Table 5: Domain predictions for Tyramine-β-hydroxylase and octopamine receptors (OAR).** Using the prediction tool “NCBI Conserved Domain Database” all transcripts of each gene of interest were analyzed. The interval indicates the region that is predicted to be a conserved domain. The Expect value (E-value) shows the probability of a sequence of interest to be found in the database. The smaller the E-value the higher the statistical significance of a hit. GPCR = G-protein coupled receptor.

| Gene | Isoform | Predicted domain | Interval | E-value |
| --- | --- | --- | --- | --- |
| Tβh | B | Copper type II ascorbate-dependent monooxygenase, C-terminal domain | 1439-1909 | 1.74e^-78^ |
|  |  | Copper type II ascorbate-dependent monooxygenase, N-terminal domain | 1007-1384 | 8.27e^-56^ |
|  |  | DOMON (dopamine beta-monooxygenase N-terminal) | 530- 871 | 3.15e^-28^ |
|  | C | Copper type II ascorbate-dependent monooxygenase, C-terminal domain | 1438-1908 | 1.73e^-78^ |
|  |  | Copper type II ascorbate-dependent monooxygenase, N-terminal domain | 1006-1383 | 8.26e^-56^ |
|  |  | DOMON (dopamine beta-monooxygenase N-terminal) | 529-870 | 3.15e^-28^ |
| Oamb | B | class A family of seven-transmembrane GPCR | 1057-2754 | 2.66e^-114^ |
|  | C | class A family of seven-transmembrane GPCR | 631-2319 | 5.86e^-127^ |
|  | D | class A family of seven-transmembrane GPCR | 1057-2745 | 5.28e^-126^ |
|  | E | class A family of seven-transmembrane GPCR | 1057-2754 | 2.66e^-114^ |
|  | F | class A family of seven-transmembrane GPCR | 1057-2745 | 5.28e^-126^ |
|  | G | class A family of seven-transmembrane GPCR | 1057-2745 | 5.28e^-126^ |
| Octα2R | A | alpha-2 adrenergic receptor, class A family of seven-transmembrane GPCR | 1227-1754 | 2.27e^-97^ |
|  |  | seven-transmembrane GPCR superfamily | 2679-2912 | 1.87e^-38^ |
|  | B | alpha-2 adrenergic receptor, class A family of seven-transmembrane GPCR | 1227-1754 | 2.27e^-97^ |
|  |  | seven-transmembrane GPCR superfamily | 2679-2912 | 1.87e^-38^ |
|  | C | alpha-2 adrenergic receptor, class A family of seven-transmembrane GPCR | 1227-1754 | 1.43e^-97^ |
|  |  | seven-transmembrane GPCR superfamily | 2592-2825 | 1.56e^-38^ |
| Octβ1R | A | beta-adrenergic receptor-like OAR, class A family of seven-transmembrane GPCR | 616-1536 | 1.11e^-172^ |
|  | B | beta-adrenergic receptor-like OAR, class A family of seven-transmembrane GPCR | 616-1536 | 5.08e^-171^ |
|  | C | beta-adrenergic receptor-like OAR, class A family of seven-transmembrane GPCR | 616-1536 | 2.42e^-172^ |
|  | E | beta-adrenergic receptor-like OAR, class A family of seven-transmembrane GPCR | 616-1536 | 1.68e^-163^ |
| Octβ2R | A | beta-adrenergic receptor-like OAR, class A family of seven-transmembrane GPCR | 675-1640 | 1.83e^-170^ |
|  | C | beta-adrenergic receptor-like OAR, class A family of seven-transmembrane GPCR | 1644-2609 | 1.67e^-166^ |
|  | D | beta-adrenergic receptor-like OAR, class A family of seven-transmembrane GPCR | 1478-2443 | 3.95e^-167^ |
|  | E | beta-adrenergic receptor-like OAR, class A family of seven-transmembrane GPCR | 3144-4109 | 4.50e^-159^ |
|  | F | beta-adrenergic receptor-like OAR, class A family of seven-transmembrane GPCR | 675-1640 | 1.83e^-170^ |
|  | G | beta-adrenergic receptor-like OAR, class A family of seven-transmembrane GPCR | 3223-4188 | 5.22e^-159^ |
| Octβ3R | F | seven-transmembrane GPCR superfamily | 1511-2101 | 9.39e^-125^ |
|  |  | seven-transmembrane GPCR superfamily | 4568-4789 | 2.29e^-47^ |
|  | G | seven-transmembrane GPCR superfamily | 1511-2101 | 6.08e^-125^ |
|  |  | seven-transmembrane GPCR superfamily | 4154-4375 | 3.67e^-47^ |
|  | J | beta-adrenergic receptor-like OAR, class A family of seven-transmembrane GPCR | 1511-2356 | 3.19e^-166^ |
|  | K | seven-transmembrane GPCR superfamily | 2284-2505 | 2.26e^-41^ |
|  |  | seven-transmembrane GPCR superfamily | 1511-2101 | 1.47e^-119^ |
| Oct-TyrR | A | tyramine/octopamine receptor-like, class A family of seven-transmembrane GPCR | 999-1553 | 2.75e^-119^ |
|  |  | seven-transmembrane GPCR superfamily | 2226-2453 | 8.49e^-47^ |
|  | B | tyramine/octopamine receptor-like, class A family of seven-transmembrane GPCR | 992-1546 | 2.24e^-119^ |
|  |  | seven-transmembrane GPCR superfamily | 2219-2446 | 7.84e^-47^ |
