## Supplementary Table 7 for "Octopamine receptors at a glance: from expression and anatomical maps to their role in development and behavior in the *Drosophila melanogaster* larva"

**Supplementary Table 1: Overview of expression pattern of Octopamine receptors in the entire body and central nervous system (CNS) of 3^rd^ instar larva.** Summary of earlier published and in the study identified expression locations with the mutant line or method used. Whole body scans of 1^st^ instar larvae show a similar, yet weaker expression pattern (Supplementary Figure 2). VNC: ventral nerve cord.

| **Oamb** | | | | |
| --- | --- | --- | --- | --- |
| **Site** | **Expressed in** | | **Mutant/ Method used** | **Source** |
| Body | Bipolar dendrite neuron | | Mi{Trojan-GAL4.1}Oamb[MI12417-TG4.1] | This study |
|  | Chordotonal organ | | Mi{Trojan-GAL4.1}Oamb[MI12417-TG4.1] | This study |
|  | Dendritic arborization neuron | | Mi{Trojan-GAL4.1}Oamb[MI12417-TG4.1] | Boivin et al., 2025; This study |
|  | Dorsal organ | | Mi{Trojan-GAL4.1}Oamb[MI12417-TG4.1] | This study |
|  | External sense neuron | | Mi{Trojan-GAL4.1}Oamb[MI12417-TG4.1] | This study |
|  | Labial organ | | Mi{Trojan-GAL4.1}Oamb[MI12417-TG4.1] | This study |
|  | Motor neurons | | scRNAseq, OAMB-10xV5 | Jetti et al., 2023; Bakshinska et al., 2025 |
|  |  |  | Mi{Trojan-GAL4.1}Oamb[MI12417-TG4.1] | This study |
|  | Spiracles | | P{Oamb-GAL4.E} | El-Kholy et al., 2015 |
|  | Terminal organ | | Mi{Trojan-GAL4.1}Oamb[MI12417-TG4.1] | This study |
|  | Terminal sensory cone | | Mi{Trojan-GAL4.1}Oamb[MI12417-TG4.1] | This study |
|  | Tracheal system | | P{Oamb-GAL4.E} | El-Kholy et al., 2015 |
|  | Tracheal dendrite neuron | | Mi{Trojan-GAL4.1}Oamb[MI12417-TG4.1] | This study |
|  | Ventral organ | | Mi{Trojan-GAL4.1}Oamb[MI12417-TG4.1] | This study |
| CNS | Antennal lobe | | Mi{Trojan-GAL4.1}Oamb[MI12417-TG4.1] | This study |
|  | Calyx | | Mi{Trojan-GAL4.1}Oamb[MI12417-TG4.1] | This study |
|  | Mushroom Body | | P{Oamb-GAL4.E} | El-Kholy et al., 2015 |
|  |  |  | Mi{Trojan-GAL4.1}Oamb[MI12417-TG4.1] | This study |
|  | Somata in hemispheres and VNC | | P{Oamb-GAL4.E} | El-Kholy et al., 2015 |
|  |  |  | Mi{Trojan-GAL4.1}Oamb[MI12417-TG4.1] | This study |
|  | Subesophageal zone | | Mi{Trojan-GAL4.1}Oamb[MI12417-TG4.1] | This study |
| **Octα2R** | | | | |
| **Site** | **Expressed in** | **Mutant/ Method used** | | **Source** |
| Body | Chordotonal organ | Mi{Trojan-GAL4.0}Octalpha2R[MI10227-TG4.0] | | This study |
|  | Dendritic arborization neuron | Mi{Trojan-GAL4.0}Octalpha2R[MI10227-TG4.0] | | This study |
|  | Dorsal organ ganglion | Mi{Trojan-GAL4.0}Octalpha2R[MI10227-TG4.0] | | This study |
|  | External sense neuron | Mi{Trojan-GAL4.0}Octalpha2R[MI10227-TG4.0] | | This study |
|  | Motor neurons | scRNAseq | | Jetti et al., 2023; Bakshinska et al., 2025 |
|  |  | Mi{Trojan-GAL4.0}Octalpha2R[MI10227-TG4.0] | | This study |
|  | Spiracle sense organ | Mi{Trojan-GAL4.0}Octalpha2R[MI10227-TG4.0] | | This study |
|  | Terminal organ | Mi{Trojan-GAL4.0}Octalpha2R[MI10227-TG4.0] | | This study |
|  | Terminal sensory cone | Mi{Trojan-GAL4.0}Octalpha2R[MI10227-TG4.0] | | This study |
|  | Ventral organ | Mi{Trojan-GAL4.0}Octalpha2R[MI10227-TG4.0] | | This study |
| CNS | Calyx | Mi{Trojan-GAL4.0}Octalpha2R[MI10227-TG4.0] | | This study |
|  | Mushroom Body | Mi{Trojan-GAL4.0}Octalpha2R[MI10227-TG4.0] | | This study |
|  | Somata in hemispheres and VNC | Mi{Trojan-GAL4.0}Octalpha2R[MI10227-TG4.0] | | This study |
|  | Subesophageal zone | Mi{Trojan-GAL4.0}Octalpha2R[MI10227-TG4.0] | | This study |
| **Octβ1R** | | | | |
| **Site** | **Expressed in** | **Mutant/ Method used** | | **Source** |
| Body | Bipolar dendrite neurons | Mi{Trojan-GAL4.2}Octbeta1R[MI05807-TG4.2] | | This study |
|  | Chordotonal organ | Mi{Trojan-GAL4.2}Octbeta1R[MI05807-TG4.2] | | This study |
|  | Dendritic arborization neuron | Mi{Trojan-GAL4.2}Octbeta1R[MI05807-TG4.2] | | This study |
|  | Dorsal organ | Mi{Trojan-GAL4.2}Octbeta1R[MI05807-TG4.2] | | This study |
|  | Eye-antennal imaginal disc | In situ hybridization, P{GMR19H07-GAL4}, P{GMR20C11-GAL4}, P{GMR21E03-GAL4} | | Ohhara et al., 2012; Koon and Budnik, 2012 |
|  | Eye imaginal disc nuclei | Bulk RNA-sequencing analysis | | This study |
|  | External sense neuron | Mi{Trojan-GAL4.2}Octbeta1R[MI05807-TG4.2] | | This study |
|  | Malpighian tubule | P{Octβ1R-GAL4.E} | | El-Kholy et al., 2015 |
|  | Motor neuron, type I, II and III | P{GMR19H07-GAL4}, P{GMR20C11-GAL4}, P{GMR20E11-GAL4}, P{GMR21E03-GAL4} | | Koon and Budnik, 2012 |
|  |  | scRNAseq | | Jetti et al., 2023; Bakshinska et al., 2025 |
|  |  | Mi{Trojan-GAL4.2}Octbeta1R[MI05807-TG4.2] | | This study |
|  | Salivary gland | In situ hybridization | | Ohhara et al., 2012 |
|  | Spiracle sense organ | Mi{Trojan-GAL4.2}Octbeta1R[MI05807-TG4.2] | | This study |
|  | Terminal organ | Mi{Trojan-GAL4.2}Octbeta1R[MI05807-TG4.2] | | This study |
|  | Terminal sensory cone | Mi{Trojan-GAL4.2}Octbeta1R[MI05807-TG4.2] | | This study |
|  | Tracheal system | P{Octβ1R-GAL4.E} | | El-Kholy et al., 2015 |
|  | Ventral organ | Mi{Trojan-GAL4.2}Octbeta1R[MI05807-TG4.2] | | This study |
| CNS | Antennal lobe | Mi{Trojan-GAL4.2}Octbeta1R[MI05807-TG4.2] | | This study |
|  | Calyx | Mi{Trojan-GAL4.2}Octbeta1R[MI05807-TG4.2] | | This study |
|  | Mushroom Body | Mi{Trojan-GAL4.2}Octbeta1R[MI05807-TG4.2] | | This study |
|  | Somata in hemispheres and VNC | P{GMR19H07-GAL4}, P{GMR20C11-GAL4}, P{GMR20E11-GAL4}, P{GMR21E03-GAL4} | | Koon and Budnik, 2012 |
|  |  | P{Octβ1R-GAL4.E} | | El-Kholy et al., 2015 |
|  |  | In situ hybridization | | Ohhara et al., 2012 |
|  |  | Mi{Trojan-GAL4.2}Octbeta1R[MI05807-TG4.2] | | This study |
|  | Subesophageal zone | Mi{Trojan-GAL4.2}Octbeta1R[MI05807-TG4.2] | | This study |
| **Octβ2R** | | | | |
| **Site** | **Expressed in** | **Mutant/ Method used** | | **Source** |
| Body | Fat body | RT-PCR | | El-Kholy et al., 2015 |
|  | Imaginal disc | In situ hybridization | | Ohhara et al., 2012 |
|  | Intestine | RT-PCR | | El-Kholy et al., 2015 |
|  | Malpighian tubule | RT-PCR | | El-Kholy et al., 2015 |
|  | Midgut | In situ hybridization | | Ohhara et al., 2012 |
|  | Motor neuron, type I and II | PBac{WH}Octß2R[f05679], P{GD2954}, P{KK109808}; scRNAseq | | Koon et al., 2011; Koon and Budnik, 2012; Jetti et al., 2023 |
|  | Muscles | RT-PCR | | Koon et al., 2011 |
|  |  | P{Octβ2R-GAL4.L} | | El-Kholy et al., 2015 |
|  |  | Mi{Trojan-GAL4.2}Octbeta2R[MI13416-TG4.2]; Bulk RNA-sequencing analysis | | This study |
|  | Salivary gland | RT-PCR | | El-Kholy et al., 2015 |
|  |  | In situ hybridization | | Ohhara et al., 2012 |
|  | Terminal organ | Mi{Trojan-GAL4.2}Octbeta2R[MI13416-TG4.2] | | This study |
|  | Tracheal system | RT-PCR | | El-Kholy et al., 2015 |
| CNS | Somata in hemispheres and VNC | In situ hybridization | | Ohhara et al., 2012 |
|  |  | P{Octβ2R-GAL4.L}, RT-PCR | | El-Kholy et al., 2015 |
|  |  | Mi{Trojan-GAL4.2}Octbeta2R[MI13416-TG4.2] | | This study |
|  | Subesophageal zone | Mi{Trojan-GAL4.2}Octbeta2R[MI13416-TG4.2] | | This study |
| **Octβ3R** | | | | |
| **Site** | **Expressed in** | **Mutant/ Method used** | | **Source** |
| Body | Bipolar dendrite neuron | Mi{Trojan-GAL4.un}Octbeta3R[MI06217-TG4.un] | | This study |
|  | Chordotonal organ | Mi{Trojan-GAL4.un}Octbeta3R[MI06217-TG4.un] | | This study |
|  | Dorsal organ | Mi{Trojan-GAL4.un}Octbeta3R[MI06217-TG4.un] | | This study |
|  | Dendritic arborization neuron | Mi{Trojan-GAL4.un}Octbeta3R[MI06217-TG4.un] | | This study |
|  | External sense neuron | Mi{Trojan-GAL4.un}Octbeta3R[MI06217-TG4.un] | | This study |
|  | Eye imaginal disc nuclei | Bulk RNA-sequencing analysis | | This study |
|  | Imaginal disc | In situ hybridization | | Ohhara et al., 2012 |
|  | Malpighian tubule | RT-PCR | | El-Kholy et al., 2015 |
|  | Midgut | In situ hybridization | | Ohhara et al., 2012 |
|  | Motor neuron | scRNAseq | | Jetti et al., 2023 |
|  |  | Mi{Trojan-GAL4.un}Octbeta3R[MI06217-TG4.un] | | This study |
|  | Posterior located sensilla on body-wall | P{Octβ3R-GAL4.E} | | El-Kholy et al., 2015 |
|  | Prothoracic gland | In situ hybridization | | Ohhara et al., 2012 |
|  | Reproductive organ | In situ hybridization | | Ohhara et al., 2012 |
|  | Salivary gland | In situ hybridization | | Ohhara et al., 2012 |
|  | Spiracle sense organ | Mi{Trojan-GAL4.un}Octbeta3R[MI06217-TG4.un] | | This study |
|  | Terminal organ | Mi{Trojan-GAL4.un}Octbeta3R[MI06217-TG4.un] | | This study |
|  | Terminal sensory cone | Mi{Trojan-GAL4.un}Octbeta3R[MI06217-TG4.un] | | This study |
| CNS | Antennal lobe | Mi{Trojan-GAL4.un}Octbeta3R[MI06217-TG4.un] | | This study |
|  | Calyx | Mi{Trojan-GAL4.un}Octbeta3R[MI06217-TG4.un] | | This study |
|  | Mushroom Body | P{Octβ3R-GAL4.E} | | El-Kholy et al., 2015 |
|  |  | Mi{Trojan-GAL4.un}Octbeta3R[MI06217-TG4.un] | | This study |
|  | Somata in hemispheres and VNC | P{Octβ3R-GAL4.E} | | El-Kholy et al., 2015 |
|  |  | Mi{Trojan-GAL4.un}Octbeta3R[MI06217-TG4.un] | | This study |
|  | Subesophageal zone | Mi{Trojan-GAL4.un}Octbeta3R[MI06217-TG4.un] | | This study |
| **Oct-TyrR** | | | | |
| **Site** | **Expressed in** | **Mutant/ Method used** | | **Source** |
| Body | Aorta | P{Oct-TyrR-GAL4.E} | | El-Kholy et al., 2015 |
|  | Chordotonal organ | Mi{Trojan-GAL4.un}Oct-TyrR[MI03485-TG4.un] | | This study |
|  | Dendritic arborization neuron | Mi{Trojan-GAL4.un}Oct-TyrR[MI03485-TG4.un] | | This study |
|  | Dorsal multidendritic neuron 1 | Mi{Trojan-GAL4.un}Oct-TyrR[MI03485-TG4.un] | | This study |
|  | External sense neuron | Mi{Trojan-GAL4.un}Oct-TyrR[MI03485-TG4.un] | | This study |
|  | Lymph gland | Bulk RNA-sequencing analysis | | This study |
|  | Malpighian tubule | RT-PCR | | El-Kholy et al., 2015 |
|  | Motor neuron | PBac{Oct-TyrR-GAL4.S} | | Stowers, 2011 |
|  |  | scRNAseq | | Jetti et al., 2023 |
|  |  | Mi{Trojan-GAL4.un}Oct-TyrR[MI03485-TG4.un] | | This study |
|  | Peripheral nervous system | P{Oct-TyrR-GAL4.E} | | El-Kholy et al., 2015 |
|  | Salivary gland | P{Oct-TyrR-GAL4.E} | | El-Kholy et al., 2015 |
|  | Spiracles (posterior) | P{Oct-TyrR-GAL4.E} | | El-Kholy et al., 2015 |
|  | Terminal sensory cone | Mi{Trojan-GAL4.un}Oct-TyrR[MI03485-TG4.un] | | This study |
|  | Tracheal system | P{Oct-TyrR-GAL4.E} | | El-Kholy et al., 2015 |
| CNS | Somata in the hemispheres | P{Oct-TyrR-GAL4.E} | | El-Kholy et al., 2015 |
|  |  | Mi{Trojan-GAL4.un}Oct-TyrR[MI03485-TG4.un] | | This study |
|  | Somata in the VNC | PBac{Oct-TyrR-GAL4.S} | | Stowers, 2011 |
|  |  | Mi{Trojan-GAL4.un}Oct-TyrR[MI03485-TG4.un] | | This study |
|  | Subesophageal zone | Mi{Trojan-GAL4.un}Oct-TyrR[MI03485-TG4.un] | | This study |
